# Phosphorylation Mimicking Mutations Cause TDP-43 to Adopt Different Fibril Conformations

**DOI:** 10.64898/2026.05.14.725298

**Authors:** Blake D. Fonda, Dylan T. Murray

**Affiliations:** Department of Chemistry, University of California, Davis, 95616, United States of America; Department of Molecular and Cell Biology, University of Connecticut, Storrs, Connecticut, 06269, United States of America

## Abstract

The Tar-DNA Binding Protein-43 C-terminal region, TDP43LC, has been previously shown to form amyloid-like fibrils with distinct folds in ALS and FTD. In both diseases, proteinaceous inclusions contain TDP43 C-terminal protein fragments as well as phosphorylated TDP43. Here, we use solution NMR to show that soluble phosphomimetic TDP43LC, P-TDP43LC, is structurally similar to wild-type TDP43LC. Disperse P-TDP43LC, like wild-type protein, contains a central helical region flanked by long disordered regions. Despite this similarity, our turbidity measurements, imaging, and kinetic assays show that P-TDP43LC has different aggregation behavior than wild-type protein. Using solid state NMR measurements we find that that phosphomimetic mutations alter the wild-type fibril conformation. Electrostatic repulsion from negatively charged sidechains, despite having little effect on the soluble protein’s structure, perturbs amyloid-like fibril formation and selects for a different conformation in vitro. These results shed light on the structural role of TDP43LC phosphorylation in fibril formation in disease.

**TOC Graphic:** 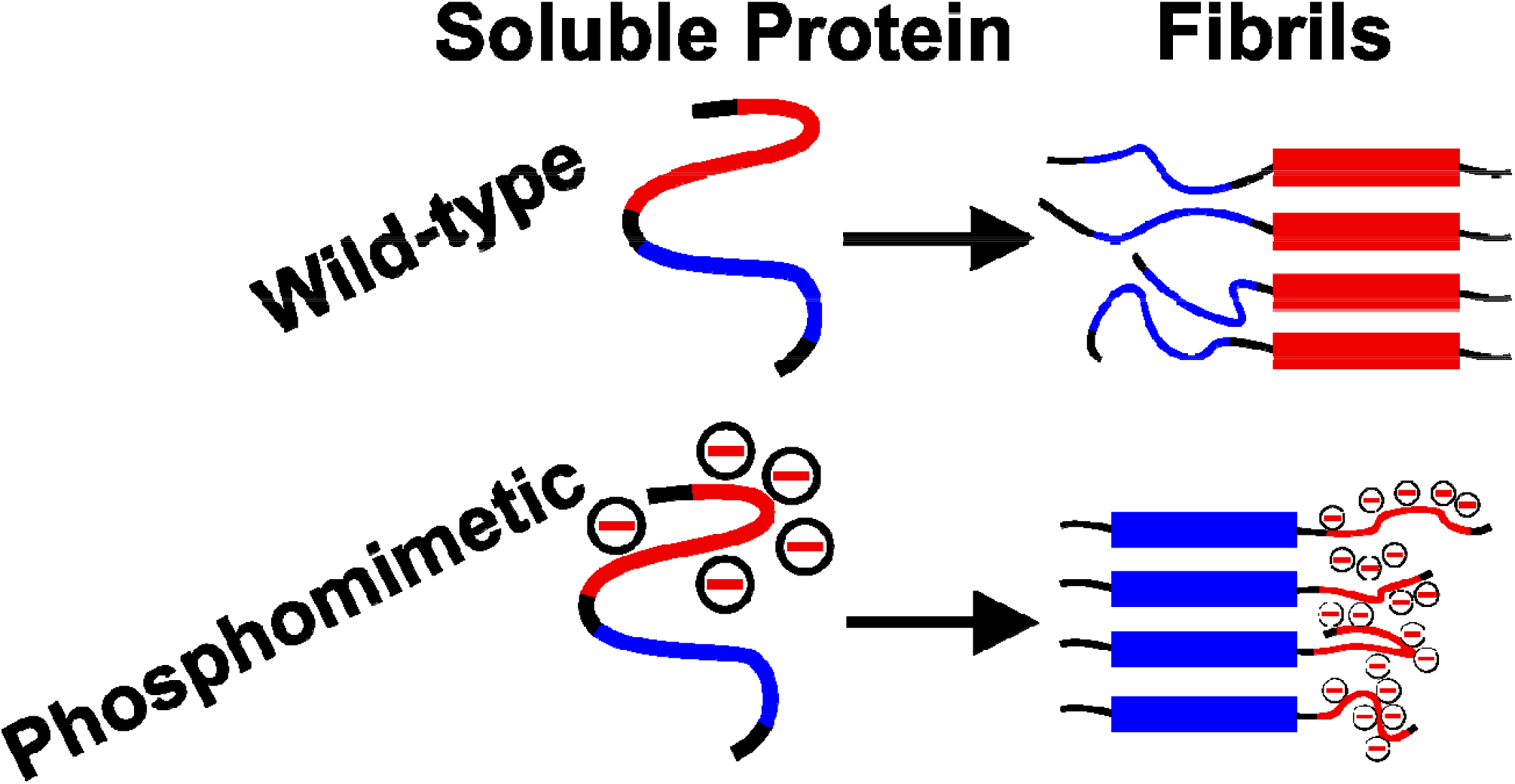

**Synopsis:** Phosphomimetic mutations at ALS and FTD neurodegeneration-associated sites in an amyloid forming protein perturbs the aggregated structure compared to wild-type protein.

## Introduction

Tar-DNA Binding Protein-43, TDP43, is a human transcription factor protein with a RNA splicing function activity. One of the essential targets of TDP43 splicing activity is the protein cystic transmembrane conductance regulator, an essential transporter of anions for human cells that is important in proper cardiac health^1^. TDP43 misfunction is observed in a myriad of diseases including ALS and Frontal-Temporal Dementia (FTD). Ex vivo tissue samples from multiple individuals with these neurodegenerative diseases show TDP43, and in particular its C-terminal domain, forms amyloid-like fibrils, rich in repetitive cross-β fibril structure^2^. High resolution cryogenic electron microscopy (cryo-EM) structural studies have recently identified unique fibril conformations in brain tissue from ALS and FTLD type-A patients, suggesting that the molecular structure of TDP43 fibrils might be indicative of specific neurodegenerative disease pathology^3,4^.

The cryo-EM studies determined at high resolution the core region that stabilizes each fibril structure, yet necessary proteolytic processing during sample preparation^3,4^ led to cleavage of a significant portion of the C-terminal region of the TDP43LC^3,4^. The regions flanking the disease-associated fibril core were not structurally characterized and therefore remain to be investigated.

The proteinaceous inclusions from tissue samples show TDP43 undergoing post-translational modifications^5,6^. Hyperphosphorylation occurs across the C-terminal region of TDP43 and includes sites *outside* of the fibril core region associated with diease^3,4,7^. No phosphorylation events were observed within the core fibril-forming regions determined from the ALS and FTLD type-A. However, phosphorylation of residues outside of the disease-associated fibril core might be influencing TDP43 aggregation behavior^5,6^. In addition, the patterning of phosphorylation in the TDP43LC C-terminal domain is disease dependent^8^, which could influence conformational selection and explain the different ALS and FTLD-type A structures. The molecular mechanism for how disease-specific phosphorylation patterns select TDP43 fibril conformations remain uncharacterized.

The behavior of phosphorylated TDP43 and corresponding TDP43 phosphomimetic mutants, was recently investigated using both purified proteins and cellular studies^9,10^. Phosphorylation appears to prolong the aggregation process of TDP43, with little impact on TDP43 function^9^. Phosphomimetic TDP43 is able to functionally shuttle back and forth between the nucleus and the cytoplasm. However, these mutations reduce TDP43’s ability to phase separate into membrane-less organelles in cultured cells^9^.

Our previous in vitro structural study of TDP43LC fibrils showed, alongside the work many others, that different regions of the TDP43 C-terminal region can adopt fibril conformations in vitro^11–20^. Based on the TDP43LC phosphorylation sites residing outside of the disease-associated fibril core, we hypothesize that electrostatic repulsion drives the TDP43 fibril conformation. To investigate this question, we analyzed phosphomimetic TDP43LC using solution NMR, turbidity measurements, imaging, kinetic assays, and solid state NMR.

## Results

Using TDP43 phosphorylation patters in pathological proteinaceous inclusions from patient tissues^5,7^ as a guide, we mutated five Ser residues to Asp residues. These mutations do not perfectly mimic phosphorylated Ser, but they do introduce negative charges at the disease-identified phosphorylated sites with complete specificity and therefore provide a model system probe the effect electrostatics have on perturbing TDP43 ds in solution, its aggregation kinetics, and its fibril structure. Supplementary Figure 1 show the amino acid sequence of the TDP43LC, illustrating its biased amino acids composition, enriched in Gly, Ser, Asn, and Ala residues.

To investigate the differences between P-TDP43LC and the wild-type TDP43LC using NMR, we generated protein with an N-terminal affinity tag using recombinant expression in E. coli with ^13^C and ^15^N isotopically labeled media. The protein was purified in denaturing conditions with nickel-affinity chromatography and then transferred to non-denaturing conditions. The purification tag was removed by treatment with TEV protease, and reverse nickel-affinity chromatography was used to further purify TDP43LC. This generated relatively pure protein as indicated by SDS PAGE in Supplementary Figure 1.

We first sought to study the solution behavior of P-TDP43LC as compared to wild-type protein. P-TDP43LC was prepared for solution NMR analysis in non-denaturing buffer. A suite of standard 3D heteronuclear NMR experiments were used to obtain NMR chemical shifts, and the sequence-specific assignments were made manually. The assignments are shown in Figure 2A and are tabulated in Supplementary Table 1. The few peaks remaining unassigned are tabulated in Supplementary Table 2. A few of the unassigned peaks correspond to ambiguities in assignments for the ^1^H and ^15^N backbone, but unambiguous ^13^C shifts (G380, G411, and D379). Excluding proline residues, 95% of the ^1^H and ^15^N backbone NMR chemical shifts were assigned to unique residues in the protein sequence. For all non-glycyl residues except R268, at least one ^13^C amino acid side-chain chemical shift was assigned.

**Figure 1.**
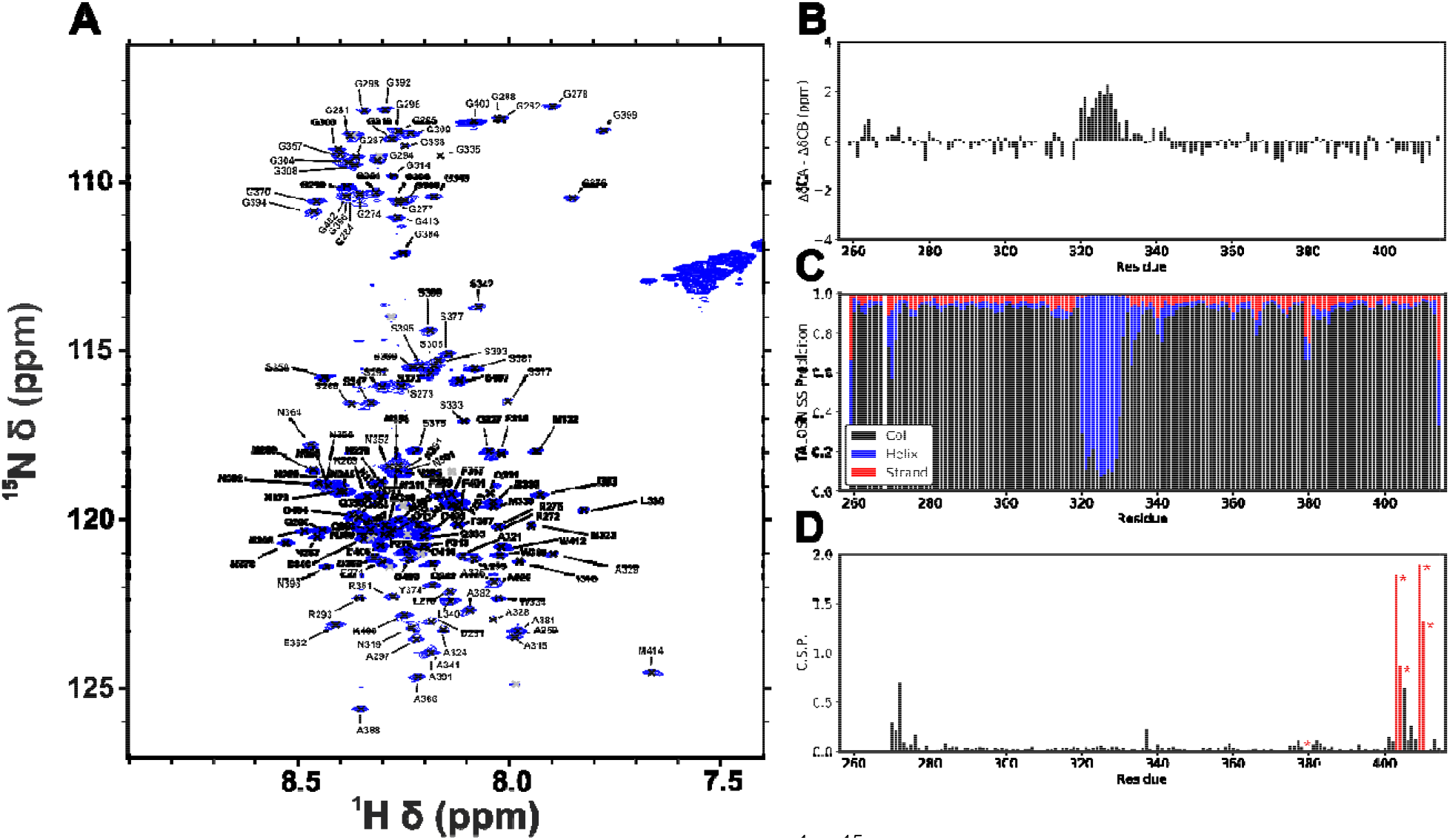
Solution NMR of P-TDP43LC. A) P-TDP43LC ^1^H-^15^N fHSQC with assignments from 3D experiments mapped on the spectrum. Grey peaks represent unassigned peaks, see Supplementary Tables 1–2. B) Secondary chemical shifts relative to the Poulsen IDP calculator for the assigned shifts^21,22^. C) TALOS-N secondary structure predictions based on the assignments^23,24^. D) Chemical shift perturbations relative to assignments from BMRB 26823 depict little perturbation^25,26^. Red sites with an asterisk symbol, *, indicate the five sites with the mutation from Ser to Asp.

**Figure 2.**
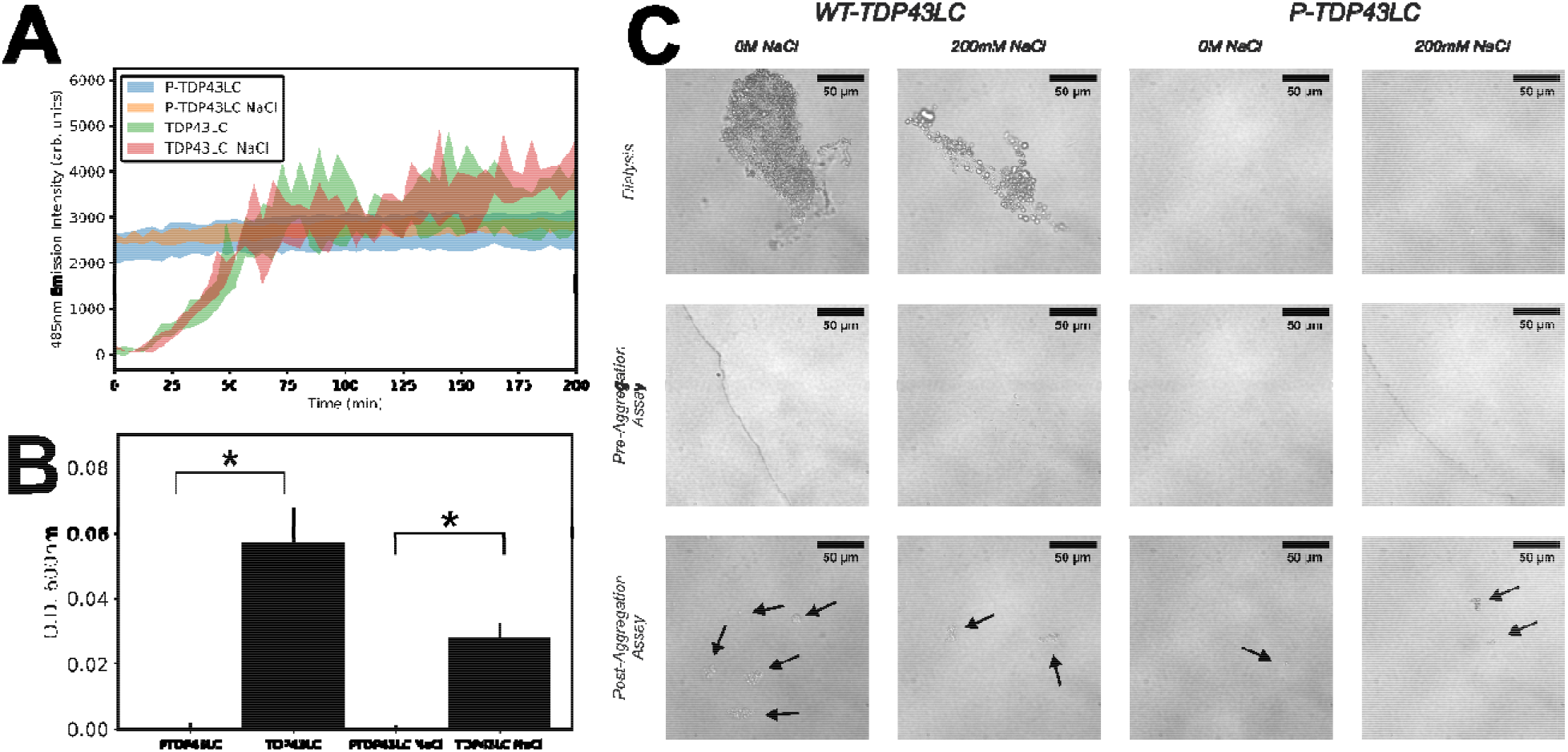
Analysis of P-TDP43LC Aggregation. A) ThT assay fluorescence data with 4.3 M protein, 20 mM sodium phosphate pH 7.4 and 13 M ThT; NaCl conditions also contain 67 mM sodium chloride. The shaded region represents the span of the data from three separate sample wells. B) Turbidity of protein solutions after aggregation according to identical conditions as A) and C) except without any ThT in the buffer. The error bars are the standard deviation across n = 3. The asterisk denotes a significant difference from a independent two-tailed T test at the 95% confidence interval. C) Bright-field microscopy images. Dialysis is after overnight incubation of 35 M protein dialyzed against non-denaturing buffer. Pre-Aggregation Assay is after centrifugation to remove preformed aggregates, followed by dilution to 13 M protein. Post-aggregation assay images are from each solution after the completion of the ThT assay in A) at 23 hrs. The arrows point to protein aggregates in the image and are meant to qualitatively highlight the aggregates in the image.

Figure 1B shows the secondary chemical shifts for P-TDP43LC, which refers to the difference between the expected disordered random coil NMR chemical shifts for each residue and the experimental NMR chemical shift assignment. Regions of residues with positive secondary chemical shifts indicate -helical structure, while regions of negative values indicate -strand structure^21,22,27^. The only secondary structure motif observed is a helical domain from residues P320–L330. Figure 2C shows TALOS-N secondary structure predictions derived from a neural network analysis of databases of proteins with known structure, sequence, and chemical shifts which predicts the residue-specific secondary structure^23,24^. The TALOS-N predictions match the secondary chemical shift result, identifying a helical region from P320-L330. This helical region additionally has reduced S/N compared to the rest of the signals in the ^1^H-^15^N HSQC spectrum. As shown in Supplementary Figure 2, the reduced signal intensity agrees with the helical region retaining slower reorientational motion than the C-terminal flanking region of the TDP43LC. The protein sequences outside the helical region are identified as long disordered regions, i.e. regions without well-defined secondary structure, by both TALOS-N and secondary chemical shift analysis.s

A previous study yielded NMR assignments on wild-type TDP43LC under similar conditions^25^ (BMRB), which we compared with our NMR chemical shift assignments for P-TDP43LC. Figure 1D shows the chemical shift perturbation plot, CSP, which quantifies the differences in ^15^N and ^1^H chemical shifts between P-TDP43LC and wild-type protein^26^. Large CSPs indicate different protein structure. Only two regions with signficant CSPs were observed. The largest differences are at the C-terminal region of P-TDP43LC, where the Ser to Asp mutation sites reside. Residues proximal to these mutation sites have larger CSPs compared to the rest of the sequence. Note the exceptionally large CSPs (>0.5) at the C-terminal region of P-TDP43LC are due to Asp mutations and not the introduction of secondary structure. The other region of P-TDP43LC with slightly elevated CSPs is the N-terminal region, which is likely due the extra residues (262–267) in our construct relative to the reference data (BMRB 26823, residues 267–414)^25^. Overall, the solution NMR analysis indicates that soluble P-TDP43LC retains largely the same structural motifs as wild-type TDP43LC, with the expected central helical region and flanking disordered domains.

Gradient-based diffusion NMR measurements measure a protein’s radius of hydration, which is indicative of oligomeric state and can be used to approximate the relative expansion of the protein chain, i.e. the degree of “foldedness”^28,29^. Supplementary Figures 3–5 show the raw and fitted diffusion NMR data, giving a radius of hydration (Rh) for soluble P-TDP43LC of 22 ± 5 Å and soluble WT-TDP43LC to be 24 ± 1 Å. Using analytical models that relate amino acid residue count to molecular size^28^, if the 156 amino acid residues in our TDP43LC constructs were completely folded the expected Rh would be ∼20.5 Å, while a completely unfolded TDP43LC molecule would have an Rh of ∼39.3 Å. Thus, within our experimental uncertainty, the WT- and P-TDP43LC Rh are indistinguishable, and the Rh value suggests each protein is a partially compacted monomer our solution NMR conditions (16 µM protein, pH 6.2, low salt concentration). Thus, the helix of P-TDP43LC and WT-TDP43LC largely does not initiate dimer formation at the low concentrations we tested, consistent with a previous WT-TDP43LC study^30^.

Next, we sought to analyze the aggregation kinetics of P-TDP43LC versus wild-type TDP43LC. Each protein was placed into identical buffer and pre-formed aggregates were removed prior to the assay via centrifugation. The dye Thioflavin-T, ThT, was added to each protein sample well. ThT is used as an indicator of amyloid formation, and has low unbound fluorescence yet gains intense fluorescence upon binding to β-sheet rich protein structures^31^. Figure 2A shows the ThT assay, with wild-type TDP43LC showing the expected lag-phase in fibril formation preceding a brief exponential growth phase, followed by a plateauing of the ThT signal^25,31–33^. Ionic strength changes revealed little difference in ThT assay signal, likely due to the sodium chloride concentration not being elevated to a high enough concentration, coupled with the 37 °C incubation and 30s of rapid linear shaking conditions used here to aggregate TDP43^31–33^.

The P-TDP43LC shows initially high ThT fluorescence, which was unexpected since turbidity measurements in Figure 2B demonstrate less aggregation or condensate formation for P-TDP43LC than wild-type protein. To our knowledge, ThT measurements have not otherwise been recorded on this P-TDP43LC construct. Turbidity measurements are consistent with the literature for in vitro full-length phosphomimic TDP43, which has been shown to have a reduced but not eliminated propensity to aggregate or condensate relative to WT-TDP43^9^. In addition, bright-field microscopy images shown in Figure 3C qualitatively indicate more aggregated protein for wild-type TDP43LC at the end of the ThT assay, but a lesser amount of aggregated material for P-TDP43LC. We hypothesize based on the ThT, turbidity, and microscopy results that P-TDP43LC is interacting with ThT before amyloid formation, likely due to the introduction of the two Asp-Asp motifs from the phosphomimetic mutations, which could theoretically bind positively charged ThT. It is unlikely that P-TDP43LC is forming a highly-populated fibril or proto-fibril state at the initial time points in these kinetic assays, since these ThT assay conditions were similar to the solution NMR conditions that showed non-strand structure and a radius of hydration consistent with a monomer. These results point to a lack of preformed fibrils in the kinetic assays, however we cannot entirely rule out if the initial ThT fluorescence might be picking up on minorly-populated preformed aggregated material in the P-TDP43LC samples that otherwise was unobservable by NMR, turbidity, and bright field microscopy data.

**Figure 3.**
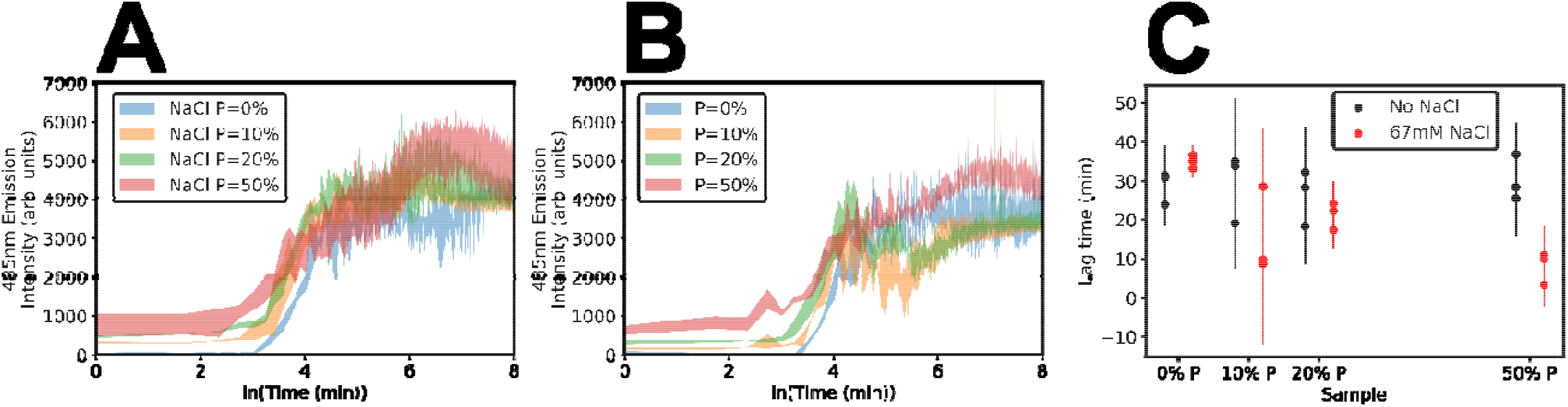
ThT Assay with Mixed Wild-type and P-TDP43LC. A) Logarithmic plot of time versus ThT fluorescence for the specified percent P-TDP43LC with 4 M protein in total, with the shaded region corresponding to the span of the data from n=3 plate reader wells. B) The same as A) except without added salt. C) t_lag_ represents the initiation of the exponential growth in fluorescence intensity in A) and B). The error bars represent the 95% confidence interval for n=3 samples.

Since our first attempt to directly measure the ThT aggregation curves for P-TDP43LC showed uncharacteristic initial ThT fluorescence, we turned towards mixed samples containing both P-TDP43LC and wild-type TDP43LC to study the phosphomimetic influence on TDP43LC aggregation kinetics. Figure 3 shows the expected sigmoidal shape of aggregation kinetics for all samples, however there were no significant changes in any of the aggregation kinetic parameters displayed in Figure 3C and Supplementary Figure 6. The highly similar aggregation lag time for 0% to 50% P-TDP43LC is consistent with the rate limiting step for all samples being the fibril nucleation event for wild-type TDP43LC (i.e. the initial formation of a amyloid-like fibril structure). These data indicate a lack of significant difference for k_app_, the rate at which aggregation occurs after the lag time. t_1/2_, the time till half the maximum fluorescence intensity, was also unperturbed by increasing percentages of P-TDP43LC. Therefore, mixed kinetic assay data indicates that P-TDP43LC does not interrupt, “chaperone”, or seed the aggregation of wild-type protein in mixed samples, yet by itself, P-TDP43LC aggregates more slowly in the absence of wild-type TDP43 as shown by the turbidity measurement, Figure 2B.

Liquid-liquid phase separation, LLPS, was sometimes but not always observed during otherwise identical buffer exchange process using spin filters. Figure 4A shows typical LLPS observed in our WT and P-TDP43LC samples and Supplemental Figure 7 shows LLPs for P-TDP43LC during dialysis into the solution NMR condition, indicating that the phosphomimetic mutations do not inhibit LLPS formation. However, sometimes LLPS was not observed, likely due to the fine-tuned balance between LLPS and the protein aggregation process. During some preparations at high concentrations of P-TDP43LC, aggregation likely occurred too rapidly for LLPS observation, but the lack of LLPS occurred sporadically and unreliably from preparation to preparation. At dilute concentrations of P-TDP43LC, 4 µM in Figure 2C, no LLPS or aggregate formation was ever observed.

**Figure 4.**
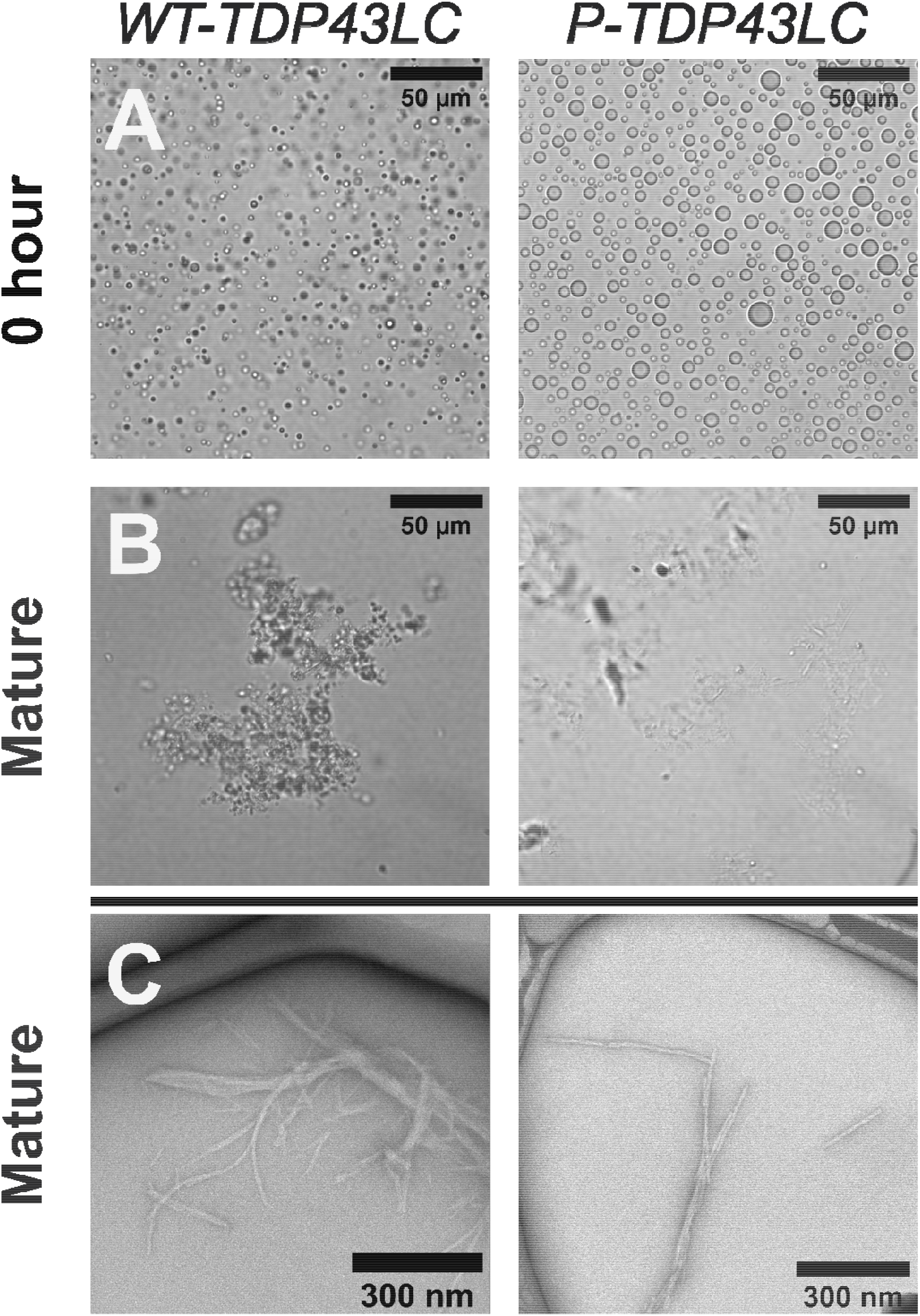
Characterization of Fibrilization after Removal of Denaturant. Comparison of the aggregation and fibrilization process between WT-TDP43LC and P-TDP43LC. A) and B) Bright-field microscopy during the flash dilution buffer exchange procedure showed a phase separation followed by the formation of large insoluble aggregates. C) Negative stain transmission electron microscopy of protein fibrils after the flash dilution protocol. “Mature” is after completion of the flash dilution and fibril incubation protocol.

To initiate TDP43LC fibril formation, we used a flash-dilution protocol, which involves a rapid buffer exchange using centrifuge filters. This procedure replaced denaturing solution with pH 7.4 phosphate buffered solution with 200mM NaCl, followed by a 7-day room temperature incubation with 300 rpm shaking, brief sonification, and then another ∼7 day incubation period^11^. Bright field microscopy images in Figure 4 show sporadic LLPS at intermediate dilution steps of P-TDP43 and WT-TDP43LC, Figure 4, although sporadically as previously discussed. After complete buffer exchange, phase separation was replaced by non-dynamic aggregate formation, see Supplementary Figure 8. At the conclusion of the flash dilution procedure after the removal of denaturant to less than 300 mM urea, with less than 5% of the initial denaturing buffer remaining, bright-field microscopy images in Figure 4B showed the rapid formation of large aggregates. Eventually, both WT and P-TDP43LC proteins form fibrils, as observed by transmission electron microscopy in Figure 4C and bright field microscopy in Supplementary Figure 8. The result was striking, as it demonstrates phosphomimetic mutations do not abolish the formation of amyloid-like fibrils despite five additional negative charges per monomer.

To characterize the fibrils at the level of individual resides, solid state NMR experiments were used. Figure 5A–B shows ^13^C -^13^C refocused-INEPT-TOBSY data for wild-type and P-TDP43LC. This experiment gives rise to signals only for atoms with significant reorientational motions with very short correlation times, i.e. from residues not participating in rigid structure. There are subtle differences in the spectrum for each protein. The shared and unique spectral peaks are tabulated in Supplementary Table 3. The TOBSY spectra is consistent with the flanking regions of both wild-type and phosphomimetic fibrils retaining some highly mobile residues, however the precise composition of these regions are different between the two samples. Due to the length and repetitive nature of the amino acid residue types in the TDP43LC sequence, it is impossible to conclusively pinpoint which residues are highly mobile each sample type. However, the lack of an Asp peak in the wild-type spectra signifies that the two Asp residues in the wild-type construct, the Asp residue leftover from the TEV-cleavage site as well as D406, are not dynamic in wild-type fibrils. Conversely, at least some of the Asp residues in P-TDP43LC remain highly mobile, consistent with either the very N-terminus or the C-terminal phosphomimetic region of P-TDP43LC retaining significant and random reorientational motion. Indeed, in the TOBSY spectra in Figure 5B, the Asp residue shifts of CA = 54.3 ppm and CB = 40.9 ppm is a near match to the ^13^C *solution* NMR assigned shift values for each Asp residue site except for the assigned N-terminal Asp (see Supplementary Table 1). Therefore, these data are consistent with at least some of the phosphomimetic sites in P-TDP43LC *fibrils* retaining highly mobile, disordered structure with rapid motion outside the fibril core region much like how these residues are disordered in *soluble* P-TDP43 (see Figure 1).

**Figure 5.**
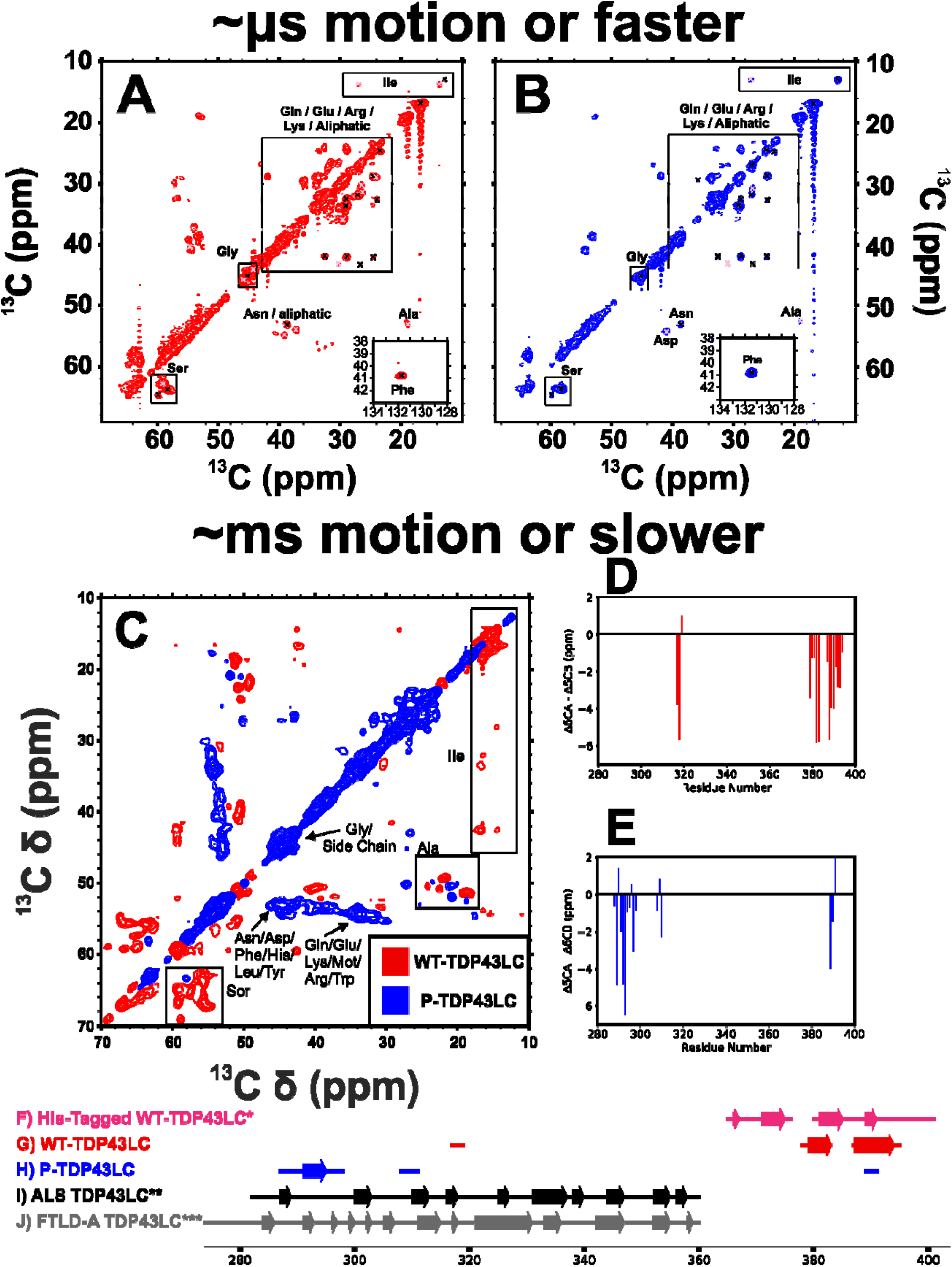
Solid State NMR Spectra Shows P-TDP43LC forms a Different Fibril Conformation than Wild-Type TDP43LC. A) Solid-state NMR refocused-INEPT-TOBSY spectra on wild-type TDP43LC fibrils. Black peak x-marks indicate peaks found in both A) and B) whereas pink x-marks depict peaks unique to wild-type fibrils. B) Solid-state NMR INEPT-TOBSY spectra of P-TDP43LC fibrils. Black peak x-marks are found in both A) and B), where pink x-marks depict peaks unique to P-TDP43LC fibrils. C) Cross polarization based ^13^C-^13^C dipolar-assisted rotational resonance spectra with 50 ms mixing difference spectra comparing WT- and P-TDP43LC. D) cross polarization CA-CB secondary chemical shifts for WT-TDP43LC relative to random coil chemical shifts. E) cross polarization CA-CB secondary chemical shifts for P-TDP43LC fibrils relative to the random coil chemical shifts. F) Arrows depict β-strand structure, and the solid line represents rigid non-β strand residues in the fibrils, taken from the analysis of the His-tagged TDP43LC fibrils^34^. G) Secondary structure for WT-TDP43LC fibrils determined in this work.H) Secondary structure for P-TDP43LC fibrils determined in this work. I) Secondary structure from the cryo-EM reconstruction for ex vivo TDP43 fibrils from ALS^3^. J) Secondary structure from the cryo-EM reconstruction for ex vivo TDP43 fibrils from FTLD-type A^4^.

The solid-state NMR ^1^H-^13^C refocused-INEPT, rINEPT, spectra in Supplementary Figure 9, like the INEPT-TOBSY spectra, also reports on highly mobile residues and corroborates the TOBSY result. A unique Asp residue is seen in the P-TDP43LC spectra, and unique Ile side chain peaks are identified for the P-TDP43LC spectra. The wild-type protein rINEPT spectra contains additional Ser signals, as well as multiple CA peaks not found in the P-TDP43LC spectra, like the TOBSY data also indicates. Overall, the INEPT-TOBSY and rINEPT experiments are consistent with different mobile regions for P-TDP43LC versus wild-type protein fibrils, with the Asp peak identified in these spectra indicating that the region containing the phosphomimetic sites in P-TDP43LC is likely, at least in-part, disordered as a flanking region to the fibril core.

To probe the rigid portion of the fibrils, we used cross-polarization based solid state NMR relaxation experiments. Amino acid residues with significant motion do not appear in these spectra since the cross-polarization based magnetization signal transfers are only efficient for relatively rigid residues with motion, i.e. correlation times, of approximately milliseconds or longer. ^1^H T^1^ρ and 15N T2 relaxation constants were measured for both wild-type and P-TDP43LC. Supplementary Table 4 shows the measured relaxation values for all samples tested. No significant difference in either ^1^H T^1^ρ or ^15^N T^2^ was found between the WT and P-TDP43LC fibrils. T^1^ρ measurements for both protein fibril constructs each showed a biexponential decay with a fast and slow component of ∼ 1 and 9 ms, respectively. The ^15^N T^2^ was well-fit by a single exponential decay, and measured T^2^ values were ∼ 5 to 6 ms. These values are similar to that measured previously for fibrils formed by a slightly different TDP43LC construct, and thus also represent an ordered fibril core^34^. The lack of difference between wild-type and P-TDP43LC signifies a similar rigidity between the fibril *core* regions.

Despite no difference in cross-polarization detected relaxation times for the fibril core region, the multidimensional solid state NMR experiments that give rise to signals from rigid portions of the proteins in Figure 5C immediately revealed clear differences in chemical shifts. Amino acid types by spectral subregion are indicated and show that there are significant differences in the intense Ser signals, as expected from the five mutation sites. However, Figure 5C also shows large differences in Ile, Ala, and Gly, among other amino acid types. The Asp mutation sites for P-TDP43LC have nearest neighbor residues Asn, Gly, Met, and Lys. Therefore, the large differences we see in the difference spectra for amino acids like Ala and Ile cannot be attributed solely to mutational nearest neighbor effects, but rather indicates a structural rearrangement of distant amino acids.

To identify the rigidified residues on the protein sequence, we made sequence specific assignments as depicted in Supplementary Figures 10–14 and Supplementary Tables 6–13 from the 2D & 3D multidimensional heteronuclear cross-polarization based experiments. Signals were tabulated and input into the MCASSIGN 2b Monte-Carlo annealing algorithm^34,35^. Manual verification of chemical shift inter-residue correlations was performed to ensure robust assignments. Despite our best efforts, there remains unassigned resonances in each spectra due to a mixture of low signal-to-noise as well as ambiguity in assignments on the sequence from the low-complexity nature of the TDP43LC sequence. Therefore, there are regions of both fibrils that are rigidified but remain unidentifiable beyond general amino acid designations. In addition, the presence of multiple fibril polymorphs cannot be conclusively determined from these data for either wild-type or P-TDP43LC. However, we did not observe multiple NMR assignments for the same amino acid residues in the sequence, which would have signified multiple fibril conformations present in the sample.

In all, for WT-TDP43LC fibrils sequence specific assignments were made for residues: S317–N319, N378–I383, and S387–S395; and, for P-TDP43LC residues: G287–G298, G308–M311, and S389–A391. Additional signals in the WT spectra were determined to likely correspond to a SGN motif and, separately, a QNQ motif, each of which is found multiple times in the low-complexity TDP43LC sequence (see Supplementary Tables 10 and 11).

Figure 5D–E show the secondary chemical shifts for WT and P-TDP43LC where large negative values indicate β-strand regions unique to each construct. As shown in Figure 5, we identified from secondary chemical shift analysis β-strand secondary structure character for WT-TDP43LC from residues: S379–I383 and S387–G394; and, for P-TDP43LC: N291–G295. These predictions are in agreement with φ and ψ torsion angle predictions based on the TALOS-N analysis shown in Supplementary Figure 15^23,24^.

For both WT and P-TDP43LC, replicate solid state NMR samples and corresponding cross polarization-based spectra show nearly identical signals. Supplementary Figures 10 and 12 show the spectra from WT and P-TDP43LC fibril replicate samples with nearly identical spectra as for the samples used to make the assignments. Therefore, the procedure outlined here was found to reproducibly yield the same fibril conformer, or the same relative population of conformers from preparation to preparation.

Despite only obtaining partial NMR assignments, we can conclusively make some statements about the insights the rigid-residue and mobile-residue solid state NMR experiments uncover. One, both WT and P-TDP43LC assigned signals are consistent with β-strand rich fibrilization (Figure 5). Two, the assignments are consistent with multiple regions of the protein forming β-strands; both the region seen in the ex vivo cryo-EM maps and the “core 2” region we observed with a related wild-type TDP43LC construct^11^. Three, the spectra demonstrate that P-TDP43LC forms a different fibril construct than WT protein, and therefore phosphomimetic substitution changes which fibril conformations formed. We note that the P-TDP43LC fibril assignments have more overlap with the regions from ex vivo TPD43LC fibril samples derived from disease, although phosphomimetic substitution does not preclude residues S389–A391 from rigidifying sufficiently to appear in the cross-polarization solid state NMR spectra in Figure 5 F–J.

## Discussion

WT-TDP43LC fibrils formed in this work differ from previous His-tag linked WT-TDP43LC fibrils, Figure 5F–G^11^. This observation of different chemical shift assignments and thus a different fibril conformer between WT-TDP43LC and the 6xHis tagged WT-TDP43LC previously studied highlights the importance of flanking domains in protein amyloid fibril core selection. Despite the 6xHis tag domain not rigidifying into a well-formed secondary structure motif, the tagged version of WT-TDP43LC studied previously adopted different fibril conformations compared to the tagless WT-TDP43LC studied in this work^11^. Thus, a distant motif can perturb the free-energy landscape of the fibril formation process, selecting for a globally different fibril conformation. The effect of flanking domains on fibril core selection was commented on previously as well for the protein FUS, where it was demonstrated that a disordered domain can act as an entropic tail that modifies protein folding without folding itself^36^.

In addition, it was recently shown that full length TDP43 induced to form fibrils in vitro forms yet another fibril conformation compared to amyloids formed by fragments of the WT TDP43 C-terminal domain^18^. Excitingly, these researcher’s in vitro full-length TDP43 amyloid fibrils had good overlap with the ALS and FTD ex vivo fibril core region rigidification. However, the precise folding, the patterning of β-strands and turn regions within the in vitro core, was different than the ALS and FTD ex vivo strucutres^18^.

It is worth noting that the P-TDP43LC construct studied here, without modification to S369, is more suggestive of FTLD-TDP type A since this disease subtype has a lack of phosphorylation at this position^8^. Different phosphorylation patterns in FTLD-TDP type A versus ALS might be influencing the observed different fibril polymorph from each of these neurodegenerative diseases, although further work is necessary to identify the relationship between post-translational modifications, fibril formation, and disease pathology. A recent study on oxidized and partially phosphorylated TDP43 was also recently performed, which showed differential chaperone interaction behavior depending on post-translational modifications^37^. This study showed CK1δ phosphorylated TDP43 aggregated faster when also containing methoxidation modifications, and yet this same study showed little change in aggregation kinetics for CK1δ reacted TDP43 without methoxidation^37^. In addition, a previous study showed CK1δ reacted TDP43 aggregates more slowly than WT-TDP43. Therefore, aggregation of TDP43 appears to be highly sensitive to molecular environment and protein post-translation modifications, since even the impact of phosphorylation is different depending on the associated environmental conditions.

A procedure to generate the TDP43 ex vivo fibril conformers in vitro remain to be determined^3,4^. Such a procedure would greatly benefit in vitro research see ing to study the disease relevant conformations of TDP43 by not relying only on patient-derived tissue samples. This would aid efforts including but not limited to the development of drug-disaggregases and diagnostic imaging agents for early disease detection.

A similar study was recently performed on the neurodegeneration-associated amyloid-like fibril forming protein Tau^38^. The authors showed that different phosphomimetic patterns also lead to different fibril conformers. Interestingly, specific sets of Tau phosphomimetics led to an in vitro fibril structure that was closely related to the Progressive Supranuclear Palsy ex vivo amyloid structure^38^. This study, like the data presented here, shows the possibility of phosphorylation, for some systems, functioning as the switch between the formation of non-pathogenic to the pathologically observed fibril conformations. Phosphorylation might also select for TDP43 disease specific fibril conformations. As our studies in this paper show, the P-TDP43LC fibril conformation is different than WT-TDP43LC fibrils which shows that, in the context of the TDP43 C-terminal sequence, the addition of a few negative charges can cause a different fibril conformation to form.

## Conclusion

Previously, researchers have shown that TDP43 forms disease specific amyloid folds and that TDP43 undergoes neurodegenerative-disease specific phosphorylation patterns^3,4,8^. In this study, we sought to identify a potential link between these observations. We find via solution NMR that soluble, non-phase separated P-TDP43LC has nearly identical disordered regions as well as an α-helical region, just like wild-type protein^25^. In aggregation assays, despite being soluble and aggregating to a lesser extent as indicated by turbidity measurements, P-TDP43LC binds ThT prematurely before large aggregate formation. This led us to run mixed aggregation assays of P-TDP43LC with wild-type protein, and we observed no change in the aggregation kinetics of wild-type TDP43LC in mixed samples from 0 up to 50% P-TDP43. Therefore, under the conditions tested, P-TDP43LC does not “chaperone” nor interrupt WT-TDP43LC aggregation. Despite P-TDP43LC’s reduction in aggregation propensity in isolation, we show via electron microscopy that P-TDP43LC still forms fibril aggregates. Thus, the introduction of additional negative charges via phosphomimetics at the five mutated Ser sites does not inhibit the ability of TDP43LC to form fibrils. Lastly, we showed via solid-state NMR analysis that the fibrils formed by P-TDP43LC are different from WT-TDP43. Therefore, based on this in vitro result, it is possible that phosphorylation could select for a different TDP43 fibril conformation in vivo as well. The biological relationship between phosphorylation and TDP43 aggregation in neurodegenerative disease remains to be more thoroughly characterized, as well as the timing of phosphorylation in vivo relative to the formation of amyloid plaques.

## Supporting Information

Materials and methods section, supplementary figures of NMR spectra and assignment parameters, kinetic assays, microscopy images, as well as supplementary tables consisting of NMR chemical shift assignments, values, peak lists, and pulse program parameters.

## Acknowledgements

We thank Dr. Khaled Jami for helping optimize the TEV-cleavage for WT-TDP43LC as well as determining the shaking protocol for the ThT kinetic assays. We also thank Dr. Ping Yu, the UC Davis NMR Core Facility technical director, for assistance with the NMR spectrometers. Funding for NMR spectrometers is from the National Science Foundation, DBI-0722538. This study was fully funded by the National Institute of General Medical Sciences at the National Institute of Health through award R35GM142892 to D.T.M. The content is solely the responsibility of the authors and does not necessarily represent the official views of the National Institutes of Health.

