## Supplemental Methods, Data, Tables for "Phosphorylation Mimicking Mutations Cause TDP-43 to Adopt Different Fibril Conformations"

**Contents:**

1-11	Material And Methods

12-26	Supplementary Figures

27-48	Supplementary Tables

48-49	Supplementary References

**Hazard Statement**

No novel or unusual hazards were encountered in this research.

**Data Sharing**

Solution and solid state NMR assignments, in addition to being tabulated here, are additionally uploaded to BMRB entry #<>.

**Materials and Methods**

*Protein Preparation*

Wild-type and P-TDP43LC were recombinantly expressed in BL21DE3 E. Coli with residues 262-414, uniport accession number Q13148. All protein was quantified with UV-vis spectroscopy, utilizing the ProtParam computed extinction coefficient based off sequence (https://web.expasy.org/protparam/). P-TDP43LC had mutations from Ser to Asp at residues 379, 403, 404, 409, and 410 and was ordered from GenScript inserted into a pET-11a plasmid encoding ampicillin resistance. The wild-type protein plasmid was the same as that used previously^1^. Both constructs contained an N-terminal 6xHis tag with a TEV cleavage site, MSYYHHHHHHDYDIPTTENLYFQGAMD.

All bacterial growth media contained 100 $\mu$g/mL ampicillin. Cells from -80 °C stocks were streaked onto an LB Agar plate and grown overnight in a stationary incubator at 37 °C. For non-isotopically labeled protein expression, isolated cell colonies on the overnight LB Agar plate were transferred by scraping and pipetting up and down into LB media containing 1 % W/V glucose and grown overnight without shaking at 37 °C. The next morning, 80mL of the overnight LB growth solution was transferred to a final volume of 1L LB media containing 1 % W/V glucose and grown until an OD of 0.8-1 in a shaker incubator at 220rpm and 37 °C. Protein expression was then induced via the addition of Isopropyl-$\beta$-D-thiogalactoside (IPTG; Fisher) to 0.5 mM final concentration. After 3 hrs, the cells were pelleted by centrifugation at 6k rcf, flash frozen in N_2(l)_, and stored at -80 °C.

For ^13^C ^15^N isotopically labeled WT protein and P-TDP43LC expressions whose material went into the first two solid state NMR samples, the expression protocol was identical to the unlabeled expression until the 1L LB media was grown to an OD of 0.8-1. Then, 4x1L LB growth flasks were pelleted by centrifuging at 6k rcf 10 min, the supernatant was removed, and all four cell pellets were resuspended by pipetting into 1L M9 minimal media containing 1g ^15^N ammonium chloride and 2g uniform-^13^C $\alpha$D Glucose, along with 12.27g sodium phosphate dibasic heptahydrate, 2.7g potassium phosphate monobasic anhydrous, 0.5g NaCl, 2mM MgCl_2_, 0.1mM CaCl_2_, pH’d to 7.4. This solution was grown for 30 minutes at 37 °C, then IPTG was added to a 0.5 mM final concentration to induce expression. After 3hr of expression, the cells were pelleted at 6k rcf 10min, and flash frozen in N_2(l)_ and stored at -80 °C.

For the third P-TDP43LC solid state NMR sample, and the protein for the P-TDP43LC solution NMR experiments, a slightly different M9 expression protocol was used. 1L of M9+ solution contains, in addition to the description in the previous paragraph, 0.2g Yeast extract, 10mL of 100X MEM vitamin stock (Corning) and 0.1mM FeSO_4_. An agar plate was inoculated and grown according to the unlabeled expression protocol. Then, the next morning, cells were scraped into 5mL of unlabeled LB media containing 1% W/V glucose and grown until the OD was ~1 to 3. Then, 2 mL of 37 °C prewarmed M9+ solution was combined with 200uL of the LB growth and placed at 37 °C with 220rpm shaking until the OD was ~1.8. This growth was then added to a final OD of 0.053 in 50mL of 37 °C prewarmed M9+ solution and placed into the shaker incubator at 37°C 220 rpm shaking in an Erlenmeyer flask. The following morning after 17.5 hrs, the entire M9+ overnight culture was added to the rest of the M9+ media, which was prewarmed to 37 °C. This was grown at 37 °C 220 rpm until an OD ~ 1, then IPTG was added to 0.5 mM final concentration. After 4 hrs of expression, the cells were harvested via centrifugation at 6k rcf 10 min and flash frozen in N_2(l)_ and stored at -80 °C. For the P-TDP43LC solid-state NMR sample, this procedure was doubled and the protein was pooled following the *protein purification* procedure. This M9+ procedure results in theoretically higher isotopic labeling, since the cell cultures grew to high cell density on a diet largely consisting of isotopically labeled M9+ media rather than LB media.

*Protein Purification*

The cell pellet was removed from the -80 °C and thawed on ice for ~30 min. Then, ~25mL of lysis solution containing 6M guanidinium hydrochloride, Gu-HCl, 3 pellets of Pierce protease inhibitor tablets EDTA-free (Thermo-Fisher), 1% V/V Triton X-100, 500mM NaCl, 50 mM tris(hydroxymethyl)aminomethane (Tris-HCl) pH 7.5, and 0.25 mg/mL hen egg white lysozyme (Fisher) were used with an automatic pipettor to partially resuspend the cell pellet. Then, the resuspension was sonified on ice with a Branson SFX-250 pulsed at 0.3 s on, 3 s off, 20 min total “on” time, with a 1/4-inch microtip. This solution was centrifuged at 75,600 rcf for 30 min, 4 °C, and the supernatant was loaded onto a Bio-Rad NGC Quest 10 Plus liquid chromatography instrument. A 5 mL Cytvia Histrap FF IMAC column was used with the system pre-equilibrated with equilibration buffer, EQ, which contained 6M urea, 20 mM 4-(2-hydroxyethel)-1-piperazineethanesulfonic acid (HEPES) pH 7.5 and 500 mM sodium chloride. The column was washed with EQ buffer containing an additional 20 mM imidazole until the absorption at 280 nm neared baseline. Then, the protein was eluted with EQ buffer containing 200mM imidazole. The protein fractions were verified by SDS-PAGE and stored at 4 °C.

For WT- TDP43LC, the protein’s purification tag was removed as follows. First, the purification fractions were diluted into reaction solution to contain 20$\mu$M TDP43LC, 0.5 M Gu-HCl, 20 mM pH 6.2 NaPi, and 1mM DTT. Lab made Tobacco Etch Virus protease, TEV, was added from ~40 - 80 $\mu$M glycerol stocks to a ratio of 1:10 (TEV/TDP43LC), ^13^C and ^15^N isotopically labeled protein for rINEPT and bulk relaxation NMR experiments, and 1:50 (TEV/TDP43LC) for all other WT protein samples. The cleavage reaction was left in the dark at room temperature for 1 – 3 days, and 1mM additional fresh DTT was added each day. Then, the reaction was quenched by adding twice the reaction volume of denaturing, 8M urea EQ buffer (yielding close to 6M final urea) with 20 mM Imidazole. The solution was concentrated with an Amicon ultra 15 3k MWCO tubes according to manufacturer’s recommendations. The solution was then purified using a gravity column containing Bio-Rad Nuvia^TM^ IMAC nickel (II) resin equilibrated in EQ buffer with 20mM Imidazole. The flow-through was collected and contained purification-tag-removed TDP43LC as verified by SDS-PAGE, supplementary figure 1. Then, the wild-type protein was stored in aliquots at -80 °C.

For P-TDP43LC protein, the purification tag was removed in a similar manner to WT protein. First, the protein purification fractions from affinity chromatography were diluted into reaction solution to 40$\mu$M protein, 0.5 M Gu-HCl, 20 mM pH 7.2 NaPi, 100 mM NaCl, and 1mM DTT. Then, TEV was added from 7 $\mu$M glycerol stocks to a final ratio of 1:250 TEV / protein for the unlabeled, and first two solid-state NMR sample protein. A 1:500 ratio was used for the final solid-state NMR sample due to the observed high cleavage-efficiency of the lab-made TEV. The reaction was allowed to proceed for approximately 3 days, and 1mM more DTT was added each day. Then, the protein was purified with 5 to 7mL Bio-Rad Nuvia^TM^ IMAC nickel (II) resin equilibrated in EQ buffer containing 20 mM Imidazole. The flow-through was collected and contained P-TDP43LC as verified by SDS PAGE, supplementary figure 1. The purified protein in denaturing buffer solution was stored in the fridge at 4 °C until use. For the *aggregation assays* including the turbidity measurements described below, P-TDP43LC was stored at 4 °C for up to 1.5 years without visible aggregates forming in the tube. See the *kinetic assays* section and *solution NMR*, in the diffusion NMR experiments section, for how this protein was dialyzed and clarified by centrifugation to try to ensure no aggregated material via ensuring no OD_600_ scattering at ~20 $\mu$M protein and a UV cuvette path length of 1 cm. For the solution NMR and solid state NMR experiments, the P-TDP43LC protein was used within a few months of storage at 4 °C.

*Solution NMR*

P-TDP43LC was prepared by dialyzing the protein at 40 $\mu$M in 3K MWCO 6.4 mL/cm tubing (Spectrum) into 6M urea, 20mM HEPES pH 7.5, 500mM NaCl overnight. Then, the next day ~1 mL of the protein solution was dialyzed into 300mL 20mM MES pH 6.2 buffer in 6-8 MWCO, 0.32 mL/cm dialysis tubing (Spectrum) at room temperature. After ~2hrs, the dialysis tubing was transferred to 300mL more of the same buffer and left at room temperature. The next morning, the protein was harvested then spun at 14k rcf 10 min in an Eppendorf tube to remove preformed aggregates. The supernatant was removed without mixing, leaving behind a ~100uL volume at the bottom of the tube. Concentrate was then placed at a 90:10 ratio of protein solution to D_2_O. The final protein concentration in the NMR sample was 22.2 $\mu$M for the sample used with the iCBCANH experiment, and the concentration was 17.5 $\mu$M for the NMR sample used for all other solution NMR experiments. Spectra were recorded as detailed in supplementary table 5.

Signal to noise in subsequent NMR experiments could not be increased by simply making the protein more concentrated, as this dialysis procedure for the NMR samples always lost protein concentration due to pre-formed aggregates, which were removed prior to NMR analysis as detailed in the previous paragraph.

Manual assignments were made from the compiled shifts from the 3D spectra which was assisted for the identical sites with identical shifts compared to the WT-TDP43LC BMRB entry #26823^22^. ^1^H and ^15^N shifts from the HSQC spectra were used as to identify each chemical spin system, and then the tabulation of all the corresponding shifts from the 3D spectra were identified “i”, intra-residue, or “i-1”, inter-residue, type shifts. “i” shifts came from the CBCANH, iCBCANH, and HNCA, whereas “i-1” type shifts were observed in the CBCANH, HNCA, HNCO, and CBCA(CO)NH spectra. Matching the unique “i” or “i-1” shifts allowed for unambiguous assignments for most of the sequence, where for a majority but not all of the residues this corresponds closely to the values reported in BMRB entry #26823 for wild-type TPD43LC^2^.

Despite our best efforts, the peaks in grey in figure 1A and the unassigned residues in supplementary table 2 were unable to be assigned due to ambiguity in shifts. Residues G411 and G380 had identical C$\alpha$ and C$\beta$ shifts from inter-residue information, and therefore the unique ^15^N and ^1^H spin systems observed in the HSQC spectra could not be unambiguously assigned. Residues H264, R268, Q269, S332 were unable to be assigned, and neighboring assigned residues had rather low S/N, see supplementary table 1 and supplementary figure 2. Therefore, these unassigned peaks failed to be assigned due to insufficient signal to noise. Residue D379 was unable to be assigned unambiguously between two distinct D residues observed in the spectra that each had nearly identical carbon chemical shifts determined from inter-residue type peaks.

Secondary chemical shifts were calculated by comparing the differences between the assignment chemical shift values and the predicted random coil chemical shifts. Predicted shifts came from inputting the amino acid sequence into the Poulsen IDP calculator set to pH 6.2^3,4^. TALOS-N was ran with the assigned chemical shifts and sequence, and resulted in the secondary structure predictions in figure 1^5,6^.

The CSP plot was made by comparing the chemical shifts from the WT-TDP43LC assignments from the Fawzi groups assignments at 298K, BMRB entry #26823, with the assignments reported here for P-TDP43LC via the following formula for the chemical shift perturbation^2,7^.

$$CSP=\sqrt{\left( \Delta\delta_{H} \right)^{2}+0.14\left( \Delta\delta_{N} \right)^{2}}$$

Diffusion NMR was performed on P-TDP43LC protein made using the same procedure as detailed above for isotopically labeled preparations for assignment spectra except where otherwise noted. Diffusion experiments used non-isotopically labeled protein from the 4 °C stock of purified, tag cleaved P-TDP43. After the dialysis into 20 mM MES pH 6.2 buffer, the protein was harvested and additionally 0.22um sterile filtered prior to centrifugation to remove any preformed aggregates. The final P-TDP43LC protein concentration was 16.3 $\mu$M in 90% H_2_O / 10% D_2_O, with an additional 0.05% V/V dioxane internal diffusion reference.

For WT-TDP43LC, the tag-cleaved non-isotopically labeled protein was thawed and dialyzed directly from a -80 °C fraction at 54 $\mu$M protein and prepared for NMR otherwise according to the same procedure as the isotopically labeled P-TDP43LC solution NMR samples. The final concentration was 16.3 $\mu$M protein, in 90 % H_2_O / 10 % D_2_O, with 20mM MES pH 6.2 buffer and 0.05% V/V dioxane.

Spectra were recorded on a 800 MHz NMR Spectrometer with an Avance III console and the ledbpgppr2s pulse sequence with presaturation^8,9^. One 2D spectra was ran, with 800 $\mu$s gradient pulse length * 0.5, 400 ms diffusion time, acquisition time of 1.277 seconds, 32 scans, O1 centered on H_2_O near 4.7 ppm, and a relaxation delay of 5 s. The bipolar gradient powers were incremented from 2 - 95 % power in fixed increments of 10.33% for 10 total indirect points. The spectrum was then processed in nmrPipe with 5 Hz EM broadening, zero filled twice and phased, then exported to a text file for integration in a custom python script using a midpoint formula^10^. For dioxane, the integral was calculated from 3.7525 to 3.7475ppm, and for both TDP43LC samples, 7.3802 to 7.15 ppm. These diffusion spectra were referenced by setting the dioxane peak to 3.75 ppm.

Prior to integration, a baseline for each scan at every gradient power was individually solved for across the protein integration region. This was done by optimizing a fourth order polynomial, Baseline Intensity(ppm) = ax^4^ + bx + c, using scipy curve fit across the protein integration region, taking the 50 points, ~0.01ppm, at the end of the protein integration region bounds, 7.3802 and 7.15 ppm, to be the baseline data to fit. This polynomial was then subtracted from the data, which yielded a baseline near 0 intensity for each protein integration region, see supplementary figure 3 - 5 for the before and after baseline subtraction.

The following equation was fit to the integration across gradient powers with a custom python code using scipy_curve_fit;

$$I\left( x \right)=A e^{-decay\cdot x^{2}}$$

Where *I(x)* is the integrated intensity as a function of gradient power, *A* is an amplitude factor, and *decay* describes the decrease in signal with increasing gradient coil powers in the NMR experiment. The following equation was then used to find the radius of hydration^9^;

$$Rh_{\mathrm{unknown}}=C\cdot Rh_{\mathrm{dioxane}}\cdot\frac{\mathrm{decay}_{\mathrm{dioxane}}}{\mathrm{decay}_{\mathrm{unknown}}}$$

The decay was solved for using the *I(x)* equation above, Rh_dioxane_ = 2.12 Å, and the calibration factor *C* which was determined from reference protein horse myoglobin (Sigma product # M0630).

The *C* factor was found as follows. Non-isotopically labeled horse myoglobin lyophilized sample was dissolved in 700 uL of 20 mM MES pH 6.2, 25 mM NaCl, 5 mM TCEP buffered solution by mixing gently with a rotary mixer for 1 hr and left at room temperature for 14hrs. Then, the sample was 0.22 $\mu$m sterile filtered and the protein concentration was measured by UV-vis to be 3 mM. 495 $\mu$L of this solution was then combined with 55 $\mu$L D_2_O and 0.1% V/V 1,4 dioxane. The horse myoglobin diffusion experiment also used the ledbpgppr2s pulse program with the same experimental parameters as P-TDP43LC as detailed above, and processed according to the same procedure except the myoglobin signal was integrated from 9.714 to 6.561 ppm as the amide backbone had better signal intensity than the TDP43LC samples. This resulted in a measured Rh = 20.85 Å, therefore to obtain the Horse Myoglobin literature value for Rh = 20.4 Å, C = 0.978^11^. This allowed for the Rh of WT and P-TDP43LC to be measured as reported in the results section from the data in supplementary figures 3-5. The error in measurement is the standard deviation that results from the propagation of the scipy curve fitting errors for all the decay values which came from fitting the I(x) equation to the data. The error propagation use the multiplication / division rule, through the Rh_unknown_ relationship to calculate the final standard deviation as reported in the main text.

*Bright Field Microscopy*

An Olympus BX51 light microscope with differential interference contrast (DIC) filters, a 40X objective lens, and a Diagnostics Instrumets RT Slider camera with 6 megapixel sampling was used. 2.5 - 4 $\mu$L protein solution was placed directly onto an untreated glass side (Fisher Scientific) without a coverslip and imaged.

*Transmission Electron Microscopy*

5 $\mu$L of protein solution was placed onto glow-discharged ultrathin carbon film with lacey carbon support EM grids (Ted Pella, both 01824G and 018430F were used) for 2 minutes. A chemwipe was used to wick the solution, then 2 x 10$\mu$L H_2_O was used to rinse the grid, wicking after each application, then 5$\mu$L of 3 % W/V Uranyl-Acetate solution was added for ~12s, then wicked. The grids were imaged on a FEI Talos 120C microscope.

*Aggregation Assays*

First, TEV-cleaved WT and P-TDP43LC were diluted to 40$\mu$M protein in either salt (20mM NaPi pH 7.4, 200mM NaCl) or no-salt buffer (20mM NaPi pH 7.4). Then, the protein was dialyzed into either salt or no-salt buffer with 0.32 mL/cm 6-8kDa MWCO dialysis tubing (Spectrum) via a two step dialysis, ~7hrs then ~18hrs, at a ratio of <1% protein solution to buffer for each step. Then, the protein was harvested and a 3$\mu$L fraction was imaged with bright field microscopy, figure 2C. The protein solution was centrifuged at 14k rcf for 10 min to remove preformed aggregates, figure 2C, then quantified with UV-vis and diluted in respective salt or no-salt buffer to the specified concentration. For the turbidity measurements and kinetic ThT assay in figure 2, the protein was diluted to 13$\mu$M. For the mixed assay in figure 3, the protein was diluted to 12$\mu$M.

For the ThT assays in figures 2 and 3, a 96-well black polystyrene plate (Fisher #12566620) was used. The outer two rings of wells were filled with 90$\mu$L of H_2_O, and the plate was pre-inucbated to 37 °C. ThT was prepared with dissolving powdered ThT into 20 mM NaP_i_ buffer, 0.22$\mu$m sterile filtered, then quantified with a UV-vis extinction coefficient of $\epsilon$_412nm_ = 36000 M^-1^cm^-1^, and then diluted to yield 40$\mu$M ThT solution. 60$\mu$L of this freshly made 40$\mu$M Thioflavin-T, ThT, (ACROS Thermofisher) in 20 mM NaP_i_­ pH 7.4 buffer was placed into each sample and buffer blank well. 30$\mu$L of protein solution is added to the 60$\mu$L ThT solution and pipetted up and down gently to mix. For buffer blanks, 30$\mu$L of outer dialysis buffer is added to 60$\mu$L of ThT buffered solution and mixed. Clear polyolefin sealing tape (Thermo # 232702) was used to avoid evaporation during the assay. A BioTek Synergy H1 plate reader was used with 9 nm bandpass filters, excitation at 440 nm, emission recorded at 485 nm, read height 5.75 mm, receiver gain of 100, read from top, and incubation set to 37 °C during the assay. For the assay in figure 2, every 4 minutes had a 30 s burst of linear shaking at 1096 cpm (1mm). For the mixed assay in figure 3, every 5 minutes had a 30s burst of linear shaking at 1096 cpm (1mm). A blank well was run alongside n = 3 protein wells for each condition tested.

Data was processed with a custom python and included baseline subtraction from each respective condition’s blank well. For the kinetic parameter fitting in figure 3 and supplementary figure 6, scipy was used with the curve_fit functionality to yield a best-fit sigmoid to the data from each individual protein well. The fitting used the equation described previously^12^:

Y(t) $=Y_{i}+m_{i}t+\frac{Y_{f}+m_{f}t}{1+exp\left( -\left( t-t_{1/2} \right) \right)/\tau}$

$$t_{lag}=t_{1/2}-2$$

Where *Y(t)* is the ThT intensity over time at 485nm, *Y_i_* is the initial intensity, *m_i_* is the slope at initial time points, *Y_f_* is the intensity at the end of the assay, *m_f_* is the slope of intensity after the exponential phase, *t_1/2_* is the time it takes to reach half the maximum intensity, $\tau$ is related to the inverse of the elongation rate, *k_app_*, and *t_lag_* is the time at which the exponential phase of fibril growth begins, i.e. right before the rapid increase in ThT signal^12^.

For the turbidity measurements in figure 2, 13$\mu$M protein in either salt or no-salt buffer was placed in a pre-incubated clear polystyrene (Falcon #353072) 96-well plate with sealing tape (Thermo # 232702) and ran at 37 °C with 30s linear shaking every 4 minutes for 21 hours. Due to negligibly intense turbidity readings in the plate reader assay, more sensitive *endpoint* turbidity measurements were made after the samples aggregated with an Eppendorf Kinetic UV-vis Biospectrometer from n=3 separate wells for each condition tested. A plastic UVette (Eppendorf) with b = 1 cm was used with 60 uL solution, which was pre-mixed by pipetting up and down with care to avoid any air bubble formation. An independent two-tailed t-test was ran with the scipy package stats, using the ttest_ind command with equal_var set to False. The asterisk denotes a difference at the 95% confidence level, figure 2.

*Fibril Formation*

WT and P-TDP43LC were induced to form fibrils under identical conditions to each other with a procedure highly similar to the previously described “flash-dilution” method^13^. In brief, the protein was diluted to 1.7 mg/mL, 110$\mu$M (wild-type TDP43LC) and 1.725 mg/mL, 110$\mu$M (P-TDP43LC) in fibril buffer, 20 mM NaPi pH 7.4 and 200 mM NaCl. The protein was buffer exchanged with an Amicon Ultra-15 or Amicon Ultra-4, depending on the total volume of fibrils needed, of 3 kDa MWCO centrifuge filters according to the manufacturer’s buffer exchange repetitive centrifugation procedure. Brief centrifugation periods were followed by successive resuspension steps in fibril buffer, repeated until the initial buffer remaining was less than 5%. Then, the solution was brought to volume such that the protein concentration was returned to 110 $\mu$M and this solution was placed on an Eppendorf ThermoMixer at 300 rpm, room temperature. LLPS sometimes occurred during the volume additions during buffer exchange, figure 4, but after the final step, protein particulates were observed, supplementary figure 8. After 7-8 days of incubation, the protein was sonified with a Branson SFX-250 sonifier pulsed at 1 s on, 1 s off, for a total time of 30 s with a 1/8-inch microtip, pausing after the first 20 s of this period to let the solution sit for 1 min, then the final 10 s of sonification was ran. The fibril solution was then incubated for 7 additional days at 300 rpm at room temperature on an Eppendorf ThermoMixer. Fibrils were imaged with bright field and transmission electron microscopy, figure 4 and supplementary figure 8.

This procedure was used twice to make replicate WT-TDP43LC fibril samples, and three times for P-TDP43LC fibrils for replicate solid state NMR experiments.

*Solid State NMR*

After the fibril formation procedure, 8-16 mg of ^13^C and ^15^N isotopically labeled fibrils were pelleted by centrifugation at 268k rcf for 1.5 hr iteratively into a MLS 50 centrifuge tube with a swinging bucket rotor. The supernatant was removed after each step. After the fibrils were pelleted, the fibrils were scraped into a ssNMR rotor, a thin-walled 3.2 mm pencil style zirconia rotors (Revolution NMR) that contained a ~1 mm thick PFTE disk to center the sample in the rotor, as was necessary for all samples except wild-type sample 1 which used no PFTE disk, see supplementary tables 14-15 for which sample was used for each NMR experiment. Between scrapes of the fibrils into the rotor, the rotor was centrifuged in a swinging bucket rotor for ~5 s at < 3k rcf to pack the sample in the rotor. Then, the rotor was centrifuged at 25k rcf for 2-3 hrs in a swinging bucket rotor to pack the fibrils in the rotor. Then, the rotor top cap was attached with cyanoacrylate glue gel to seal the sample and prevent dehydration. All of the solid-state NMR experiments were recorded on an 18.8 T Bruker NMR magnet with an Avance III console and a BlackFox probe. All solid state NMR samples used an external ^13^CO labeled powdered alanine sample, with the carbonyl peak was set to 179.8 ppm.

The magic-angle spinning rate was set to 13 kHz. For cross-polarization based experiments, the sample chiller was set to -40 °C and the sample heater to -30 °C. The combination of heating, MAS spinning rate, and cooling air provided a sample temperature of ~0 °C based on calibration of the system with ^79^Br^14^. For solid state NMR experiments reporting of dynamic regions of the protein, INEPT-based experiments, the sample chiller was set to -20 °C and the sample heater to 0 °C. This corresponds to a net temperature of roughly 30 °C based on calibration of the system with ^79^Br^14^. All solid state NMR multidimensional experiments were collected with States-TPPI for the indirect dimensions and DQD for the direct dimension, except for the TOBSY spectra’s indirect dimension which used TPPI data collection.

For the relaxation experiments, the data was processed and referenced in Topspin 4.1.4. 20 Hz exponential line broadening was applied. Relaxation values were calculated via custom Python/SciPy fitting the integrals to the following equation for ^15^N T2 from the NCA experiment^15^:

I(t) $=I_{0}\exp\left( -t/T \right)$

Where *I(t)* is the integrated intensity for each 1d slice, *I_0_* is the intensity with no relaxation delay, *t* is the time delay, and *T* is the relaxation parameter. For both WT-TDP43 and P-TDP43 sample 2, the 1^st^ data point was removed prior to fitting, as the 0s T2 time had a lower integrated intensity than the 2^nd^ data point.

For ^1^H T1$\rho$, a biexponential was fit;

I(t) $=I_{a}\exp\left( -t/T_{a} \right)$ + I_b_ exp(-t/T_b_)

Where *I_a_* and *T_a_* are the integrated intensity and relaxation constant for the first component, and *b* represents the second component.

*Solid-State NMR Assignments*

Assignments were made with the 2D and 3D shifts from the heteronuclear triple resonance experiments detailed in supplementary tables 14-15. MCASSIGN 2b was used to generate sequence specific assignments for both WT and P-TDP43LC constructs^16^. MCASSIGN 2b was ran with parameters for “good”, “bad”, “edge”, and “used” of, respectively, (0, 10), (10, 60), (0, 8), and (0, 2) over 200 steps with 10^7^ iterations per step. 50 independent runs with unique random number seeds were used. The MCASSIGN assignments were manually verified for agreement between the signal tables from NCACX/CC DARR, NCOCX, and CANCO spectra, see supplementary tables 7 and 13. For P-TDP43LC fibril residue assignments but not WT-TDP43LC fibril assignments, G287-F289 and G294-G295 assignments from MCASSIGN yielded multiple potential matches due to the uncertainties in the MCASSIGN tables. Therefore, for these residues, best possible assignments were manually made, see supplemental tables 12 and 13.

To check for multiple conformations of the already-assigned-once residues in the sequence, MCASSIGN was reran with the same parameters except for the removal of the already assigned resonances. This did not lead to any additional sequence specific assignments for either WT or P-TDP43LC fibrils.

Secondary chemical shifts were calculated from the fibril solid state NMR shifts like the solution NMR assignments, see previous section. Torsion angle values in supplementary figure 15 are from TALOS-N predictions for “strong” and “generous” confidences^5,6^.

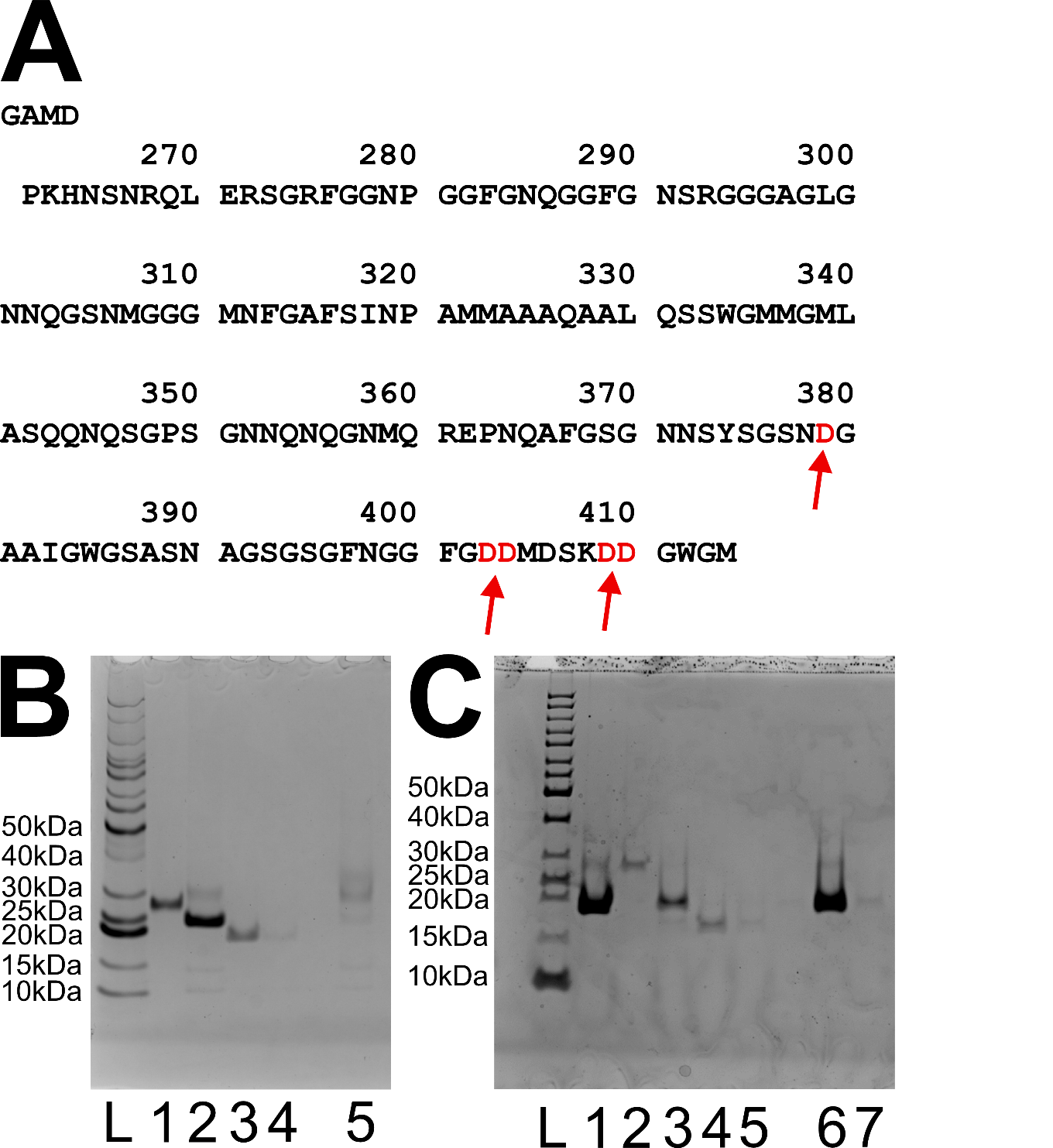
**Supplementary Figure 1. Sequence and SDS-PAGE.** A) Sequence map of the construct of P-TDP43 following TEV cleavage. Red and arrows highlight the sites mutated from Ser in the wild-type to Asp in P-TDP43LC. B) P-TDP43LC 4% to 10% bis-acrylamide gradient gel. L = protein standards ladder, 1 = TEV protease stock, 2 = P-TDP43LC before cleavage, 3 and 4 = flow-through fractions from reverse affinity chromatography, 5 = Elution fraction from reverse affinity chromatography (un-cleaved impurities). C) Wild-type TDP43 4% stacking / 10% resolving bis-acrylamide gel. L = protein standards ladder, 1 and 3 = WT-TDP43 before cleavage, 2 = TEV protease stock, 4 and 5 = flow-through fractions from reverse affinity chromatography, 6 and 7 = elution (un-cleaved impurities) fractions from reverse affinity chromatography.

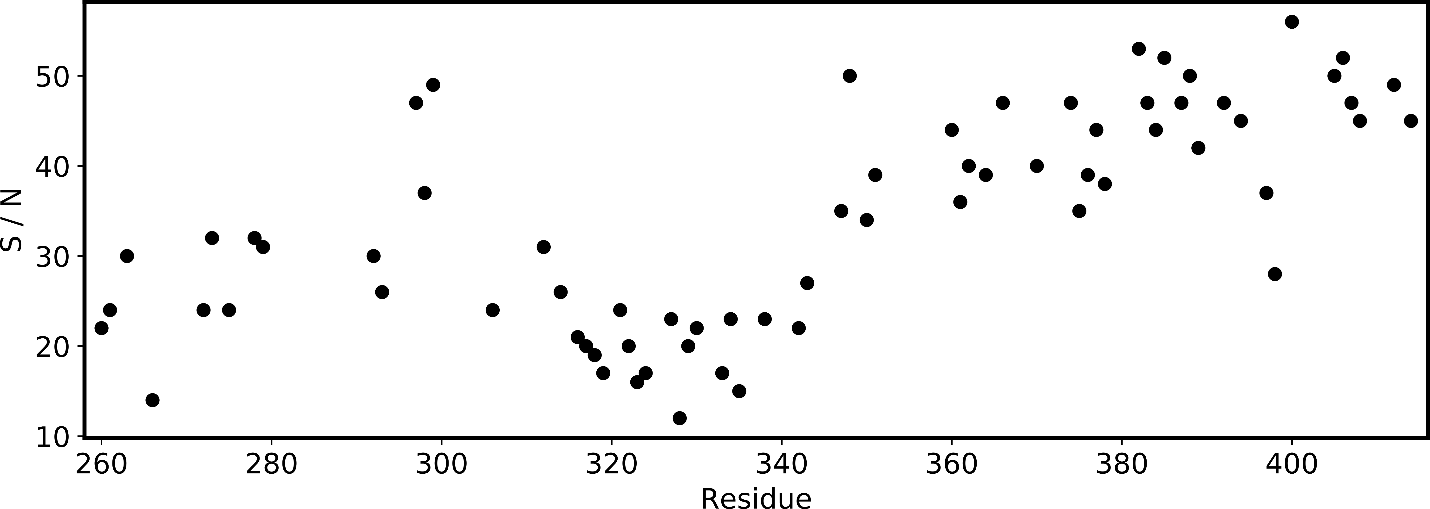
**Supplemental Figure 2. Solution NMR analysis.** The signal to noise ratio from the non-NUS (regularly sampled) fHSQC spectra as output by NMRFAM-SPARKY^17^. A dip in S/N at the N-terminus and the helical region from residues 320 to 340 is an indicator of increased protein structure for these regions, as partially structured regions in a disordered protein have increased correlation times which leads to lower signal to noise for these residues.

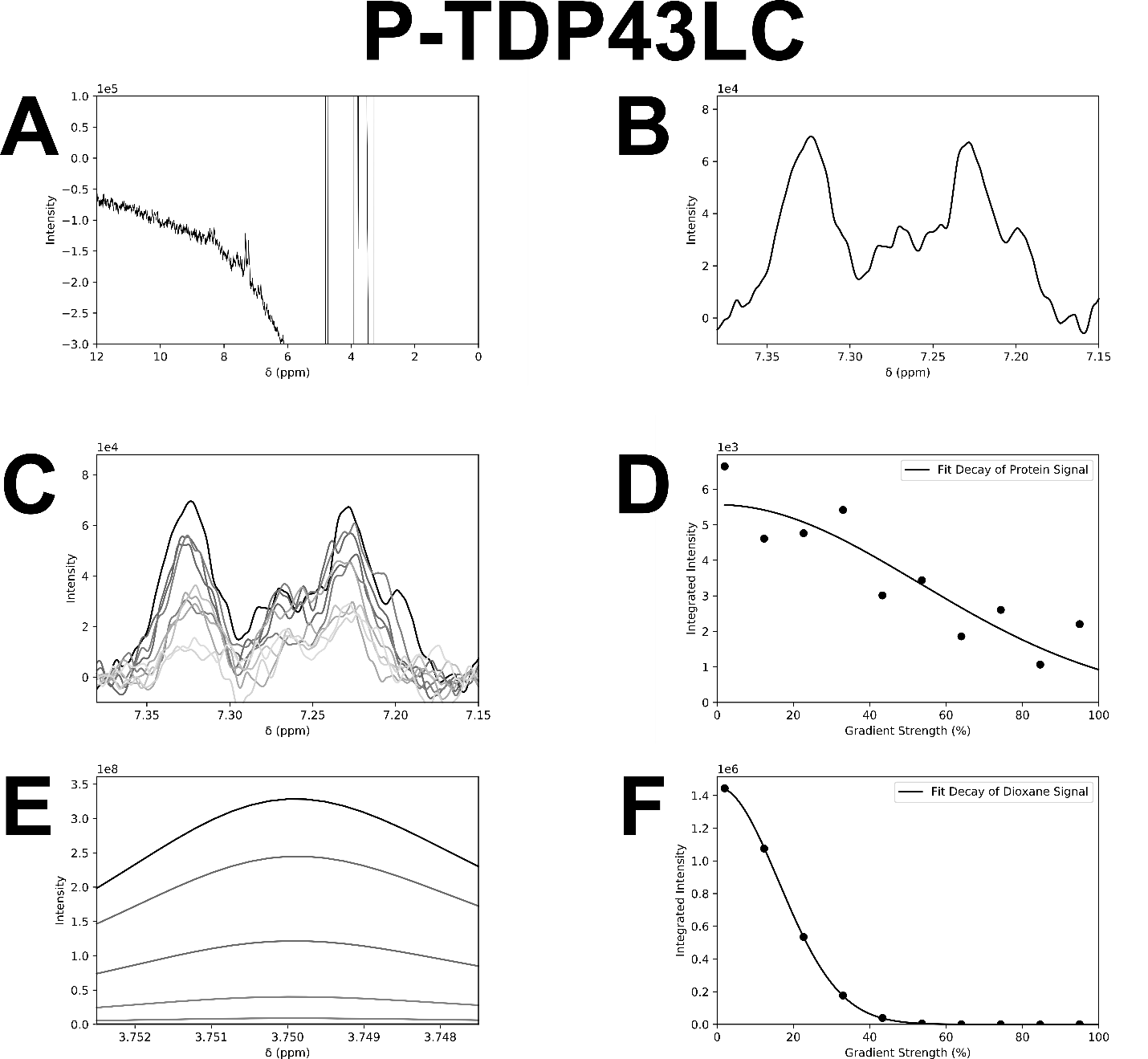
**Supplementary Figure 3. Diffusion NMR Data allows for the Measurement of Radius of Hydration of P-TDP43LC.** A) The first ledbpgppr2s scan with 2% gradient power for soluble P-TDP43LC protein shows the need for baseline fitting for the amide protein signal near 7.2 ppm, despite best efforts at phasing. B) The protein integration region following baseline subtraction demonstrates a baseline near zero at the edges of the protein signal. C) Overlayed scans from 2%, black, to 95%, grey, gradient power shows the diffusion-based signal decay at higher gradient powers. D) Integrated regions from C), fit to an exponential decay function. E) 1, 4 dioxane diffusion standard integrated region, overlayed scans from 2%, black, to 95%, grey, gradient power. F) Integrations and curve fit from E).

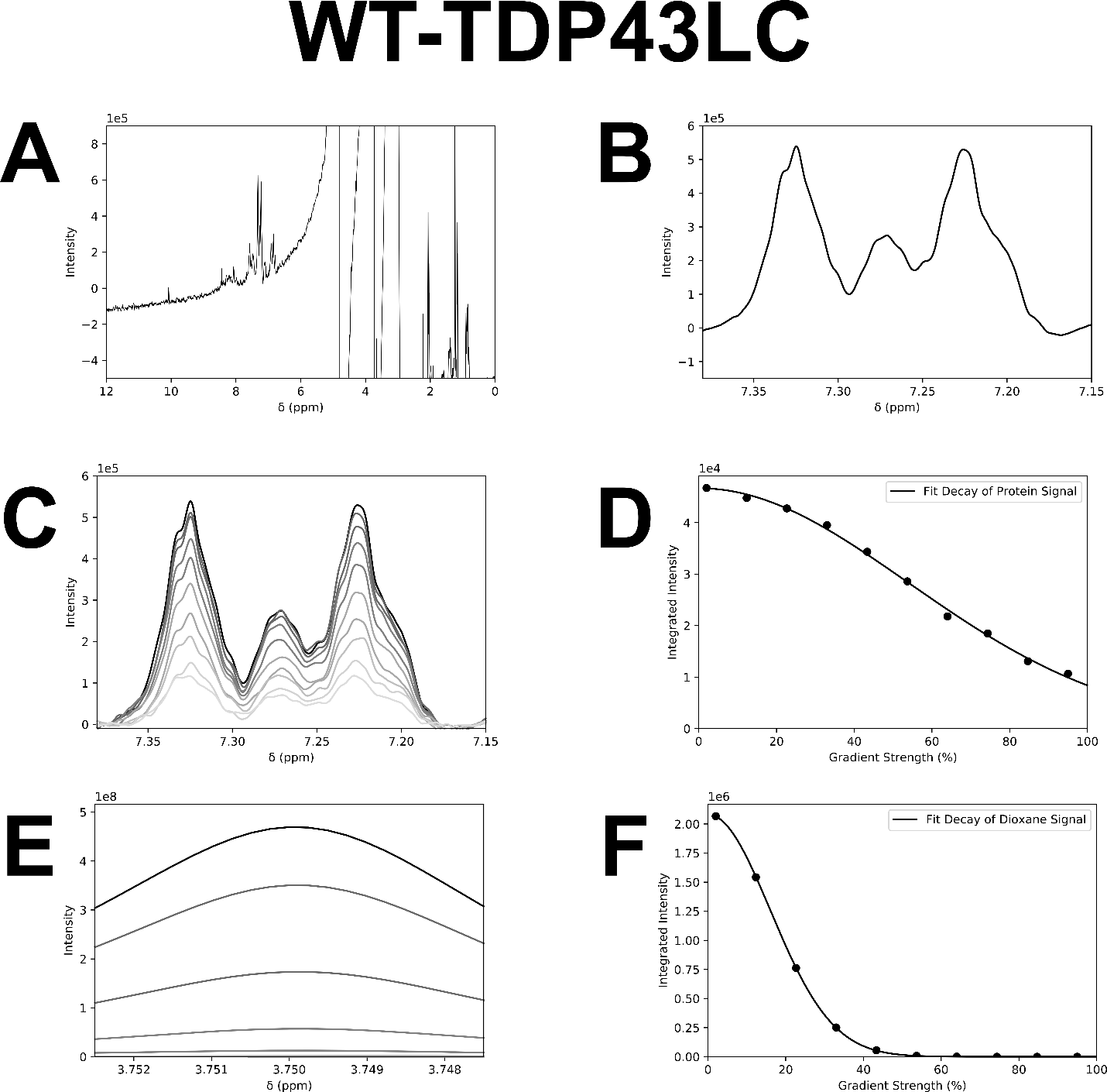
**Supplementary Figure 4. Diffusion NMR Data allows for the Measurement of Radius of Hydration of WT-TDP43LC.** A) The first ledbpgppr2s scan with 2% gradient power for soluble WT-TDP43LC protein shows the need for baseline fitting for the amide protein signal near 7.2 ppm despite best efforts at phasing. B) The protein integration region following baseline subtraction demonstrates a baseline near zero at the edges of the protein signal. C) Overlayed scans from 2%, black, to 95%, grey, gradient power shows the diffusion-based signal decay at higher gradient powers. D) Integrated regions from C), fit to an exponential decay function. E) 1, 4 Dioxane diffusion standard integrated region, overlayed scans from 2%, black, to 95%, grey, gradient power. F) Integrations and curve fit from E).

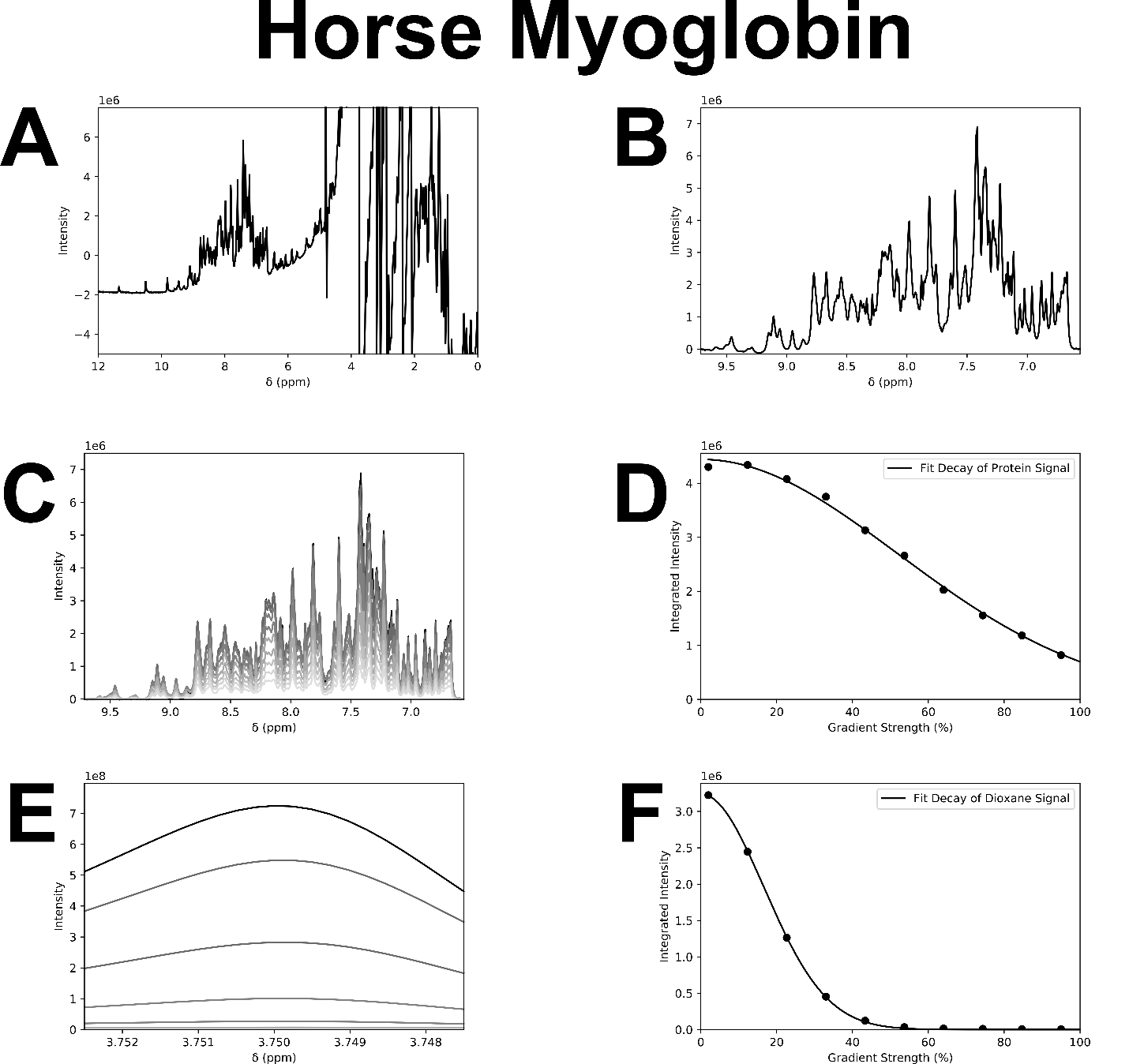
**Supplementary Figure 5. Horse Myoglobin Diffusion NMR measurements.** The same as supplementary figure 3 and 4, except on standard protein horse myoglobin which was used to calibrate the experiment to ensure proper radius of hydration measurements. A) The first ledbpgppr2s scan with 2% gradient power. B) The protein integration region following baseline subtraction demonstrates a baseline near zero at the edges of the protein signal. C) Overlayed scans from 2%, black, to 95%, grey, gradient power shows the diffusion-based signal decay at higher gradient powers. D) Integrated regions from C), fit to an exponential decay function. E) 1, 4 Dioxane diffusion standard integrated region, overlayed scans from 2%, black, to 95%, grey, gradient power. F) Integrations and curve fit from E).

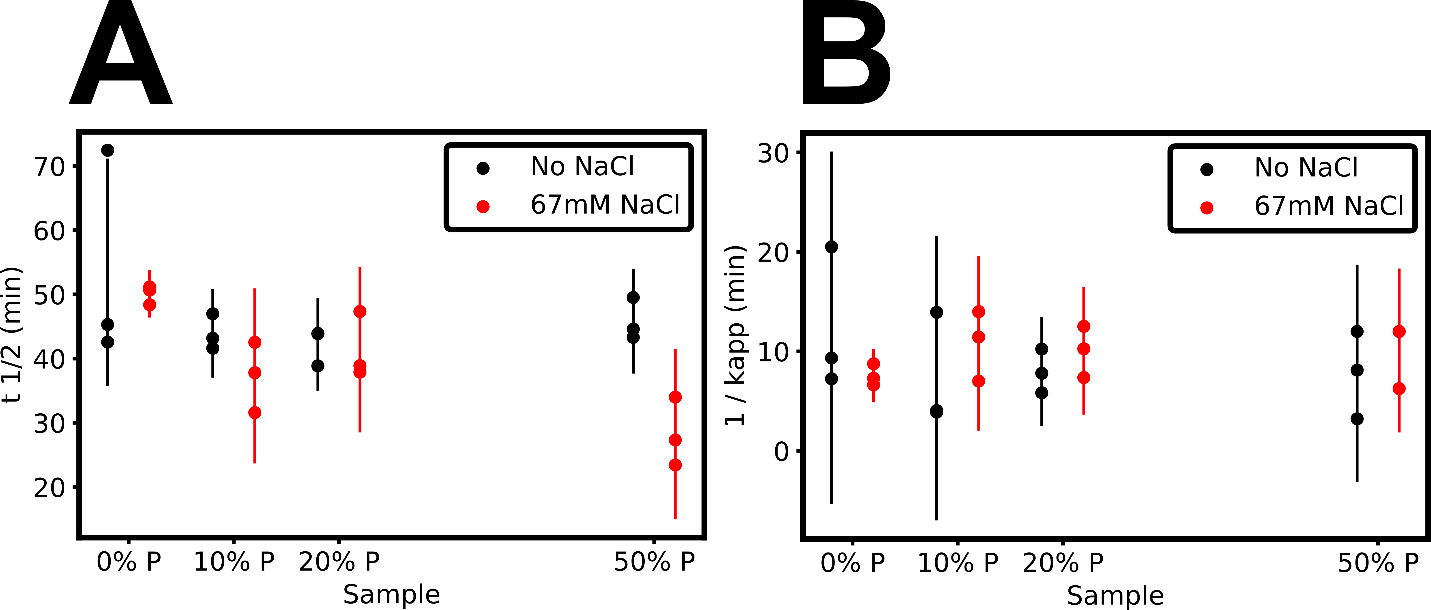

**Supplementary Figure 6. Kinetic Parameters from ThT Mixed Assays.** A) t_1/2_ versus percent P-TDP43LC mixed with wild-type protein. B) Measured $\tau$, i.e. the inverse of k_app_, versus percent P-TDP43LC.

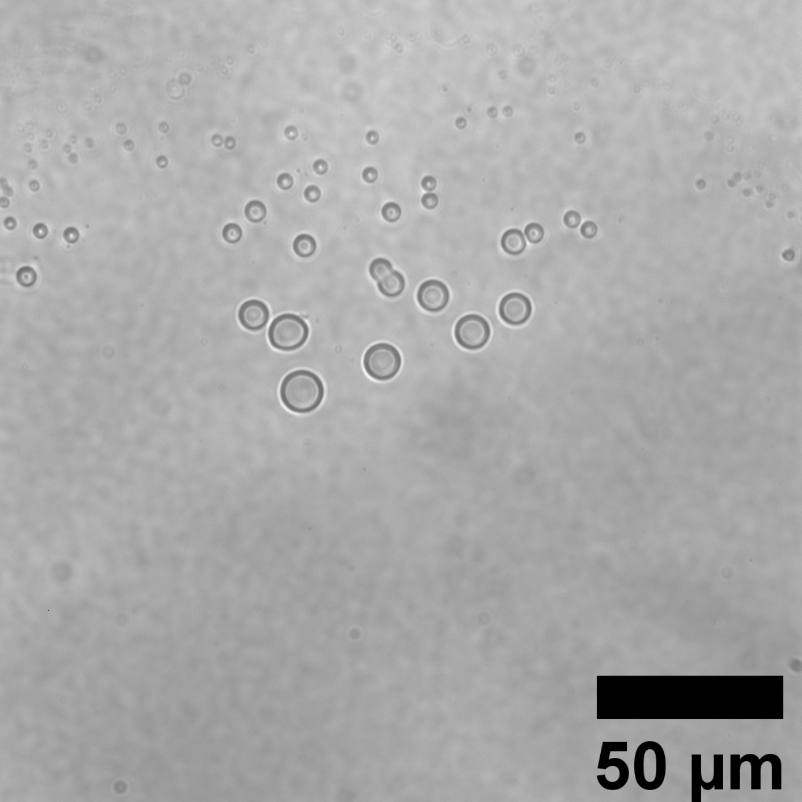
**Supplementary Figure 7. P-TDP43LC undergoes LLPS.** Bright field microscopy image of P-TDP43LC at 2.5 hrs dialysis into the *solution NMR* protein preparation procedure, with 40 $\mu$M protein. The solution at this point partially through the dialysis is an intermediate between the denaturing EQ buffer, described in the methods section, and the NMR buffer, 20 mM MES pH 6.2.

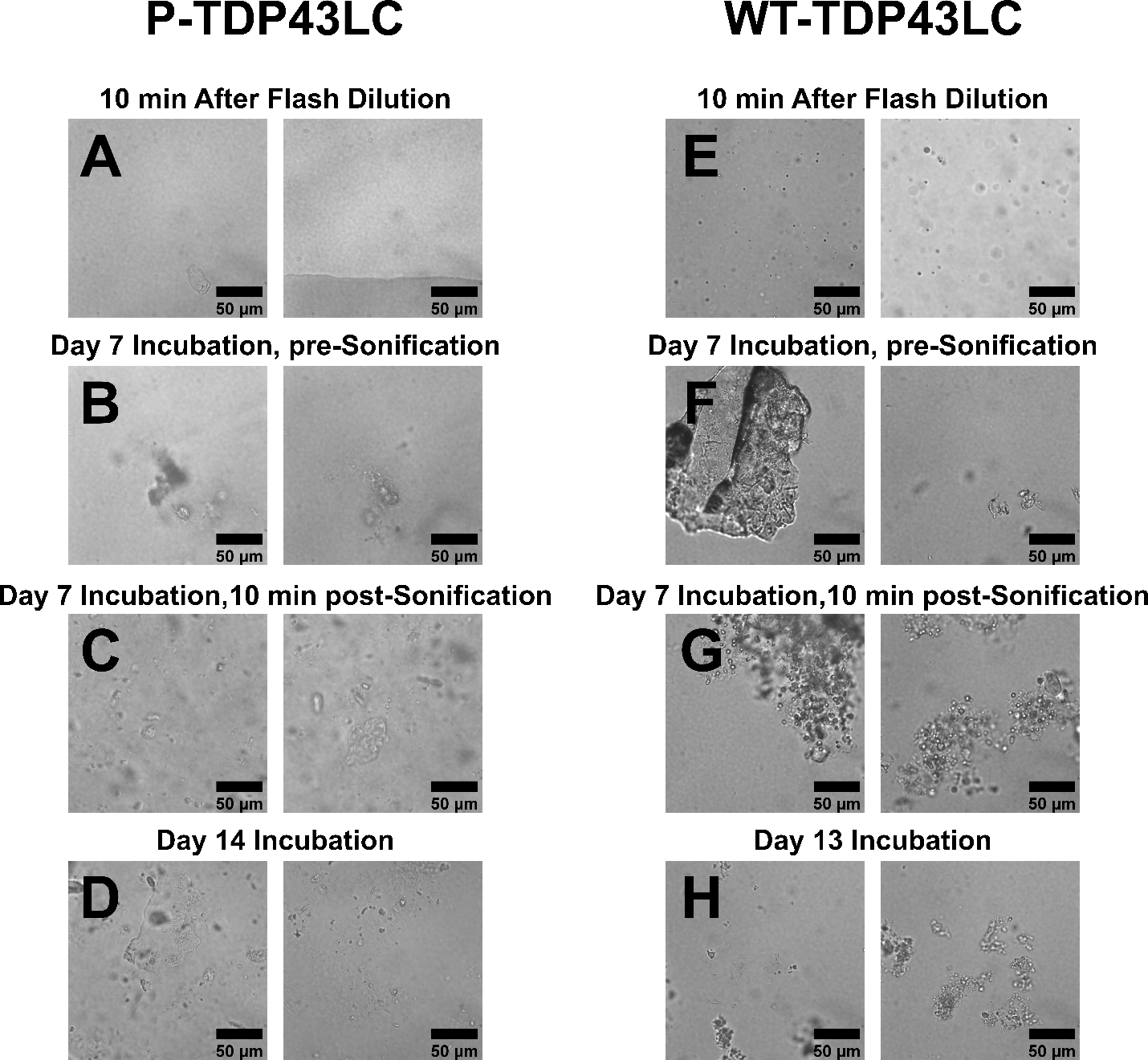
**Supplementary Figure 8. Bright Field Microscopy Fibril Formation Timecourse.** A-D) P-TDP43LC fibril formation at the specified times during the flash dilution fibril formation method. E-H) WT-TDP43LC fibril formation at the specified times during the flash dilution fibril formation method.

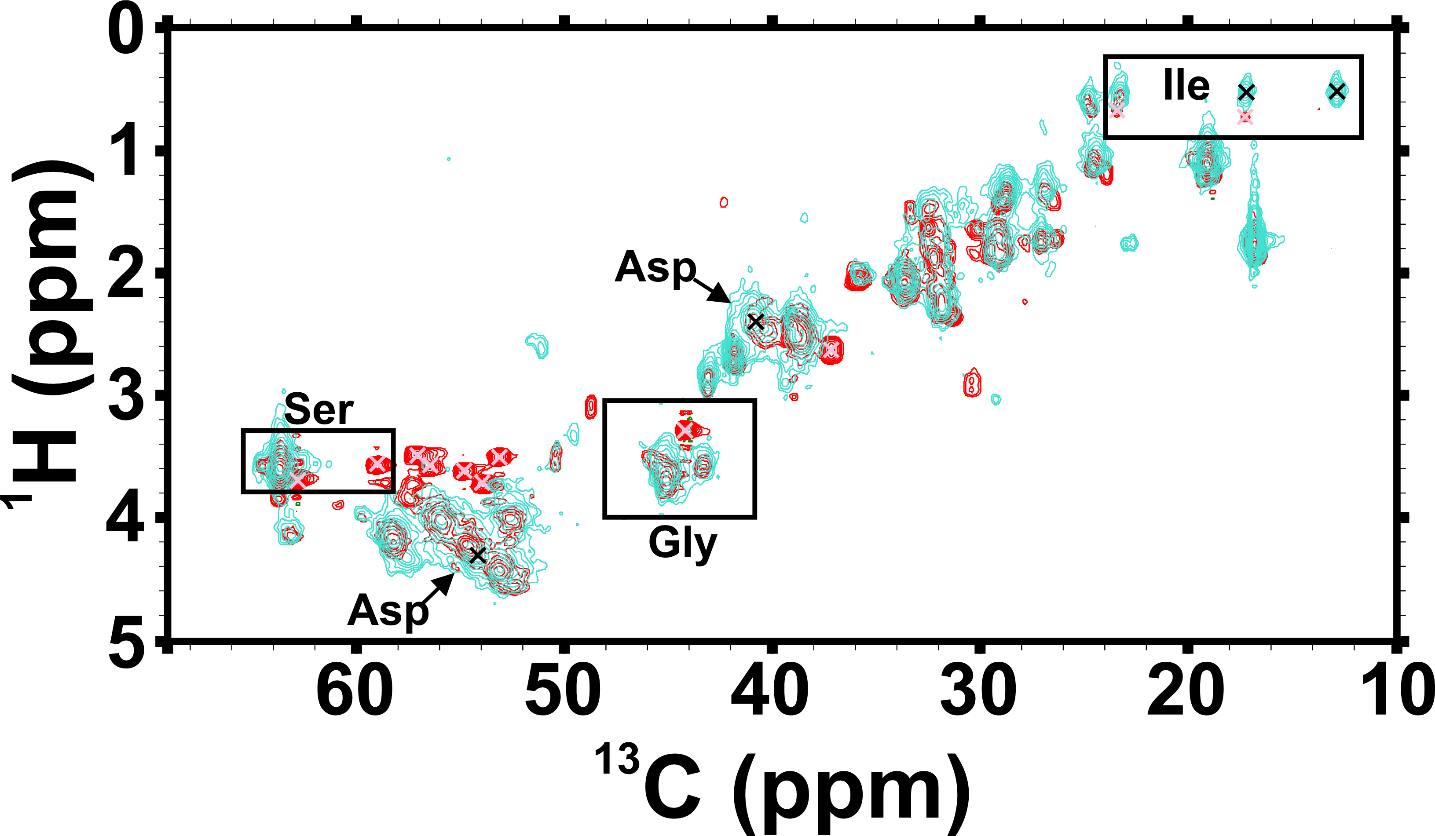

**Supplementary Figure 9. Solid-state NMR rINEPT Spectral Overlay shows Differences in Dynamic Residues.** Turquoise is the P-TDP43LC fibril sample and red is the wild-type fibril sample. Some key differences are highlighted on the plot, with black peak ‘x’s depicting unique P-TDP43LC peaks, and pink ‘x’s depict some of the unique wild-type peaks. Boxed regions show likely amino acid identities for the indicated peaks.

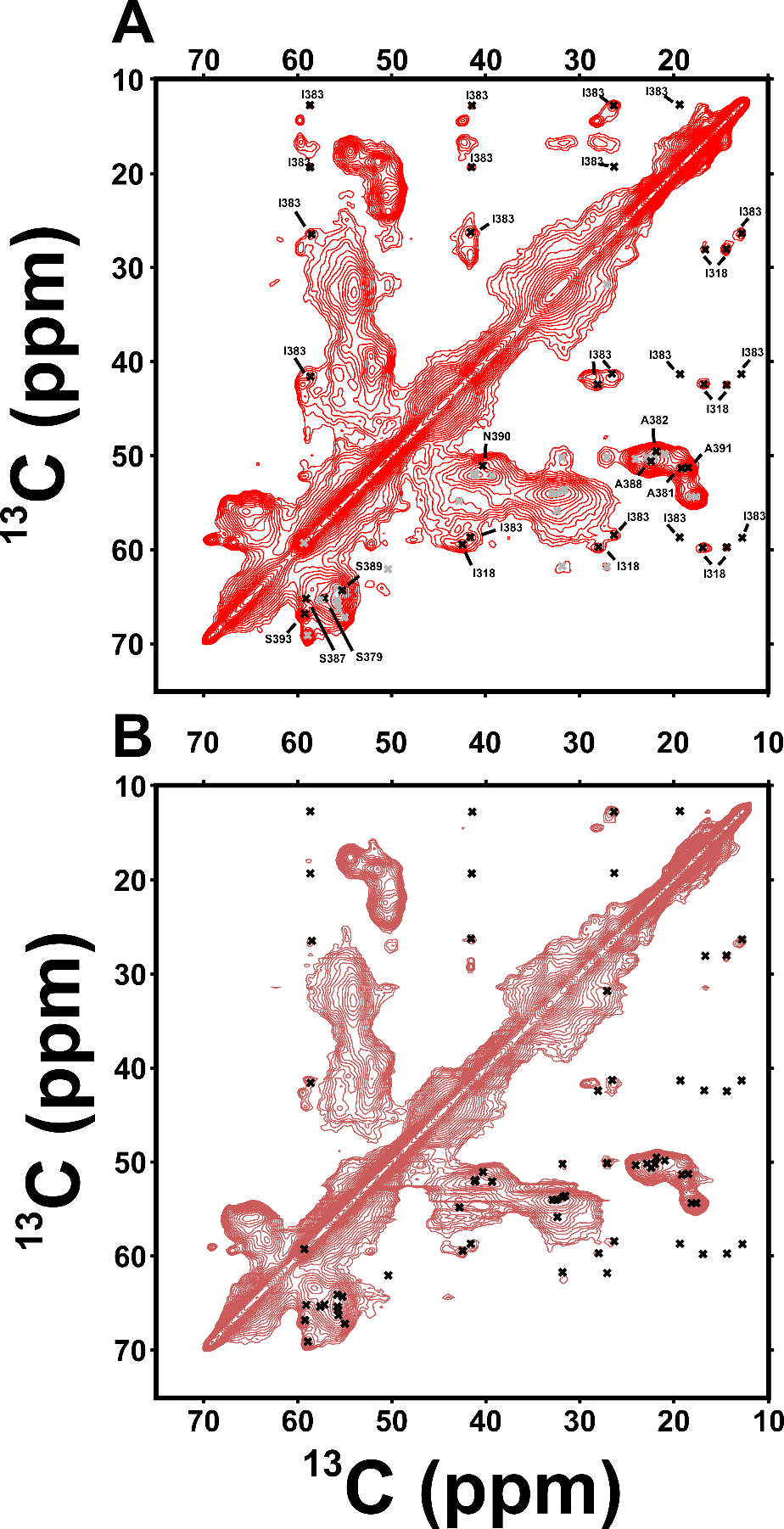
**Supplementary Figure 10. Assignments of the Rigid Regions of WT-TDP43LC Fibrils and Sample Replicate.** A) and B) Cross polarization based ^13^C-^13^C DARR solid state NMR spectra with 50ms mixing. A) Sample 1 of the WT fibrils, where grey x’s represent unassigned peaks and black represents sequence specific assignments. B) Sample replicate 2 of the WT fibrils, with the assigned and unassigned peaks from A) shown as black x’s.

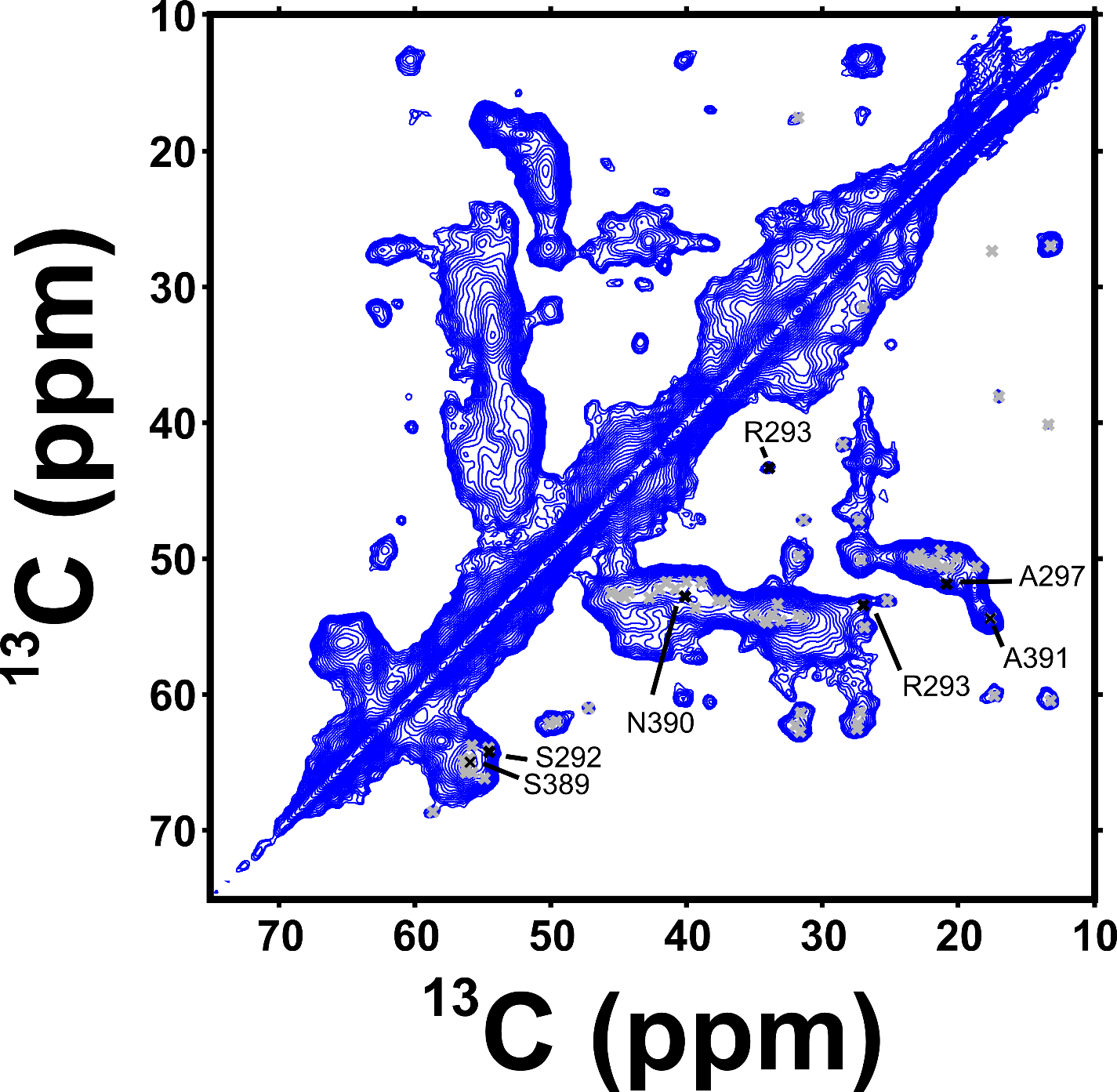

**Supplementary Figure 11. Assignments of the Rigid Regions of P-TDP43LC Fibrils.** Cross polarization based ^13^C-^13^C DARR solid state NMR spectra with 50ms mixing. Grey x’s correspond to unassigned peaks, and black x’s denote assignments.

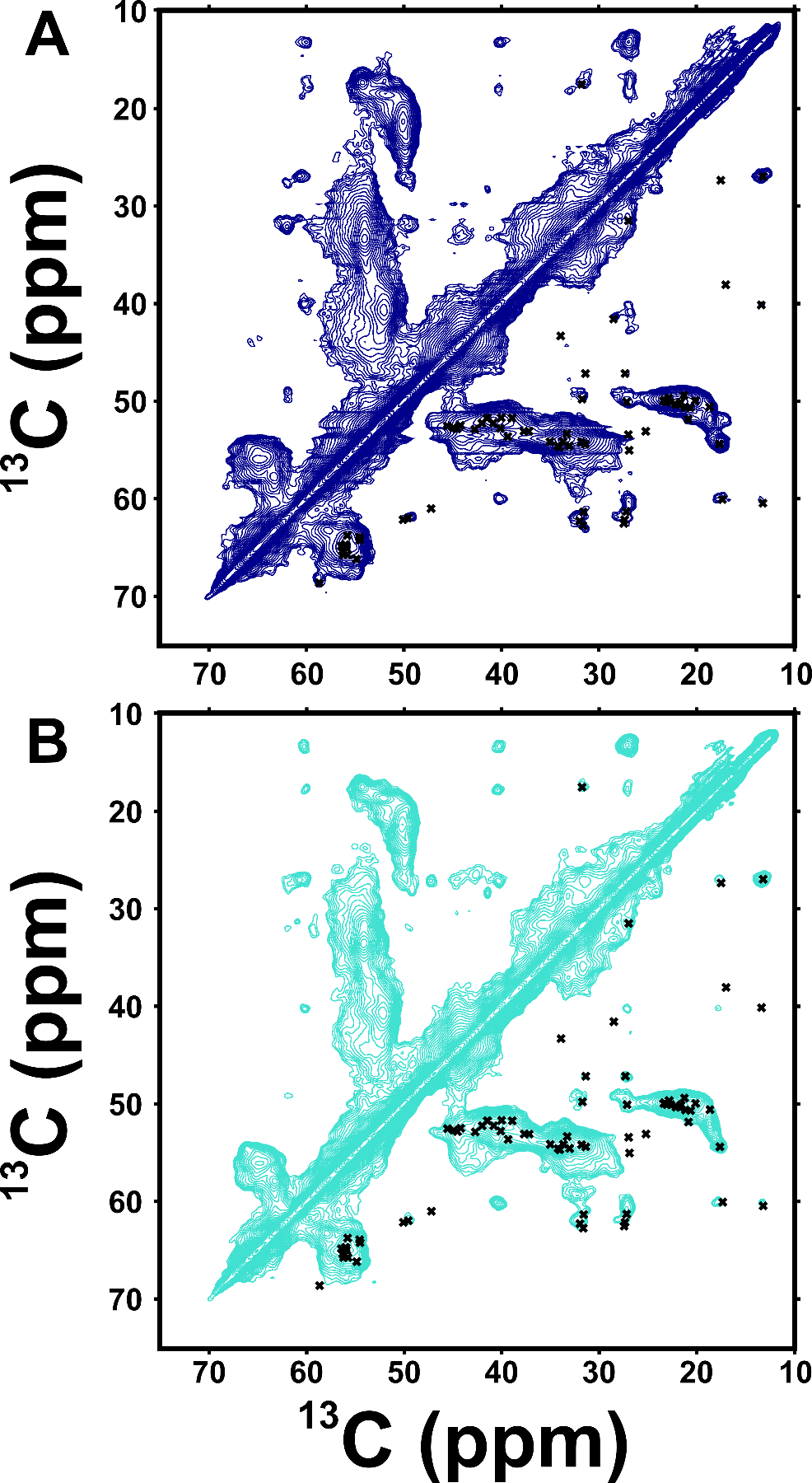
**Supplementary Figure 12. Replicates of P-TDP43LC Fibrils show Same signals.** A) and B) Cross polarization based ^13^C-^13^C DARR solid state NMR spectra with 50ms mixing. X’s are from supplementary figure 6 and denote the same shifts being present for each P-TDP43LC fibril sample.

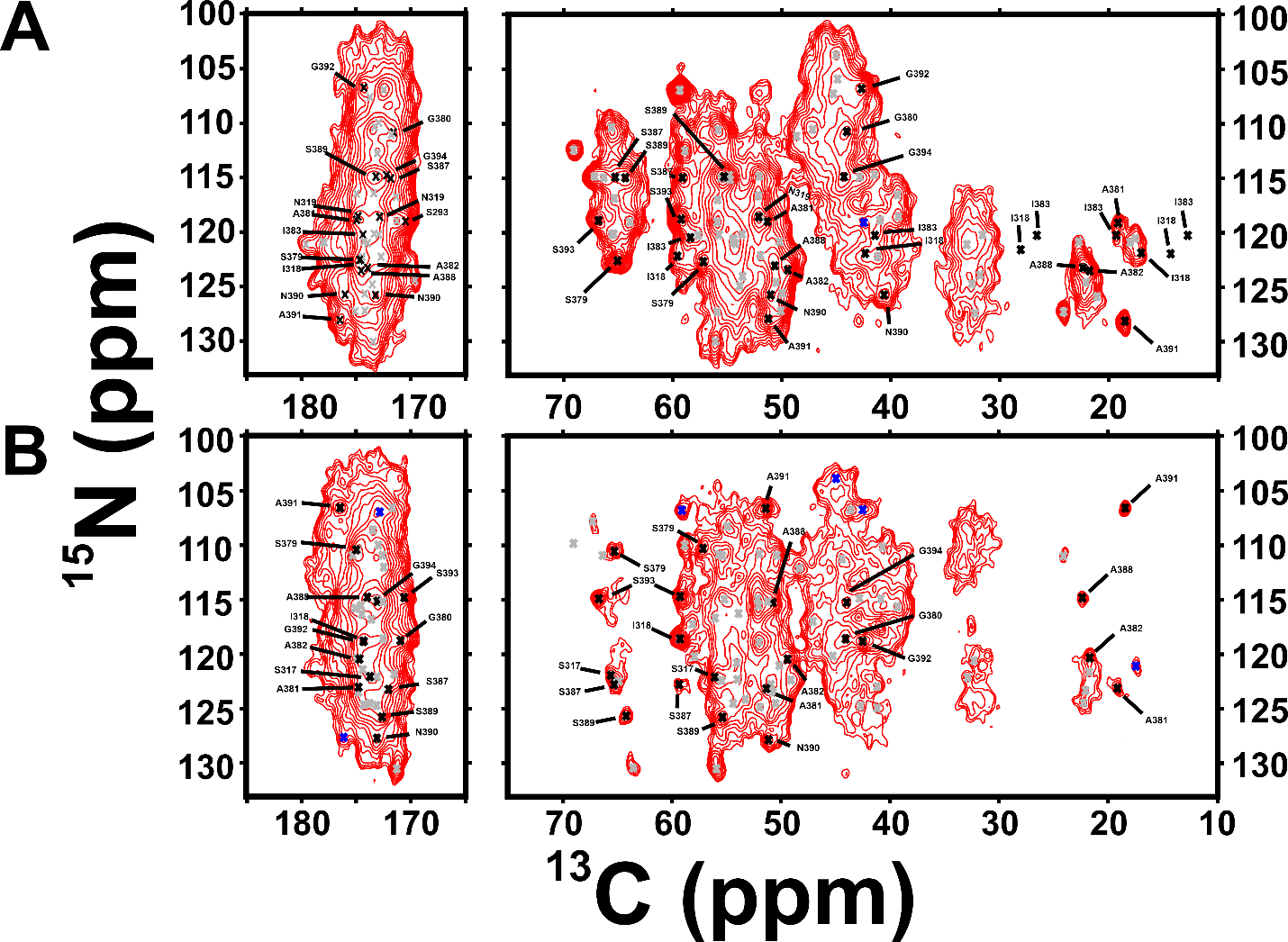
**Supplementary Figure 13. 2D Heteronuclear Solid State NMR on WT-TDP43LC Fibrils with Partial Assignments. A)** Cross polarization based 2D NCACX 50ms DARR mixing experiment shows intra-residue cross peaks. Black x’s represent assigned peaks, grey x’s unassigned peaks, and blue x’s represent weak intensity inter-residue cross peaks. B) Cross polarization based 2D NCOCX 50ms DARR mixing experiment shows inter-residue cross peaks, with the N shift from the “i+1” residue relative to the C shifts. Black x’s represent peak assignments, grey x’s unassigned peaks, and blue x’s intra-residue weak intensity peaks.

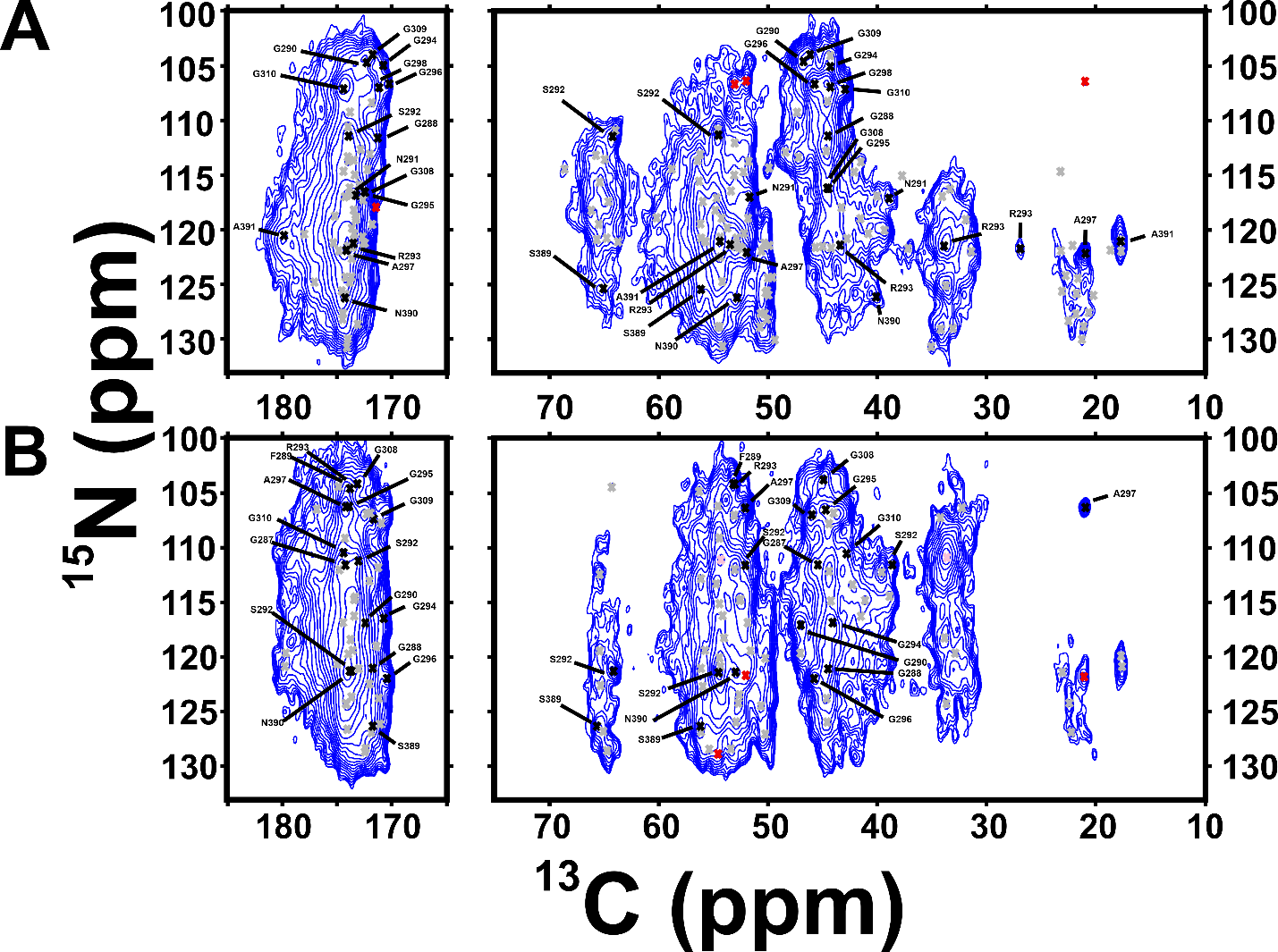
**Supplementary Figure 14. 2D Heteronuclear Solid State NMR on P-TDP43LC Fibrils with Partial Assignments. A)** Cross polarization based 2D NCACX 50ms DARR mixing experiment shows intra-residue cross peaks. Black x’s represent assigned peaks, grey x’s unassigned peaks, and red x’s represent weak intensity inter-residue cross peaks. B) Cross polarization based 2D NCOCX 50ms DARR mixing experiment shows inter-residue cross peaks, with the N shift from the “i+1” residue relative to the C shifts. Black x’s represent peak assignments, grey x’s unassigned peaks, red x’s intra-residue weak intensity peaks, and pink residues the side chain ^15^N ^13^C shifts from Gln residues.

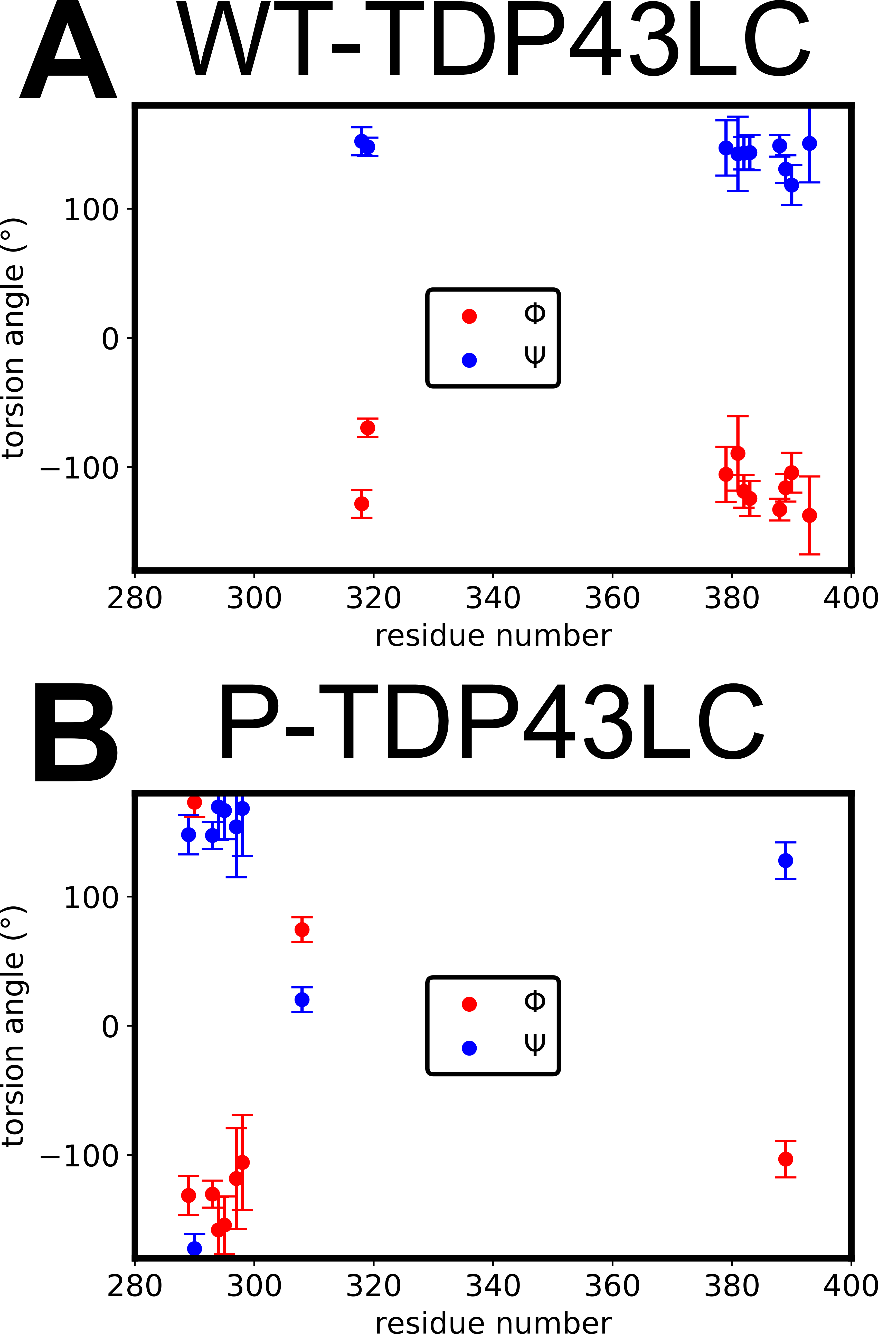
**Supplementary Figure 15. TALOSN Torsion Angle Predictions.** A) “Good” and “Generous” torsion angle predictions from the solid state NMR assignments for WT-TDP43LC fibrils. TALOSN predicts residues A382-I383 and A388-N390 form $\beta$-strands. B) “Good” and “Generous” torsion angle predictions from the solid state NMR assignments for P-TDP43LC fibrils. TALOSN predicts residues R293-G295 forms a $\beta$-strand.

**Supplementary Table 1. Solution NMR assignments for pTDP43LC**. a) S/N comes from the fhsqc spectra, as output by NMRFAM-SPARKY ^17^. b) ‘b’ type refers to overlapped peak intensity in the fhsqc spectra, and ‘g’ refers to good, non-overlapped peak intensity.

| Residue | Residue Type | N | CA | CB | HN | CO | S/N^a^ | S/N Type^b^ |
| --- | --- | --- | --- | --- | --- | --- | --- | --- |
| 258 | G | 1111.1 | 45.54 | 1111.1 | 1111.1 | 174.072 |  |  |
| 259 | A | 123.489 | 52.62 | 19.23 | 7.986 | 177.733 | 66 | b |
| 260 | M | 118.538 | 55.07 | 32.65 | 8.463 | 175.587 | 22 | g |
| 261 | D | 123.024 | 52.32 | 41.07 | 8.184 | 1111.1 | 24 | g |
| 262 | P | 1111.1 | 63.6 | 32.29 | 1111.1 | 177.238 |  |  |
| 263 | K | 119.311 | 56.73 | 32.49 | 8.335 | 1111.1 | 30 | g |
| 264 | H | 1111.1 | 56.27 | 29.24 | 1111.1 | 1111.1 |  |  |
| 265 | N | 118.997 | 53.54 | 38.75 | 8.426 | 175.247 | 61 | b |
| 266 | S | 116.585 | 58.66 | 63.75 | 8.373 | 174.38 | 14 | g |
| 267 | N | 120.535 | 53.63 | 38.7 | 8.454 | 1111.1 | 20 | b |
| 268 | R | 1111.1 | 1111.1 | 1111.1 | 1111.1 | 1111.1 |  |  |
| 269 | Q | 1111.1 | 56.05 | 29.08 | 1111.1 | 176.518 |  |  |
| 270 | L | 122.145 | 55.61 | 42.29 | 8.14 | 177.559 | 24 | b |
| 271 | E | 121.245 | 57.01 | 30.09 | 8.299 | 174.145 | 40 | b |
| 272 | R | 120.216 | 56.35 | 30.64 | 8.026 | 176.369 | 24 | g |
| 273 | S | 116.061 | 58.73 | 63.98 | 8.254 | 175.057 | 32 | g |
| 274 | G | 110.367 | 45.41 | 1111.1 | 8.353 | 174.145 | 60 | b |
| 275 | R | 120.216 | 56.35 | 30.64 | 8.026 | 176.057 | 24 | g |
| 276 | F | 120.437 | 57.61 | 39.34 | 8.282 | 176.757 | 47 | b |
| 277 | G | 110.627 | 45.47 | 1111.1 | 8.263 | 174.302 | 46 | b |
| 278 | G | 107.78 | 45.06 | 1111.1 | 7.897 | 173.307 | 32 | g |
| 279 | N | 118.89 | 51.31 | 38.89 | 8.304 | 1111.1 | 31 | g |
| 280 | P | 1111.1 | 63.77 | 31.69 | 1111.1 | 176.735 |  |  |
| 281 | G | 108.645 | 45.52 | 1111.1 | 8.376 | 174.46 | 45 | b |
| 282 | G | 108.134 | 45.24 | 1111.1 | 8.018 | 173.846 | 38 | b |
| 283 | F | 119.651 | 58.02 | 39.6 | 8.12 | 1111.1 | 62 | b |
| 284 | G | 110.436 | 45.45 | 1111.1 | 8.384 | 173.896 | 45 | b |
| 285 | N | 118.53 | 53.31 | 38.88 | 8.263 | 175.446 | 92 | b |
| 286 | Q | 120.332 | 56.22 | 29.25 | 8.442 | 176.406 | 38 | b |
| 287 | G | 109.273 | 45.46 | 1111.1 | 8.362 | 174.46 | 39 | b |
| 288 | G | 108.134 | 45.24 | 1111.1 | 8.018 | 173.921 | 25 | b |
| 289 | F | 119.585 | 58.03 | 39.59 | 8.135 | 176.402 | 47 | b |
| 290 | G | 110.149 | 45.45 | 1111.1 | 8.385 | 173.896 | 50 | b |
| 291 | N | 118.53 | 53.31 | 38.88 | 8.263 | 175.453 | 92 | b |
| 292 | S | 116.077 | 58.74 | 63.81 | 8.301 | 174.677 | 30 | g |
| 293 | R | 122.331 | 56.27 | 30.61 | 8.355 | 176.749 | 26 | g |
| 294 | G | 109.362 | 45.47 | 1111.1 | 8.311 | 174.646 | 62 | b |
| 295 | G | 108.513 | 45.34 | 1111.1 | 8.263 | 174.64 | 34 | b |
| 296 | G | 108.513 | 45.33 | 1111.1 | 8.263 | 173.928 | 34 | b |
| 297 | A | 123.54 | 52.7 | 19.32 | 8.22 | 178.164 | 47 | g |
| 298 | G | 107.903 | 45.43 | 1111.1 | 8.342 | 174.171 | 37 | g |
| 299 | L | 121.176 | 55.26 | 42.41 | 8.082 | 177.923 | 49 | g |
| 300 | G | 109.036 | 45.5 | 1111.1 | 8.404 | 173.896 | 44 | b |
| 301 | N | 118.53 | 53.23 | 38.77 | 8.263 | 175.082 | 92 | b |
| 302 | N | 118.939 | 53.51 | 38.73 | 8.45 | 175.205 | 34 | b |
| 303 | Q | 120.302 | 56.22 | 29.34 | 8.328 | 176.41 | 47 | b |
| 304 | G | 109.441 | 45.55 | 1111.1 | 8.38 | 174.281 | 44 | b |
| 305 | S | 115.464 | 58.55 | 63.92 | 8.177 | 174.357 | 51 | b |
| 306 | N | 120.353 | 53.49 | 38.63 | 8.484 | 175.261 | 24 | g |
| 307 | M | 120.182 | 55.77 | 32.67 | 8.292 | 176.732 | 41 | b |
| 308 | G | 109.498 | 45.46 | 1111.1 | 8.364 | 174.588 | 47 | b |
| 309 | G | 108.575 | 45.43 | 1111.1 | 8.233 | 174.662 | 48 | b |
| 310 | G | 108.736 | 45.31 | 1111.1 | 8.278 | 174.043 | 41 | b |
| 311 | M | 119.285 | 55.51 | 32.8 | 8.166 | 175.586 | 33 | b |
| 312 | N | 119.311 | 53.06 | 38.84 | 8.304 | 174.756 | 31 | g |
| 313 | F | 120.789 | 58.26 | 39.37 | 8.201 | 176.18 | 26 | b |
| 314 | G | 109.82 | 45.46 | 1111.1 | 8.274 | 173.868 | 26 | g |
| 315 | A | 123.489 | 52.62 | 19.23 | 7.986 | 177.32 | 27 | b |
| 316 | F | 118.064 | 57.5 | 39.51 | 8.018 | 175.505 | 21 | g |
| 317 | S | 116.505 | 58.24 | 64.09 | 8.002 | 173.866 | 20 | g |
| 318 | I | 121.229 | 60.8 | 39.04 | 7.976 | 175.206 | 19 | g |
| 319 | N | 123.212 | 51.19 | 1111.1 | 8.231 | 1111.1 | 17 | g |
| 320 | P | 1111.1 | 64.61 | 1111.1 | 1111.1 | 177.761 |  |  |
| 321 | A | 121.087 | 53.9 | 18.5 | 8.109 | 179.265 | 24 | g |
| 322 | M | 117.983 | 56.46 | 32.92 | 7.935 | 177.363 | 20 | g |
| 323 | M | 120.195 | 56.82 | 1111.1 | 7.948 | 176.958 | 16 | g |
| 324 | A | 123.259 | 53.75 | 18.67 | 8.156 | 179.001 | 17 | g |
| 325 | A | 121.856 | 53.82 | 18.59 | 8.035 | 179.001 | 36 | b |
| 326 | A | 121.856 | 53.82 | 18.59 | 8.035 | 179.004 | 36 | b |
| 327 | Q | 117.995 | 57.41 | 28.75 | 8.047 | 177.107 | 23 | g |
| 328 | A | 122.96 | 53.95 | 18.59 | 8.038 | 178.677 | 12 | g |
| 329 | A | 121.034 | 53.41 | 18.67 | 7.896 | 178.593 | 20 | g |
| 330 | L | 119.721 | 55.98 | 42.14 | 7.823 | 177.927 | 22 | g |
| 331 | Q | 119.21 | 56.06 | 29.1 | 8.045 | 1111.1 | 14 | b |
| 332 | S | 1111.1 | 59.07 | 63.44 | 1111.1 | 174.942 |  |  |
| 333 | S | 117.086 | 58.95 | 63.5 | 8.105 | 174.671 | 17 | g |
| 334 | W | 122.332 | 57.94 | 1111.1 | 8.025 | 177.151 | 23 | g |
| 335 | G | 109.234 | 45.76 | 1111.1 | 8.162 | 174.418 | 15 | g |
| 336 | M | 119.482 | 56.02 | 32.78 | 8.037 | 175.292 | 44 | b |
| 337 | M | 120.304 | 55.76 | 32.74 | 8.275 | 176.935 | 59 | b |
| 338 | G | 108.95 | 45.36 | 1111.1 | 8.247 | 174.418 | 23 | g |
| 339 | M | 119.598 | 56.02 | 32.78 | 8.037 | 176.234 | 32 | b |
| 340 | L | 122.427 | 55.61 | 42.29 | 8.14 | 177.539 | 38 | b |
| 341 | A | 123.945 | 52.98 | 19.1 | 8.185 | 178.091 | 71 | b |
| 342 | S | 113.71 | 58.78 | 63.71 | 8.071 | 174.849 | 22 | g |
| 343 | Q | 121.309 | 56.13 | 29.28 | 8.182 | 176.071 | 27 | g |
| 344 | Q | 120.181 | 56.06 | 29.42 | 8.214 | 175.745 | 32 | b |
| 345 | N | 119.168 | 53.49 | 38.82 | 8.391 | 175.193 | 42 | b |
| 346 | Q | 120.576 | 56.1 | 29.43 | 8.343 | 175.879 | 40 | b |
| 347 | S | 116.547 | 58.52 | 64.09 | 8.327 | 174.452 | 35 | g |
| 348 | G | 110.445 | 44.7 | 1111.1 | 8.177 | 1111.1 | 50 | g |
| 349 | P | 1111.1 | 62.94 | 32.35 | 1111.1 | 177.342 |  |  |
| 350 | S | 115.803 | 58.55 | 63.94 | 8.437 | 175.058 | 34 | g |
| 351 | G | 110.328 | 45.47 | 1111.1 | 8.315 | 173.881 | 39 | g |
| 352 | N | 118.429 | 53.26 | 38.92 | 8.283 | 1111.1 | 43 | b |
| 353 | N | 119.003 | 53.53 | 38.71 | 8.426 | 175.267 | 61 | b |
| 354 | Q | 119.938 | 56.19 | 29.18 | 8.357 | 1111.1 | 74 | b |
| 355 | N | 118.997 | 53.54 | 38.75 | 8.426 | 175.267 | 61 | b |
| 356 | Q | 119.938 | 56.19 | 29.18 | 8.357 | 176.464 | 74 | b |
| 357 | G | 109.203 | 45.5 | 1111.1 | 8.4 | 173.893 | 51 | b |
| 358 | N | 118.391 | 53.31 | 38.88 | 8.262 | 175.261 | 87 | b |
| 359 | M | 120.182 | 55.77 | 32.67 | 8.292 | 176.013 | 41 | b |
| 360 | Q | 121.136 | 55.9 | 29.28 | 8.319 | 175.56 | 44 | g |
| 361 | R | 122.271 | 55.95 | 31.18 | 8.275 | 175.86 | 36 | g |
| 362 | E | 123.111 | 54.45 | 29.61 | 8.41 | 1111.1 | 40 | g |
| 363 | P | 1111.1 | 63.5 | 32.3 | 1111.1 | 176.698 |  |  |
| 364 | N | 117.789 | 53.52 | 38.63 | 8.468 | 175.187 | 39 | g |
| 365 | Q | 120.516 | 56 | 29.5 | 8.2 | 175.481 | 45 | b |
| 366 | A | 124.663 | 52.56 | 19.13 | 8.215 | 177.354 | 47 | g |
| 367 | F | 119.241 | 58.01 | 39.53 | 8.136 | 176.286 | 51 | b |
| 368 | G | 110.559 | 45.46 | 1111.1 | 8.246 | 174.233 | 60 | b |
| 369 | S | 115.492 | 58.59 | 63.94 | 8.228 | 175.068 | 41 | b |
| 370 | G | 110.597 | 45.46 | 1111.1 | 8.455 | 173.892 | 40 | g |
| 371 | N | 118.53 | 53.31 | 38.88 | 8.263 | 174.948 | 92 | b |
| 372 | N | 119.181 | 53.48 | 38.88 | 8.4 | 175.069 | 45 | b |
| 373 | S | 115.663 | 58.59 | 63.86 | 8.187 | 174.054 | 47 | b |
| 374 | Y | 121.949 | 58.13 | 38.74 | 8.181 | 175.812 | 47 | g |
| 375 | S | 117.947 | 58.43 | 63.96 | 8.22 | 174.669 | 35 | g |
| 376 | G | 110.479 | 45.38 | 1111.1 | 7.85 | 173.905 | 39 | g |
| 377 | S | 115.09 | 58.39 | 63.97 | 8.142 | 174.423 | 44 | g |
| 378 | N | 120.713 | 53.41 | 38.9 | 8.528 | 1111.1 | 38 | g |
| 379 | D | 1111.1 | 54.58 | 40.95 | 1111.1 | 1111.1 |  |  |
| 380 | G | 1111.1 | 45.54 | 1111.1 | 1111.1 | 174.072 |  |  |
| 381 | A | 123.306 | 52.62 | 19.23 | 7.979 | 177.478 | 66 | b |
| 382 | A | 122.692 | 52.54 | 19.05 | 8.094 | 177.629 | 53 | g |
| 383 | I | 119.255 | 61.34 | 38.69 | 7.926 | 176.665 | 47 | g |
| 384 | G | 112.109 | 45.34 | 1111.1 | 8.251 | 173.89 | 44 | g |
| 385 | W | 121.08 | 57.56 | 29.63 | 8.022 | 176.757 | 52 | g |
| 386 | G | 110.536 | 45.47 | 1111.1 | 8.262 | 174.076 | 46 | b |
| 387 | S | 115.537 | 58.4 | 63.97 | 8.081 | 174.41 | 47 | g |
| 388 | A | 125.612 | 52.7 | 19.16 | 8.351 | 177.792 | 50 | g |
| 389 | S | 114.419 | 58.51 | 63.81 | 8.187 | 174.35 | 42 | g |
| 390 | N | 120.453 | 53.24 | 38.93 | 8.295 | 174.904 | 45 | b |
| 391 | A | 123.945 | 52.98 | 19.1 | 8.185 | 178.078 | 71 | b |
| 392 | G | 107.871 | 45.37 | 1111.1 | 8.292 | 174.258 | 47 | g |
| 393 | S | 115.294 | 58.53 | 63.93 | 8.168 | 175.062 | 43 | b |
| 394 | G | 110.885 | 45.46 | 1111.1 | 8.462 | 174.101 | 45 | g |
| 395 | S | 115.464 | 58.53 | 63.97 | 8.208 | 1111.1 | 55 | b |
| 396 | G | 110.436 | 45.45 | 1111.1 | 8.384 | 173.81 | 45 | b |
| 397 | F | 120.164 | 58.06 | 39.58 | 8.121 | 175.665 | 37 | g |
| 398 | N | 121.413 | 53.2 | 38.94 | 8.432 | 175.259 | 28 | g |
| 399 | G | 108.486 | 45.59 | 1111.1 | 7.776 | 174.188 | 34 | b |
| 400 | G | 108.221 | 45.16 | 1111.1 | 8.083 | 173.801 | 56 | g |
| 401 | F | 119.513 | 58.02 | 39.6 | 8.116 | 1111.1 | 54 | b |
| 402 | G | 110.436 | 45.45 | 1111.1 | 8.384 | 173.718 | 45 | b |
| 403 | D | 120.307 | 54.49 | 41.29 | 8.194 | 175.958 | 58 | b |
| 404 | D | 119.874 | 54.45 | 40.89 | 8.37 | 176.367 | 60 | b |
| 405 | M | 120.014 | 55.87 | 32.74 | 8.261 | 176.165 | 50 | g |
| 406 | D | 120.797 | 54.61 | 41.21 | 8.303 | 176.173 | 52 | g |
| 407 | S | 115.909 | 58.49 | 63.77 | 8.117 | 174.537 | 47 | g |
| 408 | K | 122.836 | 56.16 | 32.88 | 8.248 | 176.182 | 45 | g |
| 409 | D | 121.168 | 54.36 | 41.28 | 8.236 | 175.902 | 46 | b |
| 410 | D | 120.964 | 54.42 | 41.17 | 8.243 | 1111.1 | 58 | b |
| 411 | G | 1111.1 | 45.54 | 1111.1 | 1111.1 | 174.228 |  |  |
| 412 | W | 120.826 | 57.56 | 29.63 | 8.017 | 176.759 | 49 | g |
| 413 | G | 111.067 | 45.47 | 1111.1 | 8.264 | 173.134 | 44 | b |
| 414 | M | 124.524 | 56.97 | 33.74 | 7.663 | 1111.1 | 45 | g |

**Supplementary Table 2. Unassigned peaks from solution NMR spectra of P-TDP43LC**. a) S/N comes from the fhsqc spectra, as output by NMRFAM-SPARKY ^17^. b) ‘b’ type refers to overlapped peak intensity in the fhsqc spectra, and ‘g’ refers to good, non-overlapped peak intensity.

| N | HN | CA | CB | CA  (i-1) | CB  (i-1) | CO  (i-1) | S/N^a^ | Type^b^ | Notes |
| --- | --- | --- | --- | --- | --- | --- | --- | --- | --- |
| 108.632 | 8.367 | 45.52 | 1111.1 | 54.44 | 40.97 | 177.379 | 63 | b | G380/G411 |
| 109.241 | 8.307 | 45.56 | 1111.1 | 54.56 | 41.05 | 176.749 | 65 | b | G380/G411 |
| 113.984 | 8.279 | 45.49 | 1111.1 | 56.6 | 1111.1 | 176.212 | 9 | g | Unknown |
| 119.938 | 8.357 | 54.45 | 40.88 | 53.59 | 38.66 | 175.257 | 74 | b | D379? |
| 120.453 | 8.239 | 54.7 | 41.03 | 53.87 | 38.84 | 174.919 | 62 | g | D379 |
| 120.561 | 8.326 | 56.17 | 29.16 | 53.4 | 38.6 | 176.391 | 36 | b | Unknown |
| 121.401 | 8.283 | 56.4 | 30.72 | 56.32 | 1111.1 | 176.704 | 26 | b | Unknown |
| 121.039 | 8.209 | 56.74 | 30.45 | 53.43 | 38.63 | 175.38 | 27 | b | Unknown |
| 124.871 | 7.984 | 54.93 | 41.17 | 1111.1 | 1111.1 | 174.617 | 12 | g | Unknown |
| 118.564 | 8.135 | 55.95 | 1111.1 | 1111.1 | 1111.1 | 176.69 | 11 | g | Unknown |
| 119.58 | 8.25 | 59.42 | 1111.1 | 1111.1 | 1111.1 | 176.68 | 13 | g | Unknown |

**Supplementary Table 3. Similarities and Differences in the INEPT-TOBSY Spectra.** The shared peaks are those that appear at nearly identical chemical shifts in the spectra. The unique residues are identified from peaks that are measured in only one of the fibril samples. The number of amino acid residues giving rise to each peak could not be determined as the dynamic residues were unable to be assigned from these data.

| Peaks Shared Between Spectra: Gly, Ile, Lys, Ser, Asn, Glu, Phe, Arg | | |
| --- | --- | --- |
| Motif: | Wild-Type | P-TDP43LC |
| Unique Residues | Ala, Gly, Ile, Ser, Asn | Ala, Asp |

**Supplementary Table 4. Fibril Rigid Residue Relaxation Measurements.** For experimental details, see supplementary table 14-15 and the methods section. The errors from the individual solid-state NMR samples are the scipy output standard deviations for the curve fit relaxation constant. The error reported for the averaged P-TDP43LC sample is not from fitting, but rather the standard deviation across the three independent samples of P-TDP43LC. For wild-type, only one sample had the relaxation measurements made, and the errors are the curve-fitting constants output standard deviation. The CA represents the integrals for relaxation for all non-glycine peaks, ~61 to 48 ppm, and Gly CA signifies the integrals for relaxation fitting from the Glycine region, ~48 to 41 ppm.

| **P-TDP43LC** | | | |
| --- | --- | --- | --- |
| **Sample** | **^1^H T1**$\boldsymbol{\rho}$ **(ms)** | **^15^N ncaT2 CA (ms)** | **^15^N ncaT2 Gly CA (ms)** |
| **P-TDP43LC s3** | 70.5 to 9.9 ppm  1.3 ± 0.1 and 7.8 ± 0.1 | 61.1 to 48.5 ppm  5.94 ± 0.06 | 48.5 to 41 ppm  5.81 ± 0.07 |
| **P-TDP43LC s2** | 70.5 to 11.1 ppm  1.33 ± 0.04 and 9.5 ± 0.08 | 61.8 to 48.7 ppm  6.26 ± 0.04 | 48.4 to 40.9 ppm  6.06 ± 0.06 |
| **P-TDP43LC** s1 | 71.5 to 10.2 ppm  1.66 ± 0.05 and 10.1 ± 0.1 | 61.3 to 48.3 ppm  6.49 ± 0.06 | 48.2 to 41 ppm  6.06 ± 0.08 |
| **Averages** | | | |
| **Average P-TDP43LC** | 1.4 ± 0.2 and 9 ± 1 | 6.2 ± 0.3 | 6.0 ± 0.1 |
| **Wild-Type TDP43LC** | | | |
| **Sample** | **^1^H T1**$\boldsymbol{\rho}$ **(ms)** | **^15^N ncaT2 CA (ms)** | **^15^N ncaT2 Gly CA (ms)** |
| Wild-type TDP43LC | 71.7 to 9.8 ppm  1.16 ± 0.09 and 8.7 ± 0.2 | 61.1 to 47.9 ppm  4.88 ± 0.06 | 47.9 to 41.1 ppm  5.71 ± 0.06 |

**Supplementary Table 5. NMR Experimental Parameters for P-TDP43LC Solution NMR Experiments.** All experiments except HNCACBi used a 17.5 $\mu$M P-TDP43LC concentration, and the HNCACBi used a fresh sample at 22.2 $\mu$M. Spectra were initially referenced according to the D_2_O lock signal via NMRPipe automatic referencing based on the calibrated sample temperature^10^. Spectra were re-referenced in NMRFAM SPARKY to correct for the slight difference in referencing relative to BMRB entry #26823^2,17^.

| **Spectrum** | **Acquisition Parameters^a^** | **Processing Parameters^b^** |
| --- | --- | --- |
| 2D HSQC  (fhsqcf3gpph) | Spectrometer: 800Mhz Bruker AVANCE III; ns = 40; $\nu$_1H-carr_= 4.725 ppm; $\nu$_15N-carr_= 117.029 ppm; $\tau_{t1}$ = 45.1 ms; $\tau$_aq_ = 79.8 ms with 2046 x 512 pts (indirect dim w/ States-TPPI); Temp = 295.8 K, 50% NUS randomly sampled points | ist2D.com standard nmrPipe processing with applying phasing correction and -yFTARG alt.  Re-reference in Sparky:  1H: 0.04 ppm |
| 2D HSQC  (fhsqcf3gpph) | Spectrometer: 800Mhz Bruker AVANCE III; ns = 16; $\nu$_1H-carr_= 4.725 ppm; $\nu$_15N-carr_= 117.029 ppm; $\tau_{t1}$ = 45.1 ms; $\tau$_aq_ = 79.8 ms with 2048 x 256 pts (indirect dim w/ States-TPPI); Temp = 295.8 K | SP_t1_ -off 0.5 -end 1 -pow 1 -c 1  SP_t2_ -off 0.5 -end 1 -pow 1 -c 0.5  Re-reference in Sparky:  1H: 0.04 ppm |
| 3D HNCO  (hncogpwg3d) | Spectrometer: 800Mhz Bruker AVANCE III; ns = 8; $\nu$_1H-carr_= 4.725 ppm; $\nu$_15N-carr_= 117.03 ppm; $\nu$_13C-carr_= 175.66 ppm; $\tau_{t1 N}$ = 12.3 ms; $\tau_{t2 C}$ = 22.7 ms; $\tau$_aq_ = 91.8 ms with 2048 x 60 x 128 pts (H, N, C, indirect w/ States-TPPI); Temp = 295.8 K | SOL  SP_t1_ -off 0.5 -end 1 -pow 1 -c 1  SP_t2_ -off 0.5 -end 1 -pow 1 -c 0.5  SP_t3_ -off 0.5 -end 1 -pow 2 -c 0.5  POLY -auto  Re-reference in Sparky:  15N: 0 ppm  13C: 0ppm  1H: 0.04ppm |
| 3D CBCANH  (hncacbgpwg3d) | Spectrometer: 800Mhz Bruker AVANCE III; ns = 8; $\nu$_1H-carr_= 4.725 ppm; $\nu$_15N-carr_= 117.034 ppm; $\nu$_13C-carr_= 41.691 ppm; $\tau_{t1 N}$ = 12.3 ms; $\tau_{t2 C}$ = 4.3 ms; $\tau$_aq_ = 91.8 ms with 2048 x 60 x 120 pts (H, N, C, indirect dim w/ States-TPPI); Temp = 295.8 K | SOL  SP_t1_ -off 0.5 -end 1 -pow 1 -c 1  SP_t2_ -off 0.5 -end 1 -pow 1 -c 0.5  SP_t3_ -off 0.5 -end 1 -pow 1 -c 0.5  Re-reference in Sparky:  15N: 0 ppm  13C: 0ppm  1H: 0.04ppm |
| 3D CBCA(CO)NH  (gpwg3d) | Spectrometer: 600Mhz Bruker AVANCE III; ns = 8; $\nu$_1H-carr_= 4.718 ppm; $\nu$_15N-carr_= 118.033 ppm; $\nu$_13C-carr_= 41.688 ppm; $\tau_{t1 N}$ = 16.5 ms; $\tau_{t2 C}$ = 6.1 ms; $\tau$_aq_ = 113.6 ms with 2048 x 64 x 128 pts (H, N, C, indirect dim. w/ States-TPPI); Temp = 298.2 K | SOL  SP_t1_ -off 0.5 -end 1 -pow 1 -c 1  SP_t2_ -off 0.5 -end 1 -pow 1 -c 0.5  SP_t3_ -off 0.5 -end 1 -pow 1 -c 0.5  EXT_t3_ -xn 24 (due to loss in FID quality at later times likely due to tuning / topshim spectrometer drift)  Re-reference in Sparky:  15N: 0 ppm  13C: 1.1 ppm  1H: 0.04ppm |
| 3D HNCA  (hncagpwg3d) | Spectrometer: 600Mhz Bruker AVANCE III; ns = 16; $\nu$_1H-carr_= 4.718 ppm; $\nu$_15N-carr_= 117.033 ppm; $\nu$_13C-carr_= 55.888 ppm; $\tau_{t1 N}$ = 15.0 ms; $\tau_{t2 C}$ = 14.1 ms; $\tau$_aq_ = 121.7 ms with 2048 x 64 x 128 pts (H, N, C, indirect dim. w/ States-TPPI); Temp = 298.2 K | SOL  SP_t1_ -off 0.5 -end 1 -pow 1 -c 1  SP_t2_ -off 0.5 -end 1 -pow 1 -c 0.5  SP_t3_ -off 0.5 -end 1 -pow 1 -c 0.5  Re-reference in Sparky:  15N: 0ppm  13C: 0ppm  1H: 0.04ppm |
| 3D iCBCANH  (HNCACBigpwg3d) | Spectrometer: 800Mhz Bruker AVANCE III; ns = 16; $\nu$_1H-carr_= 4.787 ppm; $\nu$_15N-carr_= 118.093 ppm; $\nu$_13C-carr_= 45.750 ppm; $\tau_{t1 N}$ = 12.3 ms; $\tau_{t2 C}$ = 8.5 ms; $\tau$_aq_ = 91.8 ms with 2048 x 60 x 240 pts (H, N, C, indirect dim. w/ States-TPPI); Temp = 296.5 K, 50% NUS randomly sampled points | ist3D.com standard nmrPipe processing script with applying phasing corrections and -yFTARG alt & neg, as well as -zFTARG alt.  Re-reference in Sparky:  15N: -0.05ppm  13C: 0ppm  1H: -0.023ppm |

^a^ ns is the number of scans averaged; $\tau_{aq}$ is the total acquisition time in the directly detected dimension; $\nu$_X-carr_ is the center frequency; $\tau_{tx}$ is the total acquisition time for the specified dimension. The proton pulse length was optimized and used from the value given by the bruker “pulsecal” command with a 1d proton experiment, zggpwg.

**Supplementary Table 6. NMR Assignments for WT-TDP43LC.** See methods for assignment procedure. S/N*^a^* corresponds to the value measured by NMRFAM-SPARKY for the corresponding 3D CANCO signal^17^. The * denotes overlapped peak intensity, and the “b” corresponds to residues without a unique assignment from the CANCO spectra specifically, but with signals from the NCACX / NCOCX type spectra.

| **Residue** | **Type** | **N** | **CA** | **CO** | **CB** | **CG** | **CD** | **S/N^a^** |
| --- | --- | --- | --- | --- | --- | --- | --- | --- |
| **317** | **S** |  | 56.124 | 173.547 | 65.657 |  |  | b |
| **318** | **I** | 122.037 | 59.329 | 174.649 | 42.365 | 28.055 | 16.81 | 15 |
| **319** | **N** | 118.5 | 52 | 173.5 |  | 175.119 |  | b |
| **378** | **N** |  |  | 173 |  |  |  | b |
| **379** | **S** | 122.607 | 57.134 | 174.717 | 65.126 |  |  | 22 |
| **380** | **G** | 110.794 | 43.959 | 171.452 |  |  |  | 18 |
| **381** | **A** | 118.987 | 51.321 | 174.901 | 19.151 |  |  | 20^*^ |
| **382** | **A** | 123.467 | 49.365 | 174.624 | 21.773 |  |  | 25 |
| **383** | **I** | 120.368 | 58.568 | 174.398 | 41.496 | 26.437 | 19.324 | 17 |
| **387** | **S** | 115.064 | 59.058 | 171.783 | 65.255 |  |  | b |
| **388** | **A** | 123.191 | 50.583 | 173.988 | 22.361 |  |  | b |
| **389** | **S** | 114.831 | 55.316 | 173.109 | 64.313 |  |  | 32^*^ |
| **390** | **N** | 125.813 | 51.029 | 173.213 | 40.447 | 176.059 |  | b |
| **391** | **A** | 128.041 | 51.269 | 176.46 | 18.514 |  |  | b |
| **392** | **G** | 106.723 | 42.467 | 174.28 |  |  |  | 14 |
| **393** | **S** | 118.894 | 59.225 | 170.524 | 66.8 |  |  | 19 |
| **394** | **G** | 114.859 | 44.372 | 172.179 |  |  |  | 13 |
| **395** | **S** | 115.2 |  |  |  |  |  | b |

**Supplementary Tables 7. MCASSIGN 2b tables for WT-TDP43LC Fibrils.** A) 2D/3D NCACX and CC 50ms DARR table, B) 2D/3D NCOCX, C) 3D CANCO.

| **Observed Chemical Shift (ppm)** | | | | | | **Chemical Shift Uncertainty (ppm)** | | | | | |  |  |
| --- | --- | --- | --- | --- | --- | --- | --- | --- | --- | --- | --- | --- | --- |
| **^15^N** | **^13^CA** | **^13^CO** | **^13^CB** | **^13^CG** | **^13^CD**  **/CG2** | **^15^N** | **^13^CA** | **^13^CO** | **^13^CB** | **^13^CG** | **^13^CD**  **/CG2** | **MP^a^** | **Residue Type** |
| 112.609 | 58.9 | 173.021 | 69.088 | 1111.1 | 1111.1 | 0.4 | 0.4 | 1 | 0.4 | 0.4 | 0.4 | 1 | S |
| 118.894 | 59.225 | 170.524 | 66.8 | 1111.1 | 1111.1 | 0.4 | 0.4 | 1 | 0.4 | 0.4 | 0.4 | 1 | S |
| 118.987 | 51.321 | 174.901 | 19.151 | 1111.1 | 1111.1 | 0.4 | 0.4 | 1 | 0.4 | 0.4 | 0.4 | 1 | A |
| 128.041 | 51.269 | 176.46 | 18.514 | 1111.1 | 1111.1 | 0.4 | 0.4 | 1 | 0.4 | 0.4 | 0.4 | 1 | A |
| 127.223 | 50.19 | 175.057 | 24.073 | 1111.1 | 1111.1 | 0.4 | 0.4 | 1 | 0.4 | 0.4 | 0.4 | 1 | A |
| 106.961 | 59.301 | 172.446 | 1111.1 | 1111.1 | 1111.1 | 0.4 | 0.4 | 1 | 0.4 | 0.4 | 0.4 | 1 | HY |
| 122.037 | 59.329 | 174.649 | 42.365 | 28.055 | 16.81 | 0.4 | 0.4 | 1 | 0.4 | 0.4 | 0.4 | 1 | I |
| 120.368 | 58.568 | 174.398 | 41.496 | 26.437 | 19.324 | 0.4 | 0.4 | 1 | 0.4 | 0.4 | 0.4 | 1 | I |
| 118.535 | 52.065 | 173.401 | 39.283 | 175.119 | 1111.1 | 0.4 | 0.4 | 1 | 0.4 | 0.4 | 0.4 | 1 | DN |
| 122.269 | 52.035 | 172.929 | 41.123 | 174.511 | 1111.1 | 0.4 | 0.4 | 1 | 0.4 | 0.4 | 0.4 | 1 | DN |
| 106.723 | 42.467 | 174.28 | 1111.1 | 1111.1 | 1111.1 | 0.4 | 0.4 | 1 | 0.4 | 0.4 | 0.4 | 1 | G |
| 103.513 | 45.001 | 170.665 | 1111.1 | 1111.1 | 1111.1 | 0.4 | 0.4 | 1 | 0.4 | 0.4 | 0.4 | 1 | G |
| 110.367 | 55.77 | 173.311 | 65.543 | 1111.1 | 1111.1 | 0.4 | 0.4 | 1 | 0.4 | 0.4 | 0.4 | 1 | S |
| 122.607 | 57.134 | 174.717 | 65.126 | 1111.1 | 1111.1 | 0.4 | 0.4 | 1 | 0.4 | 0.4 | 0.4 | 1 | S |
| 118.882 | 55.815 | 171.346 | 63.893 | 1111.1 | 1111.1 | 0.4 | 0.4 | 1 | 0.4 | 0.4 | 0.4 | 1 | S |
| 110.189 | 47.293 | 173.092 | 1111.1 | 1111.1 | 1111.1 | 0.4 | 0.4 | 1 | 0.4 | 0.4 | 0.4 | 1 | G |
| 115.064 | 59.058 | 171.783 | 65.255 | 1111.1 | 1111.1 | 0.4 | 0.4 | 1 | 0.4 | 0.4 | 0.4 | 1 | S |
| 114.832 | 55.316 | 173.109 | 64.313 | 1111.1 | 1111.1 | 0.4 | 0.4 | 1 | 0.4 | 0.4 | 0.4 | 1 | S |
| 114.808 | 54.862 | 173.292 | 67.209 | 1111.1 | 1111.1 | 0.4 | 0.4 | 1 | 0.4 | 0.4 | 0.4 | 1 | S |
| 114.968 | 55.527 | 173.218 | 66.269 | 1111.1 | 1111.1 | 0.4 | 0.4 | 1 | 0.4 | 0.4 | 0.4 | 1 | S |
| 120.104 | 55.722 | 173.319 | 65.672 | 1111.1 | 1111.1 | 0.4 | 0.4 | 1 | 0.4 | 0.4 | 0.4 | 1 | S |
| 120.124 | 57.737 | 172.927 | 65.398 | 1111.1 | 1111.1 | 0.4 | 0.4 | 1 | 0.4 | 0.4 | 0.4 | 1 | S |
| 116.586 | 52.083 | 173.447 | 39.429 | 174.978 | 1111.1 | 0.4 | 0.4 | 1 | 0.4 | 0.4 | 0.4 | 1 | DN |
| 124.539 | 50.401 | 169.6 | 22.065 | 1111.1 | 1111.1 | 0.4 | 0.4 | 1 | 0.4 | 0.4 | 0.4 | 1 | A |
| 125.813 | 51.029 | 173.213 | 40.447 | 176.059 | 1111.1 | 0.4 | 0.4 | 1 | 0.4 | 0.4 | 0.4 | 1 | DN |
| 120.675 | 54.411 | 179.817 | 17.595 | 1111.1 | 1111.1 | 0.4 | 0.4 | 1 | 0.4 | 0.4 | 0.4 | 1 | A |
| 120.86 | 54.399 | 178.006 | 18.102 | 1111.1 | 1111.1 | 0.4 | 0.4 | 1 | 0.4 | 0.4 | 0.4 | 1 | A |
| 120.862 | 50.167 | 174.179 | 22.798 | 1111.1 | 1111.1 | 0.4 | 0.4 | 1 | 0.4 | 0.4 | 0.4 | 1 | A |
| 123.467 | 49.365 | 174.624 | 21.773 | 1111.1 | 1111.1 | 0.4 | 0.4 | 1 | 0.4 | 0.4 | 0.4 | 1 | A |
| 123.191 | 50.583 | 173.988 | 22.361 | 1111.1 | 1111.1 | 0.4 | 0.4 | 1 | 0.4 | 0.4 | 0.4 | 1 | A |
| 125.688 | 49.816 | 174.178 | 21.062 | 1111.1 | 1111.1 | 0.4 | 0.4 | 1 | 0.4 | 0.4 | 0.4 | 1 | A |
| 130.137 | 56.032 | 173.303 | 1111.1 | 1111.1 | 1111.1 | 0.4 | 0.4 | 1 | 0.4 | 0.4 | 0.4 | 1 | RDEQHLKMFWY |
| 121.046 | 54.159 | 173.856 | 32.943 | 1111.1 | 177.846 | 0.4 | 0.4 | 1 | 0.4 | 0.4 | 0.4 | 1 | EQ |
| 124.826 | 53.95 | 173.531 | 32.53 | 1111.1 | 1111.1 | 0.4 | 0.4 | 1 | 0.4 | 0.4 | 0.4 | 1 | RCEQHKMW |
| 123.959 | 53.614 | 173.547 | 31.806 | 1111.1 | 1111.1 | 0.4 | 0.4 | 1 | 0.4 | 0.4 | 0.4 | 1 | RCEQHKMW |
| 120.174 | 53.598 | 173.394 | 31.557 | 1111.1 | 1111.1 | 0.4 | 0.4 | 1 | 0.4 | 0.4 | 0.4 | 1 | RCEQHKMW |
| 114.799 | 52.046 | 172.925 | 41.297 | 1111.1 | 1111.1 | 0.4 | 0.4 | 1 | 0.4 | 0.4 | 0.4 | 1 | DNCLFY |
| 114.859 | 44.372 | 172.179 | 1111.1 | 1111.1 | 1111.1 | 0.4 | 0.4 | 1 | 0.4 | 0.4 | 0.4 | 1 | G |
| 127.26 | 55.855 | 174.137 | 32.345 | 1111.1 | 1111.1 | 0.4 | 0.4 | 1 | 0.4 | 0.4 | 0.4 | 1 | RCEQHKMW |
| 110.794 | 43.959 | 171.452 | 1111.1 | 1111.1 | 1111.1 | 0.4 | 0.4 | 1 | 0.4 | 0.4 | 0.4 | 1 | G |
| 105.894 | 44.91 | 172.754 | 1111.1 | 1111.1 | 1111.1 | 0.4 | 0.4 | 1 | 0.4 | 0.4 | 0.4 | 1 | G |
| 107.454 | 45.338 | 173.73 | 1111.1 | 1111.1 | 1111.1 | 0.4 | 0.4 | 1 | 0.4 | 0.4 | 0.4 | 1 | G |
| 115.011 | 54.77 | 173.167 | 43.079 | 1111.1 | 1111.1 | 0.4 | 0.4 | 1 | 0.4 | 0.4 | 0.4 | 1 | DNCLFY |
| 116.82 | 56.017 | 171.219 | 65.313 | 1111.1 | 1111.1 | 0.4 | 0.4 | 1 | 0.4 | 0.4 | 0.4 | 1 | S |
| 111.06 | 48.656 | 171.715 | 1111.1 | 1111.1 | 1111.1 | 0.4 | 0.4 | 1 | 0.4 | 0.4 | 0.4 | 1 | G |
| 118.556 | 52.011 | 173.646 | 41.281 | 175.156 | 1111.1 | 0.4 | 0.4 | 1 | 0.4 | 0.4 | 0.4 | 1 | DN |
| ^a^multiplicity (MP), the number of times a signal can be assigned | | | | | | | | | | | | | |

B)

| **Observed Chemical Shift (ppm)** | | | | | | **Chemical Shift Uncertainty (ppm)** | | | | | |  |  |
| --- | --- | --- | --- | --- | --- | --- | --- | --- | --- | --- | --- | --- | --- |
| **^15^N** | **^13^CA** | **^13^CO** | **^13^CB** | **^13^CG** | **^13^CD** | **^15^N** | **^13^CA** | **^13^CO** | **^13^CB** | **^13^CG** | **^13^CD** | **MP^a^** | **Residue Type** |
| 109.952 | 59.016 | 173.016 | 69.192 | 1111.1 | 1111.1 | 0.4 | 0.4 | 0.4 | 0.4 | 0.4 | 0.4 | 1 | S |
| 114.719 | 59.321 | 170.569 | 66.807 | 1111.1 | 1111.1 | 0.4 | 0.4 | 0.4 | 0.4 | 0.4 | 0.4 | 1 | S |
| 123.067 | 51.372 | 174.863 | 19.2 | 1111.1 | 1111.1 | 0.4 | 0.4 | 0.4 | 0.4 | 0.4 | 0.4 | 1 | A |
| 106.567 | 51.517 | 176.497 | 18.514 | 1111.1 | 1111.1 | 0.4 | 0.4 | 0.4 | 0.4 | 0.4 | 0.4 | 1 | A |
| 110.979 | 50.468 | 175.233 | 24.185 | 1111.1 | 1111.1 | 0.4 | 0.4 | 0.4 | 0.4 | 0.4 | 0.4 | 1 | A |
| 118.548 | 59.409 | 172.557 | 1111.1 | 1111.1 | 1111.1 | 0.4 | 0.4 | 0.4 | 0.4 | 0.4 | 0.4 | 1 | HSY |
| 130.541 | 55.948 | 171.33 | 63.68 | 1111.1 | 1111.1 | 0.4 | 0.4 | 0.4 | 0.4 | 0.4 | 0.4 | 1 | S |
| 125.746 | 55.486 | 172.619 | 64.255 | 1111.1 | 1111.1 | 0.4 | 0.4 | 0.4 | 0.4 | 0.4 | 0.4 | 1 | S |
| 122.433 | 55.441 | 172.132 | 64.84 | 1111.1 | 1111.1 | 0.4 | 0.4 | 0.4 | 0.4 | 0.4 | 0.4 | 1 | S |
| 122.129 | 56.124 | 173.547 | 65.657 | 1111.1 | 1111.1 | 0.4 | 0.4 | 0.4 | 0.4 | 0.4 | 0.4 | 1 | S |
| 110.415 | 57.267 | 174.973 | 65.263 | 1111.1 | 1111.1 | 0.4 | 0.4 | 0.4 | 0.4 | 0.4 | 0.4 | 1 | S |
| 108.213 | 54.929 | 173.568 | 67.337 | 1111.1 | 1111.1 | 0.4 | 0.4 | 0.4 | 0.4 | 0.4 | 0.4 | 1 | S |
| 120.395 | 49.448 | 174.79 | 21.832 | 1111.1 | 1111.1 | 0.4 | 0.4 | 0.4 | 0.4 | 0.4 | 0.4 | 1 | A |
| 121.349 | 50.124 | 174.408 | 21.711 | 1111.1 | 1111.1 | 0.4 | 0.4 | 0.4 | 0.4 | 0.4 | 0.4 | 1 | A |
| 123.465 | 50.65 | 174.745 | 22.032 | 1111.1 | 1111.1 | 0.4 | 0.4 | 0.4 | 0.4 | 0.4 | 0.4 | 1 | A |
| 124.562 | 50.57 | 174.355 | 22.22 | 1111.1 | 1111.1 | 0.4 | 0.4 | 0.4 | 0.4 | 0.4 | 0.4 | 1 | A |
| 116.87 | 47.114 | 173.496 | 1111.1 | 1111.1 | 1111.1 | 0.4 | 0.4 | 0.4 | 0.4 | 0.4 | 0.4 | 1 | G |
| 118.844 | 42.553 | 174.383 | 1111.1 | 1111.1 | 1111.1 | 0.4 | 0.4 | 0.4 | 0.4 | 0.4 | 0.4 | 1 | G |
| 115.665 | 52.2 | 175.213 | 39.335 | 1111.1 | 1111.1 | 0.4 | 0.4 | 0.4 | 0.4 | 0.4 | 0.4 | 1 | DNCLMFY |
| 111.018 | 55.472 | 174.086 | 66.428 | 1111.1 | 1111.1 | 0.4 | 0.4 | 0.4 | 0.4 | 0.4 | 0.4 | 1 | S |
| 122.965 | 59.284 | 172.07 | 65.298 | 1111.1 | 1111.1 | 0.4 | 0.4 | 0.4 | 0.4 | 0.4 | 0.4 | 1 | S |
| 106.644 | 43.785 | 171.682 | 1111.1 | 1111.1 | 1111.1 | 0.4 | 0.4 | 0.4 | 0.4 | 0.4 | 0.4 | 1 | G |
| 115.33 | 54.966 | 174.595 | 40.866 | 1111.1 | 1111.1 | 0.4 | 0.4 | 0.4 | 0.4 | 0.4 | 0.4 | 1 | DNCILMFY |
| 114.922 | 52.194 | 173.032 | 1111.1 | 175.397 | 1111.1 | 0.4 | 0.4 | 0.4 | 0.4 | 0.4 | 0.4 | 1 | DN |
| 118.861 | 52.139 | 174.487 | 1111.1 | 1111.1 | 1111.1 | 0.4 | 0.4 | 0.4 | 0.4 | 0.4 | 0.4 | 1 | ARDNCEQHLKMFWY |
| 117.3 | 58.322 | 173.186 | 1111.1 | 1111.1 | 1111.1 | 0.4 | 0.4 | 0.4 | 0.4 | 0.4 | 0.4 | 1 | RDNCEQHILKMFPSTWYV |
| 112.227 | 48.36 | 172.454 | 1111.1 | 1111.1 | 1111.1 | 0.4 | 0.4 | 0.4 | 0.4 | 0.4 | 0.4 | 1 | ANG |
| 122.205 | 54.007 | 173.77 | 32.887 | 1111.1 | 1111.1 | 0.4 | 0.4 | 0.4 | 0.4 | 0.4 | 0.4 | 1 | RCEQHIKMWY |
| 118.575 | 44.294 | 170.716 | 1111.1 | 1111.1 | 1111.1 | 0.4 | 0.4 | 0.4 | 0.4 | 0.4 | 0.4 | 1 | G |
| 114.91 | 50.812 | 174.036 | 22.384 | 1111.1 | 1111.1 | 0.4 | 0.4 | 0.4 | 0.4 | 0.4 | 0.4 | 1 | A |
| 120.136 | 45.212 | 172.162 | 1111.1 | 1111.1 | 1111.1 | 0.4 | 0.4 | 0.4 | 0.4 | 0.4 | 0.4 | 1 | G |
| 115.248 | 43.953 | 173.026 | 1111.1 | 1111.1 | 1111.1 | 0.4 | 0.4 | 0.4 | 0.4 | 0.4 | 0.4 | 1 | G |
| 115.106 | 47.045 | 172.45 | 1111.1 | 1111.1 | 1111.1 | 0.4 | 0.4 | 0.4 | 0.4 | 0.4 | 0.4 | 1 | G |
| 111.004 | 55.698 | 172.595 | 44.422 | 1111.1 | 1111.1 | 0.4 | 0.4 | 0.4 | 0.4 | 0.4 | 0.4 | 1 | DNCILFY |
| 120.343 | 58.153 | 174.887 | 1111.1 | 1111.1 | 1111.1 | 0.4 | 0.4 | 0.4 | 0.4 | 0.4 | 0.4 | 1 | DNCILFY |
| 118.71 | 59.352 | 174.269 | 1111.1 | 1111.1 | 1111.1 | 0.4 | 0.4 | 0.4 | 0.4 | 0.4 | 0.4 | 1 | DCILFY |
| 127.873 | 51.261 | 173.13 | 1111.1 | 1111.1 | 1111.1 | 0.4 | 0.4 | 0.4 | 0.4 | 0.4 | 0.4 | 1 | ARDNCEQHLKMFWY |
| 124.905 | 52.07 | 173.077 | 41.106 | 174.358 | 1111.1 | 0.4 | 0.4 | 0.4 | 0.4 | 0.4 | 0.4 | 1 | DN |
| 110.291 | 52.017 | 174.834 | 40.658 | 1111.1 | 1111.1 | 0.4 | 0.4 | 0.4 | 0.4 | 0.4 | 0.4 | 1 | DNLMFY |
| 116.373 | 53.888 | 174.539 | 1111.1 | 1111.1 | 1111.1 | 0.4 | 0.4 | 0.4 | 0.4 | 0.4 | 0.4 | 1 | ARDNCEQHLKMFSWY |
| 120.576 | 54.094 | 173.856 | 32.52 | 1111.1 | 1111.1 | 0.4 | 0.4 | 0.4 | 0.4 | 0.4 | 0.4 | 1 | RCEQHIKMW |
| 122.111 | 48.945 | 171.481 | 1111.1 | 1111.1 | 1111.1 | 0.4 | 0.4 | 0.4 | 0.4 | 0.4 | 0.4 | 1 | ADNGH |
| 124.327 | 53.29 | 174.629 | 42.71 | 1111.1 | 1111.1 | 0.4 | 0.4 | 0.4 | 0.4 | 0.4 | 0.4 | 1 | DNCILFY |
| 121.405 | 55.319 | 173.564 | 1111.1 | 1111.1 | 1111.1 | 0.4 | 0.4 | 0.4 | 0.4 | 0.4 | 0.4 | 1 | ARDNCEQHILKMFSWY |
| 122.406 | 51.026 | 173.149 | 41.091 | 1111.1 | 1111.1 | 0.4 | 0.4 | 0.4 | 0.4 | 0.4 | 0.4 | 1 | DNLMFY |
| ^a^multiplicity (MP), the number of times a signal can be assigned | | | | | | | | | | | | | |

C)

| **Observed Chemical Shift (ppm)** | | | | | | **Chemical Shift Uncertainty (ppm)** | | | | | |  |  |
| --- | --- | --- | --- | --- | --- | --- | --- | --- | --- | --- | --- | --- | --- |
| **^15^N** | **^13^CA** | **^13^CO** | **^13^CB** | **^13^CG** | **^13^CD** | **^15^N** | **^13^CA** | **^13^CO** | **^13^CB** | **^13^CG** | **^13^CD** | **MP^a^** | **Residue Type** |
| 124.8 | 54.134 | 173.906 | 1111.1 | 1111.1 | 1111.1 | 0.5 | 0.5 | 0.5 | 0.5 | 0.5 | 0.5 | 1 | ARDNEQHILKMFSWY |
| 110.983 | 54.313 | 172.996 | 1111.1 | 1111.1 | 1111.1 | 0.5 | 0.5 | 0.5 | 0.5 | 0.5 | 0.5 | 1 | RDNEQHMFSWY |
| 110.983 | 54.313 | 172.996 | 1111.1 | 1111.1 | 1111.1 | 0.5 | 0.5 | 0.5 | 0.5 | 0.5 | 0.5 | 1 | RDNEQHMFSWY |
| 113.148 | 46.829 | 175.205 | 1111.1 | 1111.1 | 1111.1 | 0.5 | 0.5 | 0.5 | 0.5 | 0.5 | 0.5 | 1 | G |
| 113.7 | 52.311 | 173.639 | 1111.1 | 1111.1 | 1111.1 | 0.5 | 0.5 | 0.5 | 0.5 | 0.5 | 0.5 | 1 | RDNEQHLKMFSWY |
| 113.7 | 56.419 | 174.241 | 1111.1 | 1111.1 | 1111.1 | 0.5 | 0.5 | 0.5 | 0.5 | 0.5 | 0.5 | 1 | RDNEQHILKMFSTWY |
| 116.2 | 44.504 | 173.459 | 1111.1 | 1111.1 | 1111.1 | 0.5 | 0.5 | 0.5 | 0.5 | 0.5 | 0.5 | 1 | G |
| 114.776 | 52.651 | 173.379 | 1111.1 | 1111.1 | 1111.1 | 0.5 | 0.5 | 0.5 | 0.5 | 0.5 | 0.5 | 1 | ARDNEQHLKMFSWY |
| 115.6 | 56.174 | 174.225 | 1111.1 | 1111.1 | 1111.1 | 0.5 | 0.5 | 0.5 | 0.5 | 0.5 | 0.5 | 1 | RDNEQHILKMFSTWY |
| 117.206 | 54.588 | 173.976 | 1111.1 | 1111.1 | 1111.1 | 0.5 | 0.5 | 0.5 | 0.5 | 0.5 | 0.5 | 1 | ARDNEQHILKMFSWY |
| 116.938 | 51.738 | 172.862 | 1111.1 | 1111.1 | 1111.1 | 0.5 | 0.5 | 0.5 | 0.5 | 0.5 | 0.5 | 1 | ARDNEQHLKMFWY |
| 116.938 | 51.738 | 172.862 | 1111.1 | 1111.1 | 1111.1 | 0.5 | 0.5 | 0.5 | 0.5 | 0.5 | 0.5 | 1 | ARDNEQHLKMFWY |
| 128.732 | 54.148 | 172.62 | 1111.1 | 1111.1 | 1111.1 | 0.5 | 0.5 | 0.5 | 0.5 | 0.5 | 0.5 | 1 | ARDNEQHILKMFWY |
| 128.732 | 54.119 | 172.648 | 1111.1 | 1111.1 | 1111.1 | 0.5 | 0.5 | 0.5 | 0.5 | 0.5 | 0.5 | 1 | ARDNEQHILKMFWY |
| 126.487 | 50.16 | 173.761 | 1111.1 | 1111.1 | 1111.1 | 0.5 | 0.5 | 0.5 | 0.5 | 0.5 | 0.5 | 1 | ADNEQHLM |
| 126.487 | 50.16 | 173.761 | 1111.1 | 1111.1 | 1111.1 | 0.5 | 0.5 | 0.5 | 0.5 | 0.5 | 0.5 | 1 | ADNEQHLM |
| 121.6 | 52.943 | 173.846 | 1111.1 | 1111.1 | 1111.1 | 0.5 | 0.5 | 0.5 | 0.5 | 0.5 | 0.5 | 1 | ARDNEQHLKMFSWY |
| 116.3 | 53.374 | 172.998 | 1111.1 | 1111.1 | 1111.1 | 0.5 | 0.5 | 0.5 | 0.5 | 0.5 | 0.5 | 1 | ARDNEQHLKMFSWY |
| 117.4 | 56.032 | 174.277 | 1111.1 | 1111.1 | 1111.1 | 0.5 | 0.5 | 0.5 | 0.5 | 0.5 | 0.5 | 1 | RDNEQHILKMFSTWY |
| 118.808 | 43.132 | 172.859 | 1111.1 | 1111.1 | 1111.1 | 0.5 | 0.5 | 0.5 | 0.5 | 0.5 | 0.5 | 1 | G |
| 112.7 | 44.646 | 174.189 | 1111.1 | 1111.1 | 1111.1 | 0.5 | 0.5 | 0.5 | 0.5 | 0.5 | 0.5 | 1 | G |
| 108.4 | 44.317 | 174.068 | 1111.1 | 1111.1 | 1111.1 | 0.5 | 0.5 | 0.5 | 0.5 | 0.5 | 0.5 | 1 | G |
| 125.013 | 56.092 | 174.432 | 1111.1 | 1111.1 | 1111.1 | 0.5 | 0.5 | 0.5 | 0.5 | 0.5 | 0.5 | 1 | RDNEQHILKMFSTWY |
| 128.9 | 50.419 | 173.73 | 1111.1 | 1111.1 | 1111.1 | 0.5 | 0.5 | 0.5 | 0.5 | 0.5 | 0.5 | 1 | ARDNEQHLKM |
| 129.709 | 49.609 | 171.167 | 1111.1 | 1111.1 | 1111.1 | 0.5 | 0.5 | 0.5 | 0.5 | 0.5 | 0.5 | 1 | ADNQH |
| 120.3 | 52.107 | 173.279 | 1111.1 | 1111.1 | 1111.1 | 0.5 | 0.5 | 0.5 | 0.5 | 0.5 | 0.5 | 1 | ARDNEQHLKMFWY |
| 119 | 51.87 | 173.279 | 1111.1 | 1111.1 | 1111.1 | 0.5 | 0.5 | 0.5 | 0.5 | 0.5 | 0.5 | 1 | ARDNEQHLKMFWY |
| 112.7 | 55.95 | 171.6 | 1111.1 | 1111.1 | 1111.1 | 0.5 | 0.5 | 0.5 | 0.5 | 0.5 | 0.5 | 1 | RDNEQHIKMFSTWY |
| ^a^multiplicity (MP), the number of times a signal can be assigned | | | | | | | | | | | | | |

**Supplementary Table 10. SGNN Sets of Resonances for WT-TDP43LC Fibrils.** The following two sets of signals could correspond to S350 to N353 or to S369 to N372.

|  | **NCACX / DARR “i" type.** | | | | |  | **NCOCX “i-1” type** | | | | |  | **CANCO “i and i-1” type** | | |
| --- | --- | --- | --- | --- | --- | --- | --- | --- | --- | --- | --- | --- | --- | --- | --- |
| **Residue** | **^15^N** | **^13^CA** | **^13^CO** | **^13^CB** | **^13^CG** |  | **^15^N** | **^13^CA** | **^13^CO** | **^13^CB** | **^13^CG** |  | **^15^N** | **^13^CA** | **^13^CO** |
| S | 112.609 | 58.9 | 173.021 | 69.088 |  |  | 109.952 | 59.016 | 173.016 | 69.192 |  |  | 112.447 | 58.838 | 172.217 |
| G | 110.189 | 47.293 | 173.092 |  |  |  | 116.87 | 47.114 | 173.496 |  |  |  |  |  |  |
| N | 116.586 | 52.083 | 173.447 | 39.429 | 174.978 |  | 118.861 | 52.139 | 174.487 |  |  |  | 117.06 | 52.256 | 173.359 |
| N | 118.535 | 52.065 | 173.401 | 39.283 | 175.119 |  | 114.922 | 52.194 | 173.032 | 1111.1 | 175.397 |  | 118.542 | 51.654 | 174.328 |

|  | **NCACX / DARR “i" type.** | | | | |  | **NCOCX “i-1” type** | | | | |  | **CANCO “i and i-1” type** | | |
| --- | --- | --- | --- | --- | --- | --- | --- | --- | --- | --- | --- | --- | --- | --- | --- |
| **Residue** | **^15^N** | **^13^CA** | **^13^CO** | **^13^CB** | **^13^CG** |  | **^15^N** | **^13^CA** | **^13^CO** | **^13^CB** | **^13^CG** |  | **^15^N** | **^13^CA** | **^13^CO** |
| S | 122.607 | 57.134 | 174.717 | 65.126 |  |  | 110.415 | 57.267 | 174.973 | 65.263 |  |  |  |  |  |
| G | 110.794 | 43.959 | 171.452 |  |  |  | 118.575 | 44.294 | 170.716 |  |  |  | 110.586 | 43.785 | 174.863 |
| N | 118.535 | 52.065 | 173.401 | 39.283 | 175.119 |  | 114.922 | 52.194 | 173.032 | 1111.1 | 175.397 |  | 118.96 | 51.485 | 170.81 |
| N | 115.011 | 54.77 | 173.167 | 43.079 |  |  | 121.405 | 55.319 | 173.564 |  |  |  | 114.879 | 54.594 | 172.928 |

**Supplementary Table 11. QNQ Sets of Resonances for WT-TDP43LC Fibrils.** The following set of signals could correspond to Q344 to Q346 or Q354 to Q356.

|  | **NCACX / DARR “i" type.** | | | | |  | **NCOCX “i-1” type** | | | | |  | **CANCO “i and i-1” type** | | |
| --- | --- | --- | --- | --- | --- | --- | --- | --- | --- | --- | --- | --- | --- | --- | --- |
| **Residue** | **^15^N** | **^13^CA** | **^13^CO** | **^13^CB** | **^13^CG** |  | **^15^N** | **^13^CA** | **^13^CO** | **^13^CB** | **^13^CG** |  | **^15^N** | **^13^CA** | **^13^CO** |
| Q | 121.046 | 54.159 | 173.856 | 32.943 | 1111.1 |  | 122.205 | 54.007 | 173.77 | 32.887 |  |  |  |  |  |
| N | 122.269 | 52.035 | 172.929 | 41.123 | 174.511 |  | 124.905 | 52.07 | 173.078 | 41.106 | 174.358 |  | 122.159 | 52.172 | 173.684 |
| Q | 124.826 | 53.95 | 173.531 | 32.53 |  |  | 122.205 | 54.007 | 173.77 | 32.887 |  |  | 125.142 | 54.466 | 173.225 |

**Supplementary Table 12. Assignments for P-TDP43LC Fibrils.** See methods for assignment procedure. S/N*^a^* corresponds to the S/N measured by NMRFAM-SPARKY for the corresponding 3D CANCO signal^17^. The * denotes overlapped peak intensity, and the “b” corresponds to residues without a unique assignment from the CANCO spectra specifically, but with signals from the NCACX / NCOCX type spectra.

| **Residue** | **Type** | **N** | **CA** | **CO** | **CB** | **CG** | **CD** | **S/N^a^** |
| --- | --- | --- | --- | --- | --- | --- | --- | --- |
| **287** | **G** |  | 45.327 | 174.5 |  |  |  | b |
| **288** | **G** | 111.551 | 44.392 | 171.297 |  |  |  | 8* |
| **289** | **F** | 121 | 53.1 | 173.8 |  |  |  | b |
| **290** | **G** | 104.653 | 46.761 | 172.303 |  |  |  | 9 |
| **291** | **N** | 116.892 | 51.632 | 173.272 | 38.867 |  |  | b |
| **292** | **S** | 111.381 | 54.521 | 173.894 | 64.236 |  |  | b |
| **293** | **R** | 121.392 | 53.323 | 173.563 | 33.916 | 26.9 | 43.425 | b |
| **294** | **G** | 105.032 | 44.278 | 170.742 |  |  |  | 8* |
| **295** | **G** | 116.364 | 44.545 | 172.48 |  |  |  | b |
| **296** | **G** | 106.61 | 45.699 | 170.267 |  |  |  | 12 |
| **297** | **A** | 121.933 | 51.92 | 174.201 | 20.929 |  |  | 19 |
| **298** | **G** | 106.912 | 44.25 | 171.177 |  |  |  | 10* |
| **308** | **G** | 116.364 | 44.545 | 172.48 |  |  |  | b |
| **309** | **G** | 103.888 | 46.168 | 171.659 |  |  |  | 8 |
| **310** | **G** | 107.09 | 42.873 | 174.398 |  |  |  | 13 |
| **311** | **M** | 110.5 |  |  |  |  |  | b |
| **389** | **S** | 125.396 | 55.979 | 172.019 | 65.024 |  |  |  |
| **390** | **N** | 126.211 | 52.905 | 174.307 | 39.779 |  |  | 10 |
| **391** | **A** | 120.798 | 54.401 | 179.886 | 17.608 |  |  | 15* |

**Supplementary Tables 13. MCASSIGN 2b tables for P-TDP43LC Fibrils.** A) 2D/3D NCACX and CC 50ms DARR table, B) 2D/3D NCOCX, C) 3D CANCO.

A)

| **Observed Chemical Shift (ppm)** | | | | | | **Chemical Shift Uncertainty (ppm)** | | | | | |  |  |
| --- | --- | --- | --- | --- | --- | --- | --- | --- | --- | --- | --- | --- | --- |
| **^15^N** | **^13^CA** | **^13^CO** | **^13^CB** | **^13^CG** | **^13^CD** | **^15^N** | **^13^CA** | **^13^CO** | **^13^CB** | **^13^CG** | **^13^CD** | **MP^a^** | **Residue Type** |
| 121.933 | 51.92 | 174.201 | 20.929 | 1111.1 | 1111.1 | 0.5 | 0.5 | 1 | 0.5 | 0.5 | 0.5 | 1 | A |
| 103.888 | 46.168 | 171.659 | 1111.1 | 1111.1 | 1111.1 | 0.5 | 0.5 | 1 | 0.5 | 0.5 | 0.5 | 1 | G |
| 106.61 | 45.699 | 170.267 | 1111.1 | 1111.1 | 1111.1 | 0.5 | 0.5 | 0.8 | 0.5 | 0.5 | 0.5 | 1 | G |
| 104.653 | 46.761 | 172.303 | 1111.1 | 1111.1 | 1111.1 | 0.5 | 0.5 | 1 | 0.5 | 0.5 | 0.5 | 1 | G |
| 107.09 | 42.873 | 174.398 | 1111.1 | 1111.1 | 1111.1 | 0.5 | 0.5 | 1 | 0.5 | 0.5 | 0.5 | 1 | G |
| 124.863 | 51.533 | 173.824 | 33.681 | 1111.1 | 177.058 | 0.5 | 0.5 | 1 | 0.5 | 0.5 | 0.5 | 1 | EQ |
| 106.912 | 44.25 | 171.177 | 1111.1 | 1111.1 | 1111.1 | 0.5 | 0.5 | 1.5 | 0.5 | 0.5 | 0.5 | 1 | G |
| 105.032 | 44.278 | 170.742 | 1111.1 | 1111.1 | 1111.1 | 0.5 | 0.5 | 1 | 0.5 | 0.5 | 0.5 | 1 | G |
| 110.76 | 54.507 | 173.969 | 63.949 | 1111.1 | 1111.1 | 0.5 | 0.5 | 1 | 0.5 | 0.5 | 0.5 | 1 | S |
| 111.381 | 54.521 | 173.894 | 64.236 | 1111.1 | 1111.1 | 0.5 | 0.5 | 1 | 0.5 | 0.5 | 0.5 | 1 | S |
| 120.798 | 54.401 | 179.886 | 17.608 | 1111.1 | 1111.1 | 0.5 | 0.5 | 1 | 0.5 | 0.5 | 0.5 | 1 | A |
| 113.413 | 47.106 | 174.004 | 1111.1 | 1111.1 | 1111.1 | 0.5 | 0.5 | 1 | 0.5 | 0.5 | 0.5 | 1 | G |
| 113.004 | 48.202 | 171.972 | 1111.1 | 1111.1 | 1111.1 | 0.5 | 0.5 | 1 | 0.5 | 0.5 | 0.5 | 1 | G |
| 111.551 | 44.392 | 171.297 | 1111.1 | 1111.1 | 1111.1 | 0.5 | 0.5 | 1 | 0.5 | 0.5 | 0.5 | 1 | G |
| 113.553 | 51.83 | 173.451 | 41.374 | 1111.1 | 1111.1 | 0.5 | 0.5 | 1 | 0.5 | 0.5 | 0.5 | 1 | DNLFY |
| 113.638 | 56.445 | 173.908 | 64.953 | 1111.1 | 1111.1 | 0.5 | 0.5 | 1 | 0.5 | 0.5 | 0.5 | 1 | S |
| 116.364 | 44.545 | 172.48 | 1111.1 | 1111.1 | 1111.1 | 0.5 | 0.5 | 2 | 0.5 | 0.5 | 0.5 | 1 | G |
| 114.687 | 52.347 | 174.411 | 42.075 | 1111.1 | 1111.1 | 0.5 | 0.5 | 2 | 0.5 | 0.5 | 0.5 | 1 | DNLFY |
| 115.659 | 56.283 | 172.954 | 65.402 | 1111.1 | 1111.1 | 0.5 | 0.5 | 2 | 0.5 | 0.5 | 0.5 | 1 | S |
| 116.916 | 54.721 | 173.673 | 34.046 | 1111.1 | 1111.1 | 0.5 | 0.5 | 1 | 0.5 | 0.5 | 0.5 | 1 | RNEQHKMFWY |
| 116.965 | 51.666 | 174.329 | 39.976 | 1111.1 | 1111.1 | 0.5 | 0.5 | 1.5 | 0.5 | 0.5 | 0.5 | 1 | DNLMFY |
| 116.892 | 51.632 | 173.272 | 38.867 | 1111.1 | 1111.1 | 0.5 | 0.5 | 1 | 0.5 | 0.5 | 0.5 | 1 | DNLKMFY |
| 119.888 | 53.718 | 173.574 | 39.222 | 1111.1 | 1111.1 | 0.5 | 0.5 | 1 | 0.5 | 0.5 | 0.5 | 1 | DNLMFY |
| 128.858 | 54.421 | 173.101 | 34.129 | 1111.1 | 1111.1 | 0.5 | 0.5 | 1 | 0.5 | 0.5 | 0.5 | 1 | RNEQHKMFWY |
| 128.862 | 54.357 | 173.101 | 33.053 | 1111.1 | 1111.1 | 0.5 | 0.5 | 1 | 0.5 | 0.5 | 0.5 | 1 | REQHKMWY |
| 125.886 | 50.189 | 174.26 | 21.663 | 1111.1 | 1111.1 | 0.5 | 0.5 | 1 | 0.5 | 0.5 | 0.5 | 1 | A |
| 125.602 | 50.059 | 174.59 | 23.102 | 1111.1 | 1111.1 | 0.5 | 0.5 | 1 | 0.5 | 0.5 | 0.5 | 1 | A |
| 121.616 | 52.989 | 173.526 | 37.198 | 1111.1 | 1111.1 | 0.5 | 0.5 | 1 | 0.5 | 0.5 | 0.5 | 1 | DNHLKMFY |
| 119.429 | 56.118 | 171.715 | 65.515 | 1111.1 | 1111.1 | 0.5 | 0.5 | 1 | 0.5 | 0.5 | 0.5 | 1 | S |
| 116.294 | 53.379 | 173.861 | 33.336 | 1111.1 | 1111.1 | 0.5 | 0.5 | 1 | 0.5 | 0.5 | 0.5 | 1 | REQHKMWY |
| 117.376 | 55.947 | 172.632 | 64.593 | 1111.1 | 1111.1 | 0.5 | 0.5 | 2 | 0.5 | 0.5 | 0.5 | 1 | S |
| 109.215 | 47.259 | 173.784 | 1111.1 | 1111.1 | 1111.1 | 0.5 | 0.5 | 1 | 0.5 | 0.5 | 0.5 | 1 | G |
| 117.931 | 43.301 | 171 | 1111.1 | 1111.1 | 1111.1 | 0.5 | 0.5 | 1 | 0.5 | 0.5 | 0.5 | 1 | G |
| 108.347 | 44.434 | 171.827 | 1111.1 | 1111.1 | 1111.1 | 0.5 | 0.5 | 2 | 0.5 | 0.5 | 0.5 | 1 | G |
| 113.12 | 56.119 | 174.781 | 65.725 | 1111.1 | 1111.1 | 0.5 | 0.5 | 1.5 | 0.5 | 0.5 | 0.5 | 1 | S |
| 118.221 | 54.781 | 173.452 | 66.178 | 1111.1 | 1111.1 | 0.5 | 0.5 | 1.5 | 0.5 | 0.5 | 0.5 | 1 | S |
| 114.519 | 58.744 | 172.206 | 68.612 | 1111.1 | 1111.1 | 0.5 | 0.5 | 1 | 0.5 | 0.5 | 0.5 | 1 | S |
| 120.718 | 56.03 | 172.621 | 64.758 | 1111.1 | 1111.1 | 0.5 | 0.5 | 1 | 0.5 | 0.5 | 0.5 | 1 | S |
| 121.075 | 55.694 | 172.574 | 63.663 | 1111.1 | 1111.1 | 0.5 | 0.5 | 1 | 0.5 | 0.5 | 0.5 | 1 | S |
| 121.034 | 55.668 | 172.574 | 65.66 | 1111.1 | 1111.1 | 0.5 | 0.5 | 1 | 0.5 | 0.5 | 0.5 | 1 | S |
| 125.396 | 55.979 | 172.019 | 65.024 | 1111.1 | 1111.1 | 0.5 | 0.5 | 1 | 0.5 | 0.5 | 0.5 | 1 | S |
| 129.992 | 49.519 | 173.977 | 21.257 | 1111.1 | 1111.1 | 0.5 | 0.5 | 1 | 0.5 | 0.5 | 0.5 | 1 | A |
| 127.62 | 50.621 | 174.497 | 20.639 | 1111.1 | 1111.1 | 0.5 | 0.5 | 1 | 0.5 | 0.5 | 0.5 | 1 | A |
| 127.669 | 50.274 | 174.471 | 21.648 | 1111.1 | 1111.1 | 0.5 | 0.5 | 1 | 0.5 | 0.5 | 0.5 | 1 | A |
| 128.204 | 50.099 | 174.426 | 22.59 | 1111.1 | 1111.1 | 0.5 | 0.5 | 1 | 0.5 | 0.5 | 0.5 | 1 | A |
| 124.287 | 49.618 | 173.671 | 22.821 | 1111.1 | 1111.1 | 0.5 | 0.5 | 1 | 0.5 | 0.5 | 0.5 | 1 | A |
| 122.441 | 54.335 | 173.604 | 31.642 | 1111.1 | 1111.1 | 0.5 | 0.5 | 1 | 0.5 | 0.5 | 0.5 | 1 | M |
| 121.392 | 53.323 | 173.563 | 33.916 | 26.9 | 43.425 | 0.5 | 0.5 | 1 | 0.5 | 0.5 | 0.5 | 1 | R |
| 121.611 | 52.396 | 173.976 | 45.661 | 1111.1 | 1111.1 | 0.5 | 0.5 | 2 | 0.5 | 0.5 | 0.5 | 1 | DLFY |
| 121.58 | 52.787 | 173.529 | 45.02 | 1111.1 | 1111.1 | 0.5 | 0.5 | 2 | 0.5 | 0.5 | 0.5 | 1 | DLFY |
| 121.643 | 52.607 | 173.529 | 44.158 | 1111.1 | 1111.1 | 0.5 | 0.5 | 2 | 0.5 | 0.5 | 0.5 | 1 | DNLFY |
| 120.841 | 52.897 | 173.449 | 42.745 | 1111.1 | 1111.1 | 0.5 | 0.5 | 2 | 0.5 | 0.5 | 0.5 | 1 | DNLFY |
| 120.319 | 52.194 | 173.452 | 40.83 | 1111.1 | 1111.1 | 0.5 | 0.5 | 2 | 0.5 | 0.5 | 0.5 | 1 | DNLFY |
| 118.977 | 51.84 | 173.281 | 41.552 | 1111.1 | 1111.1 | 0.5 | 0.5 | 1 | 0.5 | 0.5 | 0.5 | 1 | DNLFY |
| 126.211 | 52.905 | 174.307 | 39.779 | 1111.1 | 1111.1 | 0.5 | 0.5 | 1 | 0.5 | 0.5 | 0.5 | 1 | DNLFY |
| 125.945 | 49.983 | 174.426 | 20.158 | 1111.1 | 1111.1 | 0.5 | 0.5 | 1 | 0.5 | 0.5 | 0.5 | 1 | A |
| 124.391 | 50.038 | 174.236 | 21.636 | 1111.1 | 1111.1 | 0.5 | 0.5 | 1 | 0.5 | 0.5 | 0.5 | 1 | A |
| 121.524 | 49.902 | 174.053 | 23.148 | 1111.1 | 1111.1 | 0.5 | 0.5 | 2 | 0.5 | 0.5 | 0.5 | 1 | A |
| 119.01 | 54.247 | 173.37 | 31.837 | 1111.1 | 1111.1 | 0.5 | 0.5 | 1 | 0.5 | 0.5 | 0.5 | 1 | REQHKMW |
| 115.008 | 53.069 | 173.385 | 37.721 | 1111.1 | 1111.1 | 0.5 | 0.5 | 1 | 0.5 | 0.5 | 0.5 | 1 | DNHLKMFY |
| 120.636 | 54.352 | 178.09 | 17.637 | 1111.1 | 1111.1 | 0.5 | 0.5 | 1 | 0.5 | 0.5 | 0.5 | 1 | A |
| 118.768 | 60.089 | 175.264 | 1111.1 | 1111.1 | 1111.1 | 0.5 | 0.5 | 1 | 0.5 | 0.5 | 0.5 | 1 | REQHILKMFSTWY |
| 104.019 | 44.237 | 171.692 | 1111.1 | 1111.1 | 1111.1 | 0.5 | 0.5 | 2 | 0.5 | 0.5 | 0.5 | 1 | G |
| 121.236 | 50.286 | 175.334 | 22.063 | 1111.1 | 1111.1 | 0.5 | 0.5 | 1 | 0.5 | 0.5 | 0.5 | 1 | A |
| 121.935 | 50.522 | 174.816 | 18.546 | 1111.1 | 1111.1 | 0.5 | 0.5 | 1 | 0.5 | 0.5 | 0.5 | 1 | A |
| ^a^multiplicity (MP), the number of times a signal can be assigned | | | | | | | | | | | | | |

B)

| **Observed Chemical Shift (ppm)** | | | | | | **Chemical Shift Uncertainty (ppm)** | | | | | |  |  |
| --- | --- | --- | --- | --- | --- | --- | --- | --- | --- | --- | --- | --- | --- |
| **^15^N** | **^13^CA** | **^13^CO** | **^13^CB** | **^13^CG** | **^13^CD** | **^15^N** | **^13^CA** | **^13^CO** | **^13^CB** | **^13^CG** | **^13^CD** | **MP^a^** | **Residue Type** |
| 106.328 | 51.953 | 174.103 | 20.882 | 1111.1 | 1111.1 | 0.5 | 0.5 | 0.5 | 0.5 | 0.5 | 0.5 | 1 | A |
| 121.967 | 45.767 | 170.24 | 1111.1 | 1111.1 | 1111.1 | 0.5 | 0.5 | 0.5 | 0.5 | 0.5 | 0.5 | 1 | G |
| 112.091 | 53.019 | 174.71 | 39.582 | 1111.1 | 1111.1 | 0.5 | 0.5 | 0.5 | 0.5 | 0.5 | 0.5 | 1 | DNLMFY |
| 112.037 | 44.302 | 171.255 | 1111.1 | 1111.1 | 1111.1 | 0.5 | 0.5 | 0.5 | 0.5 | 0.5 | 0.5 | 1 | G |
| 128.583 | 55.35 | 172.396 | 64.853 | 1111.1 | 1111.1 | 0.5 | 0.5 | 0.5 | 0.5 | 0.5 | 0.5 | 1 | S |
| 112.92 | 56.124 | 171.975 | 65.799 | 1111.1 | 1111.1 | 0.5 | 0.5 | 0.5 | 0.5 | 0.5 | 0.5 | 1 | S |
| 121.352 | 54.401 | 173.717 | 64.022 | 1111.1 | 1111.1 | 0.5 | 0.5 | 0.5 | 0.5 | 0.5 | 0.5 | 1 | S |
| 117.027 | 46.944 | 172.36 | 1111.1 | 1111.1 | 1111.1 | 0.5 | 0.5 | 0.5 | 0.5 | 0.5 | 0.5 | 1 | G |
| 119.654 | 46.939 | 174.016 | 1111.1 | 1111.1 | 1111.1 | 0.5 | 0.5 | 0.5 | 0.5 | 0.5 | 0.5 | 1 | G |
| 118.197 | 54.116 | 173.696 | 33.836 | 1111.1 | 1111.1 | 0.5 | 0.5 | 0.5 | 0.5 | 0.5 | 0.5 | 1 | RNEQHKMFWY |
| 120.904 | 54.294 | 179.72 | 17.547 | 1111.1 | 1111.1 | 0.5 | 0.5 | 0.5 | 0.5 | 0.5 | 0.5 | 1 | A |
| 126.329 | 56.161 | 171.703 | 65.652 | 1111.1 | 1111.1 | 0.5 | 0.5 | 0.5 | 0.5 | 0.5 | 0.5 | 1 | S |
| 126.654 | 56.153 | 171.743 | 65.364 | 1111.1 | 1111.1 | 0.5 | 0.5 | 0.5 | 0.5 | 0.5 | 0.5 | 1 | S |
| 107.244 | 45.99 | 171.636 | 1111.1 | 1111.1 | 1111.1 | 0.5 | 0.5 | 0.5 | 0.5 | 0.5 | 0.5 | 1 | G |
| 106.972 | 53.015 | 172.246 | 34.194 | 1111.1 | 1111.1 | 0.5 | 0.5 | 0.5 | 0.5 | 0.5 | 0.5 | 1 | RNEQHKMFWY |
| 124.274 | 50.517 | 174.244 | 22.303 | 1111.1 | 1111.1 | 0.5 | 0.5 | 0.5 | 0.5 | 0.5 | 0.5 | 1 | A |
| 106.814 | 43.999 | 171.968 | 1111.1 | 1111.1 | 1111.1 | 0.5 | 0.5 | 0.5 | 0.5 | 0.5 | 0.5 | 1 | G |
| 106.338 | 44.63 | 173.853 | 1111.1 | 1111.1 | 1111.1 | 0.5 | 0.5 | 0.5 | 0.5 | 0.5 | 0.5 | 1 | G |
| 104.392 | 53.096 | 173.834 | 1111.1 | 1111.1 | 1111.1 | 0.5 | 0.5 | 0.5 | 0.5 | 0.5 | 0.5 | 1 | ARDNEQHLKMFSWY |
| 104.002 | 44.786 | 173.108 | 1111.1 | 1111.1 | 1111.1 | 0.5 | 0.5 | 0.5 | 0.5 | 0.5 | 0.5 | 1 | G |
| 109.158 | 54.478 | 174.315 | 1111.1 | 1111.1 | 1111.1 | 0.5 | 0.5 | 0.5 | 0.5 | 0.5 | 0.5 | 1 | ARDNEQHLKMFSWY |
| 113.398 | 54.77 | 174.929 | 42.193 | 1111.1 | 1111.1 | 0.5 | 0.5 | 0.5 | 0.5 | 0.5 | 0.5 | 1 | DNLFY |
| 114.953 | 54.469 | 173.424 | 44.284 | 1111.1 | 1111.1 | 0.5 | 0.5 | 0.5 | 0.5 | 0.5 | 0.5 | 1 | DNLFY |
| 114.655 | 52.507 | 173.294 | 38.728 | 1111.1 | 1111.1 | 0.5 | 0.5 | 0.5 | 0.5 | 0.5 | 0.5 | 1 | DNLKMFY |
| 114.782 | 52.43 | 173.339 | 41.04 | 1111.1 | 1111.1 | 0.5 | 0.5 | 0.5 | 0.5 | 0.5 | 0.5 | 1 | DNLMFY |
| 121.478 | 52.963 | 173.865 | 1111.1 | 1111.1 | 1111.1 | 0.5 | 0.5 | 0.5 | 0.5 | 0.5 | 0.5 | 1 | ARDNEQHLKMFSWY |
| 122.6 | 56.031 | 171.966 | 65.179 | 1111.1 | 1111.1 | 0.5 | 0.5 | 0.5 | 0.5 | 0.5 | 0.5 | 1 | S |
| 121.1 | 56.23 | 173.072 | 1111.1 | 1111.1 | 1111.1 | 0.5 | 0.5 | 0.5 | 0.5 | 0.5 | 0.5 | 1 | RDNEQHLKMFSTWY |
| 123.489 | 52.755 | 173.68 | 1111.1 | 1111.1 | 1111.1 | 0.5 | 0.5 | 0.5 | 0.5 | 0.5 | 0.5 | 1 | ARDNEQHLKMFSWY |
| 126.822 | 50.221 | 174.197 | 22.081 | 1111.1 | 1111.1 | 0.5 | 0.5 | 0.5 | 0.5 | 0.5 | 0.5 | 1 | A |
| 120.683 | 50.285 | 174.777 | 23.227 | 1111.1 | 1111.1 | 0.5 | 0.5 | 0.5 | 0.5 | 0.5 | 0.5 | 1 | A |
| 121.103 | 44.488 | 171.746 | 1111.1 | 1111.1 | 1111.1 | 0.5 | 0.5 | 0.5 | 0.5 | 0.5 | 0.5 | 1 | G |
| 123.779 | 44.613 | 171.124 | 1111.1 | 1111.1 | 1111.1 | 0.5 | 0.5 | 0.5 | 0.5 | 0.5 | 0.5 | 1 | G |
| 116.709 | 44.127 | 170.72 | 1111.1 | 1111.1 | 1111.1 | 0.5 | 0.5 | 0.5 | 0.5 | 0.5 | 0.5 | 1 | G |
| 111.594 | 45.327 | 174.202 | 1111.1 | 1111.1 | 1111.1 | 0.5 | 0.5 | 0.5 | 0.5 | 0.5 | 0.5 | 1 | G |
| 107.796 | 44.361 | 170.952 | 1111.1 | 1111.1 | 1111.1 | 0.5 | 0.5 | 0.5 | 0.5 | 0.5 | 0.5 | 1 | G |
| 119.51 | 51.639 | 173.506 | 32.919 | 1111.1 | 1111.1 | 0.5 | 0.5 | 0.5 | 0.5 | 0.5 | 0.5 | 1 | REQHKMWY |
| 128.431 | 53.267 | 172.277 | 1111.1 | 1111.1 | 1111.1 | 0.5 | 0.5 | 0.5 | 0.5 | 0.5 | 0.5 | 1 | ARDNEQHLKMFSWY |
| 111.482 | 52.099 | 173.052 | 38.546 | 1111.1 | 1111.1 | 0.5 | 0.5 | 0.5 | 0.5 | 0.5 | 0.5 | 1 | DNHLKMFY |
| 110.471 | 42.763 | 174.386 | 1111.1 | 1111.1 | 1111.1 | 0.5 | 0.5 | 0.5 | 0.5 | 0.5 | 0.5 | 1 | G |
| 106.362 | 54.601 | 176.906 | 32.126 | 1111.1 | 1111.1 | 0.5 | 0.5 | 0.5 | 2.5 | 0.5 | 0.5 | 1 | REQHKMW |
| 104.417 | 56.373 | 175.073 | 64.121 | 1111.1 | 1111.1 | 0.5 | 0.5 | 0.5 | 0.5 | 0.5 | 0.5 | 1 | S |
| 124.312 | 52.85 | 173.944 | 33.73 | 1111.1 | 1111.1 | 0.5 | 0.5 | 0.5 | 0.5 | 0.5 | 0.5 | 1 | RNEQHKMWY |
| 116.286 | 54.175 | 173.452 | 41.292 | 1111.1 | 1111.1 | 0.5 | 0.5 | 0.5 | 0.5 | 0.5 | 0.5 | 1 | DNLFY |
| 116.883 | 51.821 | 174.422 | 1111.1 | 1111.1 | 1111.1 | 0.5 | 0.5 | 0.5 | 0.5 | 0.5 | 0.5 | 1 | ARDNEQHLKMFWY |
| 119.832 | 54.417 | 179.682 | 17.563 | 1111.1 | 1111.1 | 0.5 | 0.5 | 0.5 | 0.5 | 0.5 | 0.5 | 1 | A |
| 126.1 | 52.852 | 171.14 | 44.586 | 1111.1 | 1111.1 | 0.5 | 0.5 | 0.5 | 0.5 | 0.5 | 0.5 | 1 | DLFY |
| 119.364 | 55.619 | 171.318 | 65.39 | 1111.1 | 1111.1 | 0.5 | 0.5 | 0.5 | 0.5 | 0.5 | 0.5 | 1 | S |
| ^a^multiplicity (MP), the number of times a signal can be assigned | | | | | | | | | | | | | |

C)

| **Observed Chemical Shift (ppm)** | | | | | | **Chemical Shift Uncertainty (ppm)** | | | | | |  |  |
| --- | --- | --- | --- | --- | --- | --- | --- | --- | --- | --- | --- | --- | --- |
| **^15^N** | **^13^CA** | **^13^CO** | **^13^CB** | **^13^CG** | **^13^CD** | **^15^N** | **^13^CA** | **^13^CO** | **^13^CB** | **^13^CG** | **^13^CD** | **MP^a^** | **Residue Type** |
| 121.939 | 51.778 | 170.124 | 1111.1 | 1111.1 | 1111.1 | 0.5 | 0.5 | 0.5 | 0.5 | 0.5 | 0.5 | 1 | ARDNEQHLKMFWY |
| 103.724 | 45.977 | 172.847 | 1111.1 | 1111.1 | 1111.1 | 0.5 | 0.5 | 0.5 | 0.5 | 0.5 | 0.5 | 1 | G |
| 106.6 | 45.312 | 173.755 | 1111.1 | 1111.1 | 1111.1 | 0.5 | 0.5 | 0.5 | 0.5 | 0.5 | 0.5 | 1 | G |
| 104.549 | 46.692 | 174.003 | 1111.1 | 1111.1 | 1111.1 | 0.5 | 0.5 | 0.5 | 0.5 | 0.5 | 0.5 | 1 | G |
| 106.9 | 42.577 | 172.188 | 1111.1 | 1111.1 | 1111.1 | 0.5 | 0.5 | 0.5 | 0.5 | 0.5 | 0.5 | 1 | G |
| 124.8 | 54.134 | 173.906 | 1111.1 | 1111.1 | 1111.1 | 0.5 | 0.5 | 0.5 | 0.5 | 0.5 | 0.5 | 1 | ARDNEQHILKMFSWY |
| 106.958 | 44.252 | 174.264 | 1111.1 | 1111.1 | 1111.1 | 0.5 | 0.5 | 0.5 | 0.5 | 0.5 | 0.5 | 1 | G |
| 105.085 | 44.367 | 173.972 | 1111.1 | 1111.1 | 1111.1 | 0.5 | 0.5 | 0.5 | 0.5 | 0.5 | 0.5 | 1 | G |
| 110.983 | 54.313 | 172.996 | 1111.1 | 1111.1 | 1111.1 | 0.5 | 0.5 | 0.5 | 0.5 | 0.5 | 0.5 | 1 | RDNEQHMFSWY |
| 110.983 | 54.313 | 172.996 | 1111.1 | 1111.1 | 1111.1 | 0.5 | 0.5 | 0.5 | 0.5 | 0.5 | 0.5 | 1 | RDNEQHMFSWY |
| 121.1 | 54.451 | 173.5 | 1111.1 | 1111.1 | 1111.1 | 0.5 | 0.5 | 0.5 | 0.5 | 0.5 | 0.5 | 1 | ARDNEQHILKMFSWY |
| 113.148 | 46.829 | 175.205 | 1111.1 | 1111.1 | 1111.1 | 0.5 | 0.5 | 0.5 | 0.5 | 0.5 | 0.5 | 1 | G |
| 111.507 | 44.398 | 174.871 | 1111.1 | 1111.1 | 1111.1 | 0.5 | 0.5 | 0.5 | 0.5 | 0.5 | 0.5 | 1 | G |
| 113.7 | 52.311 | 173.639 | 1111.1 | 1111.1 | 1111.1 | 0.5 | 0.5 | 0.5 | 0.5 | 0.5 | 0.5 | 1 | RDNEQHLKMFSWY |
| 113.7 | 56.419 | 174.241 | 1111.1 | 1111.1 | 1111.1 | 0.5 | 0.5 | 0.5 | 0.5 | 0.5 | 0.5 | 1 | RDNEQHILKMFSTWY |
| 116.2 | 44.504 | 173.459 | 1111.1 | 1111.1 | 1111.1 | 0.5 | 0.5 | 0.5 | 0.5 | 0.5 | 0.5 | 1 | G |
| 114.776 | 52.651 | 173.379 | 1111.1 | 1111.1 | 1111.1 | 0.5 | 0.5 | 0.5 | 0.5 | 0.5 | 0.5 | 1 | ARDNEQHLKMFSWY |
| 115.6 | 56.174 | 174.225 | 1111.1 | 1111.1 | 1111.1 | 0.5 | 0.5 | 0.5 | 0.5 | 0.5 | 0.5 | 1 | RDNEQHILKMFSTWY |
| 117.206 | 54.588 | 173.976 | 1111.1 | 1111.1 | 1111.1 | 0.5 | 0.5 | 0.5 | 0.5 | 0.5 | 0.5 | 1 | ARDNEQHILKMFSWY |
| 116.938 | 51.738 | 172.862 | 1111.1 | 1111.1 | 1111.1 | 0.5 | 0.5 | 0.5 | 0.5 | 0.5 | 0.5 | 1 | ARDNEQHLKMFWY |
| 116.938 | 51.738 | 172.862 | 1111.1 | 1111.1 | 1111.1 | 0.5 | 0.5 | 0.5 | 0.5 | 0.5 | 0.5 | 1 | ARDNEQHLKMFWY |
| 128.732 | 54.148 | 172.62 | 1111.1 | 1111.1 | 1111.1 | 0.5 | 0.5 | 0.5 | 0.5 | 0.5 | 0.5 | 1 | ARDNEQHILKMFWY |
| 128.732 | 54.119 | 172.648 | 1111.1 | 1111.1 | 1111.1 | 0.5 | 0.5 | 0.5 | 0.5 | 0.5 | 0.5 | 1 | ARDNEQHILKMFWY |
| 126.487 | 50.16 | 173.761 | 1111.1 | 1111.1 | 1111.1 | 0.5 | 0.5 | 0.5 | 0.5 | 0.5 | 0.5 | 1 | ADNEQHLM |
| 126.487 | 50.16 | 173.761 | 1111.1 | 1111.1 | 1111.1 | 0.5 | 0.5 | 0.5 | 0.5 | 0.5 | 0.5 | 1 | ADNEQHLM |
| 121.6 | 52.943 | 173.846 | 1111.1 | 1111.1 | 1111.1 | 0.5 | 0.5 | 0.5 | 0.5 | 0.5 | 0.5 | 1 | ARDNEQHLKMFSWY |
| 116.3 | 53.374 | 172.998 | 1111.1 | 1111.1 | 1111.1 | 0.5 | 0.5 | 0.5 | 0.5 | 0.5 | 0.5 | 1 | ARDNEQHLKMFSWY |
| 117.4 | 56.032 | 174.277 | 1111.1 | 1111.1 | 1111.1 | 0.5 | 0.5 | 0.5 | 0.5 | 0.5 | 0.5 | 1 | RDNEQHILKMFSTWY |
| 118.808 | 43.132 | 172.859 | 1111.1 | 1111.1 | 1111.1 | 0.5 | 0.5 | 0.5 | 0.5 | 0.5 | 0.5 | 1 | G |
| 112.7 | 44.646 | 174.189 | 1111.1 | 1111.1 | 1111.1 | 0.5 | 0.5 | 0.5 | 0.5 | 0.5 | 0.5 | 1 | G |
| 108.4 | 44.317 | 174.068 | 1111.1 | 1111.1 | 1111.1 | 0.5 | 0.5 | 0.5 | 0.5 | 0.5 | 0.5 | 1 | G |
| 125.013 | 56.092 | 174.432 | 1111.1 | 1111.1 | 1111.1 | 0.5 | 0.5 | 0.5 | 0.5 | 0.5 | 0.5 | 1 | RDNEQHILKMFSTWY |
| 128.9 | 50.419 | 173.73 | 1111.1 | 1111.1 | 1111.1 | 0.5 | 0.5 | 0.5 | 0.5 | 0.5 | 0.5 | 1 | ARDNEQHLKM |
| 129.709 | 49.609 | 171.167 | 1111.1 | 1111.1 | 1111.1 | 0.5 | 0.5 | 0.5 | 0.5 | 0.5 | 0.5 | 1 | ADNQH |
| 120.3 | 52.107 | 173.279 | 1111.1 | 1111.1 | 1111.1 | 0.5 | 0.5 | 0.5 | 0.5 | 0.5 | 0.5 | 1 | ARDNEQHLKMFWY |
| 119 | 51.87 | 173.279 | 1111.1 | 1111.1 | 1111.1 | 0.5 | 0.5 | 0.5 | 0.5 | 0.5 | 0.5 | 1 | ARDNEQHLKMFWY |
| 126.2 | 52.948 | 171.78 | 1111.1 | 1111.1 | 1111.1 | 0.5 | 0.5 | 0.5 | 0.5 | 0.5 | 0.5 | 1 | ARDNEQHLKMFSWY |
| 112.7 | 55.95 | 171.6 | 1111.1 | 1111.1 | 1111.1 | 0.5 | 0.5 | 0.5 | 0.5 | 0.5 | 0.5 | 1 | RDNEQHIKMFSTWY |
| 117.752 | 60.4 | 172.629 | 1111.1 | 1111.1 | 1111.1 | 0.5 | 0.5 | 0.5 | 0.5 | 0.5 | 0.5 | 1 | REQHILKMFSTWY |
| ^a^multiplicity (MP), the number of times a signal can be assigned | | | | | | | | | | | | | |

**Supplementary Table 14. Experimental Parameters for Solid State NMR Measurements, WT-TDP43LC Fibrils**

| **Spectrum** | **Acquisition Parameters^a,b^** | **Processing Parameters^c^** |
| --- | --- | --- |
| **Sample 1 WT-TDP43LC Fibrils** | | |
| 2D ^13^C-^13^C CP-DARR | ns= 64;$\tau$_aq_ = 10.24 ms ; $\nu$_1H-carr_= 4.464 ppm; $\nu$_13C-carr_= 99.713 ppm; $\nu$_1H-CP_ = 63 kHz; $\nu$_13C-CP_= 50 kHz; $\nu$_1Hdec_= 83.3 kHz; $\nu$_DARR_= 13 kHz; $\tau_{1H-\pi/2}$=3 $\mu$s; $\tau_{13C-\pi/2}$=4 $\mu$s; $\tau_{\mathrm{CP}}$= 1.0 ms; $\tau_{\mathrm{DARR}}$ = 50 ms; $\tau_{1H-dec}$= 6 $\mu$s; $\Delta t_{1}$ = 12 $\mu$s; $\tau_{t1}$ = 5.7 ms | GLB_t1_ = 100 Hz  GLB_t2_ = 100 Hz  POLY -auto -ord 4 |
| 2D CP-NCACX | ns= 320; $\tau$_aq_ = 10.24 ms ; $\nu$_1H-carr_= 4.464 ppm; $\nu$_13C-carr_= 55.554 ppm; $\nu$_15N-carr_= 116.332 ppm; $\nu$_1H-CP_ = 48.7 kHz; $\nu$_15N-CP_ = 35.7 kHz; $\nu$_15N-SCP_ = 35.1 kHz; $\nu$_13CA-SCP_= 22.1 kHz; $\nu$_1Hdec_= 83.3 kHz; $\nu$_DARR_= 13 kHz; $\tau_{1H-\pi/2}$= 3 $\mu$s; $\tau_{13C-\pi/2}$= 4 $\mu$s; $\tau_{15N-\pi/2}$=6 $\mu$s; $\tau_{\mathrm{CP}}$= 0.75 ms; $\tau_{\mathrm{SCP}}$= 4 ms; $\tau_{\mathrm{DARR}}$ = 50 ms; $\tau_{1H-dec}$= 6 $\mu$s; $\Delta t_{1}$ = 108 $\mu$s; $\tau_{t1}$ = 10.044 ms | GLB_t2_ = 60 Hz |
| 2D CP-NCOCX | ns=320; $\tau$_aq_ = 10.24 ms ; $\nu$_1H-carr_= 4.464 ppm; $\nu$_13C-carr_= 182.296 ppm; $\nu$_15N-carr_= 120.623 ppm; $\nu$_1H-CP_ = 48.7 kHz; $\nu$_15N-CP_ = 35.7 kHz; $\nu$_15N-SCP_ = 35.1 kHz; $\nu$_13CO-SCP_=48.1 kHz; $\nu$_1Hdec_= 83.3 kHz; $\nu$_DARR_= 13 kHz; $\tau_{1H-\pi/2}$= 3 $\mu$s; $\tau_{13C-\pi/2}$= 4 $\mu$s; $\tau_{15N-\pi/2}$=6 $\mu$s; $\tau_{\mathrm{CP}}$= 0.75 ms; $\tau_{\mathrm{SCP}}$= 4 ms; $\tau_{\mathrm{DARR}}$ = 50 ms; $\tau_{1H-dec}$= 6 $\mu$s; $\Delta t_{1}$ = 108 $\mu$s; $\tau_{t1}$ = 10.044 ms | GLB_t1_ = 100 Hz  GLB_t2_ = 60 Hz |
| 3D CP-NCACX | ns= 48; $\tau$_aq_ = 10.24 ms ; $\nu$_1H-carr_= 4.706 ppm; $\nu$_13C-carr_= 55.797 ppm; $\nu$_15N-carr_= 116.574 ppm; $\nu$_1H-CP_ = 48.7 kHz; $\nu$_15N-CP_ = 35.7 kHz; $\nu$_15N-SCP_ = 35.1 kHz; $\nu$_13CA-SCP_= 22.1 kHz; $\nu$_1Hdec_= 83.3 kHz; $\nu$_DARR_= 13 kHz; $\tau_{1H-\pi/2}$= 3 $\mu$s; $\tau_{13C-\pi/2}$= 4 $\mu$s; $\tau_{15N-\pi/2}$=6 $\mu$s; $\tau_{\mathrm{CP}}$= 0.75 ms; $\tau_{\mathrm{SCP}}$= 4 ms; $\tau_{\mathrm{DARR}}$ = 50 ms; $\tau_{1H-dec}$= 6 $\mu$s; $\Delta t_{1}$ = 180 $\mu$s; $\tau_{t1}$ = 7.56 ms; $\Delta t_{2}$ = 132.0 $\mu$s; $\tau_{t2}$ = 3.3 ms | POLY -ord 4 |
| 3D CP-NCOCX | ns= 56; $\tau$_aq_ = 10.24 ms ; $\nu$_1H-carr_= 4.949 ppm; $\nu$_13C-carr_= 174.719 ppm; $\nu$_15N-carr_= 120.501 ppm; $\nu$_1H-CP_ = 48.7 kHz; $\nu$_15N-CP_ = 35.7 kHz; $\nu$_15N-SCP_ = 37.7 kHz; $\nu$_13CO-SCP_=50.7 kHz; $\nu$_1Hdec_= 83.3 kHz; $\nu$_DARR_= 13 kHz; $\tau_{1H-\pi/2}$= 3 $\mu$s; $\tau_{13C-\pi/2}$= 4 $\mu$s; $\tau_{15N-\pi/2}$=6 $\mu$s; $\tau_{\mathrm{CP}}$= 1.5 ms; $\tau_{\mathrm{SCP}}$= 4 ms; $\tau_{\mathrm{DARR}}$ = 50 ms; $\tau_{1H-dec}$= 6 $\mu$s; $\Delta t_{1}$ = 180 $\mu$s; $\tau_{t1}$ = 5.04 ms; $\Delta t_{2}$ = 156.0 $\mu$s; $\tau_{t2}$ = 3.9 ms | GLB_t1_ = 100 Hz  GLB_t2_ = 60 Hz  GLB_t3_ = 100 Hz  POLY -ord 4 |
| 3D CP-CANCO | ns= 16; $\tau$_aq_ = 10.24 ms ; $\nu$_1H-carr_= 4.023 ppm; $\nu$_13C-carr_= 56.363 ppm; $\nu$_15N-carr_= 121.431ppm; $\nu$_1H-CP_ = 63 kHz; $\nu$_13C-CP_ = 50 kHz; $\nu$_15N-SCP_ = 35.1 kHz; $\nu$_13CA-SCP_= 22.1 kHz; $\nu_{13CO-SCP-eff}$= 48.1 kHz; $\nu$_1Hdec_= 83.3 kHz; $\tau_{1H-\pi/2}$= 3 $\mu$s; $\tau_{13C-\pi/2}$= 4 $\mu$s; $\tau_{15N-\pi/2}$=6 $\mu$s; $\tau_{CP1H-13C}$= 0.5 ms; $\tau_{SCP13C-15N}$= 4 ms; $\tau_{SCP15N-13C}$= 4 ms;$\tau_{1H-dec}$= 6 $\mu$s; $\Delta t_{2}$ = 180 $\mu$s; $\tau_{t2}$ = 7.56 ms; $\Delta t_{1}$ = 60 $\mu$s; $\tau_{t1}$ = 3.36 ms | GLB_t1_ = 150 Hz  GLB_t2_ = 150 Hz  GLB_t3_ = 60 Hz |
| **Sample 2 WT-TDP43LC Fibrils** | | |
| 2D ^13^C-^13^C CP-DARR | ns= 64 $\tau$_aq_ = 10.24 ms ; $\nu$_1H-carr_= 3.736 ppm; $\nu$_13C-carr_= 98.985 ppm; $\nu$_1H-CP_ = 63 kHz; $\nu$_13C-CP_= 50 kHz; $\nu$_1Hdec_= 83.3 kHz; $\nu$_DARR_= 13 kHz; $\tau_{1H-\pi/2}$=3 $\mu$s; $\tau_{13C-\pi/2}$=4 $\mu$s; $\tau_{\mathrm{CP}}$= 1.0 ms; $\tau_{\mathrm{DARR}}$ = 50 ms; $\tau_{1H-dec}$= 6 $\mu$s; $\Delta t_{1}$ = 12 $\mu$s; $\tau_{t1}$ = 5.7 ms | GLB_t1_ = 100 Hz  GLB_t2_ = 100 Hz |
| 2D ^13^C-^13^C rINEPT-TOBSY | ns = 64; $\tau$_aq_ = 20.48 ms; $\nu$_1H-carr_= 3.978 ppm; $\nu$_13C-carr_= 41.153 ppm; $\tau_{J/2}$ = 1 ms; $\Delta t_{1}$ = 24 $\mu$s, $\tau_{t1}$ = 5.28 ms; $\tau_{13C-TOBSY}$=39 kHz; $\tau_{mix-TOBSY}$=7.38 ms; uses $POST-{C9}_{6}^{1}$ | GLB_t1_ = 100 Hz  GLB_t2_ = 100 Hz |
| 2D ^1^H-^13^C rINEPT | ns = 32; $\tau$_aq_ = 30.72 ms; $\nu$_1H-carr_= 3.979 ppm; $\nu$_13C-carr_= 95.527 ppm; $\tau_{J/2}$ = 1 ms; $\Delta t_{1}$ = 75 $\mu$s, $\tau_{t1}$ = 11.25 ms; $\tau_{1H-\pi/2}=3\mu s$; $\tau_{13C-\pi/2}=4\mu s$ | GLB_t1_ = 50 Hz  GLB_t2_ = 20 Hz |
| NCACX ^15^N T2 | ns = 128, $\tau_{\mathrm{aq}}$ = 7.68 ms; $\nu$_1H-carr_= 4.221 ppm; $\nu$_13C-carr_= 55.312 ppm;$\nu$_15N-carr_= 120.38 ppm; $\tau_{1H-\pi/2}=3\mu s$; $\tau_{13C-\pi/2}=4\mu s$, $\nu$_15N-CP_ = 35.7 kHz; $\nu$_15N-SCP_ = 35.1 kHz; $\nu$_13C-SCP_ = 22.1 kHz $\nu$_1H-CP_ = 48.7 kHz; $\tau_{CP1H-15N}=0.75ms$, $\tau_{CP15N-13C}=4ms$; $\nu$_1Hdec_= 83.3 kHz;  vdlist: 0.0769, 0.4614, 0.9997, 1.4611, 1.923, 2.384, 2.845, 3.23, 3.614, 3.999, 4.46, 5.075, 5.537, 6.075, 6.613, 7.306 ms | EM = 20Hz (Topspin 4.1.4 processing) |
| cp90T1rho1H | ns = 128, $\tau_{\mathrm{aq}}$ = 7.68 ms, $\nu$_1H-carr_= 4.221 ppm; $\nu$_13C-carr_= 99.471 ppm; $\tau_{1H-\pi/2}=3\mu s$; $\tau_{13C-\pi/2}=4\mu s$, $\nu$_13C-CP_ = 50 kHz, $\nu$_1H-CP_ = 63 kHz; $\tau_{CP1H-13C}=1ms$; $\nu$_1Hdec_= 83.3 kHz;  vplist: 0.01, 0.5, 1, 2, 3, 4, 5, 6, 7, 8, 9, 10 ms | EM = 20Hz (Topspin 4.1.4 processing) |

^a^All of the ^1^H-X CP transfers use a 20% ramp on the ^1^H channel. All ^15^N-^13^C Specific-CP transfers used a 5% ramp on the ^15^N channel.

^b^ns is the number of scans averaged; $\tau_{aq}$ is the total acquisition time in the directly detected dimension; $\nu$_X-carr_ is the center frequency; $\nu$_X-CP_ is the nutation frequency of the applied power during the CP step; $\nu$_X-SCP_ is the nutation frequency from the applied power during the specific CP step; $\nu$_1Hdec_ is the decoupling pulse for ^1^H; $\nu$_DARR_ is the ^13^C – ^13^C mixing pulse power on ^1^H; $\tau_{X-\pi/2}$ is the nuclei specified 90° pulse length; $\tau_{13C-TOBSY}$is the ^13^C TOBSY pulse length; $\tau_{CP}$ is the contact time for CP; $\tau_{SCP}$ is the contact time for specific CP; $\tau_{DARR}$ is the ^13^C-^13^C DARR mixing time; $\tau_{mix-TOBSY}$ is the total TOBSY mixing time; $\tau_{1H-dec}$ is the SPINAL64 $\pi$ pulse length; $\Delta t_{x}$is the increment in the specified indirect time dimension; $\tau_{tx}$ is the total acquisition time for the specified dimension, $\tau_{J/2}$is the ½ echo delay; $\nu_{13CO-SCP-eff}$ is the effective SCP pulse to ^13^C carbonyl carbon from ^15^N.

^c^ Spectra are processed in NMRPipe. Gaussian line broadening was applied according to the parameters specified. Zero filling was applied twice before applying each Fourier transform.

**Supplementary Table 15. Experimental Parameters for Solid State NMR Measurements, P-TDP43LC Fibrils**

| **Spectrum** | **Acquisition Parameters^a,b^** | **Processing Parameters^c^** |
| --- | --- | --- |
| **Sample 1 P-TDP43LC Fibrils** | | |
| 2D ^13^C-^13^C CP-DARR | ns= 64;$\tau$_aq_ = 10.24 ms ; $\nu$_1H-carr_= 3.493 ppm; $\nu$_13C-carr_= 98.743 ppm; $\nu$_1H-CP_ = 63 kHz; $\nu$_13C-CP_= 50 kHz; $\nu$_1Hdec_= 83.3 kHz; $\nu$_DARR_= 13 kHz; $\tau_{1H-\pi/2}$=3 $\mu$s; $\tau_{13C-\pi/2}$=4 $\mu$s; $\tau_{\mathrm{CP}}$= 1.0 ms; $\tau_{\mathrm{DARR}}$ = 50 ms; $\tau_{1H-dec}$= 6 $\mu$s; $\Delta t_{1}$ = 12 $\mu$s; $\tau_{t1}$ = 5.7 ms | GLB_t1_ = 100 Hz  GLB_t2_ = 100 Hz  POLY -auto -ord 4 |
| NCACX ^15^N T2 | ns = 128, $\tau_{\mathrm{aq}}$ = 7.68 ms; $\nu$_1H-carr_= 3.493 ppm; $\nu$_13C-carr_= 54.584 ppm;$\nu$_15N-carr_= 119.652 ppm; $\tau_{1H-\pi/2}=3\mu s$; $\tau_{13C-\pi/2}=4\mu s$, $\nu$_15N-CP_ = 35.7 kHz; $\nu$_15N-SCP_ = 35.1 kHz; $\nu$_13C-SCP_ = 22.1 kHz $\nu$_1H-CP_ = 48.7 kHz; $\tau_{CP1H-15N}=0.75ms$, $\tau_{CP15N-13C}=4ms$; $\nu$_1Hdec_= 83.3 kHz;  vdlist: 0.0769, 0.4614, 0.9997, 1.4611, 1.923, 2.384, 2.845, 3.23, 3.614, 3.999, 4.46, 5.075, 5.537, 6.075, 6.613, 7.306 ms | EM = 20Hz (Topspin 4.1.4 processing) |
| cp90T1rho1H | ns = 128, $\tau_{\mathrm{aq}}$ = 7.68 ms, $\nu$_1H-carr_= 3.493 ppm; $\nu$_13C-carr_= 98.742 ppm; $\tau_{1H-\pi/2}=3\mu s$; $\tau_{13C-\pi/2}=4\mu s$, $\nu$_13C-CP_ = 50 kHz, $\nu$_1H-CP_ = 63 kHz; $\tau_{CP1H-13C}=1ms$; $\nu$_1Hdec_= 83.3 kHz;  vplist: 0.01, 0.5, 1, 2, 3, 4, 5, 6, 7, 8, 9, 10 ms | EM = 20Hz (Topspin 4.1.4 processing) |
| **Sample 2 P-TDP43LC Fibrils** | | |
| 2D ^13^C-^13^C CP-DARR | ns= 48;$\tau$_aq_ = 15.36 ms ; $\nu$_1H-carr_= 3.372 ppm; $\nu$_13C-carr_= 98.621 ppm; $\nu$_1H-CP_ = 63 kHz; $\nu$_13C-CP_= 50 kHz; $\nu$_1Hdec_= 83.3 kHz; $\nu$_DARR_= 13 kHz; $\tau_{1H-\pi/2}$=3 $\mu$s; $\tau_{13C-\pi/2}$=4 $\mu$s; $\tau_{\mathrm{CP}}$= 1.0 ms; $\tau_{\mathrm{DARR}}$ = 50 ms; $\tau_{1H-dec}$= 6 $\mu$s; $\Delta t_{1}$ = 12 $\mu$s; $\tau_{t1}$ = 5.7 ms | GLB_t1_ = 100 Hz  GLB_t2_ = 100 Hz |
| NCACX ^15^N T2 | ns = 128, $\tau_{\mathrm{aq}}$ = 7.68 ms; $\nu$_1H-carr_= 3.372 ppm; $\nu$_13C-carr_= 54.463 ppm;$\nu$_15N-carr_= 119.530 ppm; $\tau_{1H-\pi/2}=3\mu s$; $\tau_{13C-\pi/2}=4\mu s$, $\nu$_15N-CP_ = 35.7 kHz; $\nu$_15N-SCP_ = 35.1 kHz; $\nu$_13C-SCP_ = 22.1 kHz; $\nu$_1H-CP_ = 48.7 kHz; $\tau_{CP1H-15N}=0.75ms$, $\tau_{CP15N-13C}=4ms$; $\nu$_1Hdec_= 83.3 kHz;  vdlist: 0.0769, 0.4614, 0.9997, 1.4611, 1.923, 2.384, 2.845, 3.23, 3.614, 3.999, 4.46, 5.075, 5.537, 6.075, 6.613, 7.306 ms | EM = 20Hz (Topspin 4.1.4 processing) |
| cp90 T1rho ^1^H | ns = 128, $\tau_{\mathrm{aq}}$ = 7.68 ms, $\nu$_1H-carr_= 3.372 ppm; $\nu$_13C-carr_= 98.621 ppm; $\tau_{1H-\pi/2}=3\mu s$; $\tau_{13C-\pi/2}=4\mu s$, $\nu$_13C-CP_ = 50 kHz, $\nu$_1H-CP_ = 63 kHz; $\tau_{CP1H-13C}=1ms$; $\nu$_1Hdec_= 83.3 kHz;  vplist: 0.01, 0.5, 1, 2, 3, 4, 5, 6, 7, 8, 9, 10 ms | EM = 20Hz (Topspin 4.1.4 processing) |
| **Sample 3 P-TDP43LC Fibrils** | | |
| 2D ^13^C-^13^C CP-DARR | ns= 48;$\tau$_aq_ = 15.36 ms ; $\nu$_1H-carr_= 3.615ppm; $\nu$_13C-carr_= 98.864 ppm; $\nu$_1H-CP_ = 63 kHz; $\nu$_13C-CP_= 50 kHz; $\nu$_1Hdec_= 83.3 kHz; $\nu$_DARR_= 13 kHz; $\tau_{1H-\pi/2}$=3 $\mu$s; $\tau_{13C-\pi/2}$=4 $\mu$s; $\tau_{\mathrm{CP}}$= 1.0 ms; $\tau_{\mathrm{DARR}}$ = 50 ms; $\tau_{1H-dec}$= 6 $\mu$s; $\Delta t_{1}$ = 12 $\mu$s; $\tau_{t1}$ = 5.7 ms | GLB_t1_ = 100 Hz  GLB_t2_ = 100 Hz |
| 2D CP-NCACX | ns= 256; $\tau$_aq_ = 10.28 ms; $\nu$_1H-carr_= 3.615 ppm; $\nu$_13C-carr_= 54.705 ppm; $\nu$_15N-carr_= 115.407 ppm; $\nu$_1H-CP_ = 48.7 kHz; $\nu$_15N-CP_ = 35.7 kHz; $\nu$_15N-SCP_ = 35.1 kHz; $\nu$_13CA-SCP_= 22.1 kHz; $\nu$_1Hdec_= 83.3 kHz; $\nu$_DARR_= 13 kHz; $\tau_{1H-\pi/2}$= 3 $\mu$s; $\tau_{13C-\pi/2}$= 4 $\mu$s; $\tau_{15N-\pi/2}$=6 $\mu$s; $\tau_{\mathrm{CP}}$= 0.75 ms; $\tau_{\mathrm{SCP}}$= 4 ms; $\tau_{\mathrm{DARR}}$ = 50 ms; $\tau_{1H-dec}$= 6 $\mu$s; $\Delta t_{1}$ = 108 $\mu$s; $\tau_{t1}$ = 10.476 ms | GLB_t1_ = 100 Hz  GLB_t2_ = 100 Hz |
| 2D CP-NCOCX | ns=256; $\tau$_aq_ = 10.28 ms ; $\nu$_1H-carr_= 3.615ppm; $\nu$_13C-carr_= 181.446 ppm; $\nu$_15N-carr_= 119.773 ppm; $\nu$_1H-CP_ = 48.7 kHz; $\nu$_15N-CP_ = 35.7 kHz; $\nu$_15N-SCP_ = 37.7 kHz; $\nu$_13CO-SCP_=50.7 kHz; $\nu$_1Hdec_= 83.3 kHz; $\nu$_DARR_= 13 kHz; $\tau_{1H-\pi/2}$= 3 $\mu$s; $\tau_{13C-\pi/2}$= 4 $\mu$s; $\tau_{15N-\pi/2}$=6 $\mu$s; $\tau_{\mathrm{CP}}$= 0.75 ms; $\tau_{\mathrm{SCP}}$= 4 ms; $\tau_{\mathrm{DARR}}$ = 50 ms; $\tau_{1H-dec}$= 6 $\mu$s; $\Delta t_{1}$ = 108 $\mu$s; $\tau_{t1}$ = 10.0476 ms | GLB_t1_ = 100 Hz  GLB_t2_ = 100 Hz |
| 3D CP-NCACX | ns= 32; $\tau$_aq_ = 15.36 ms; $\nu$_1H-carr_= 3.615 ppm; $\nu$_13C-carr_= 54.705 ppm; $\nu$_15N-carr_= 115.482 ppm; $\nu$_1H-CP_ = 48.7 kHz; $\nu$_15N-CP_ = 35.7 kHz; $\nu$_15N-SCP_ = 35.1 kHz; $\nu$_13CA-SCP_= 22.1 kHz; $\nu$_1Hdec_= 83.3 kHz; $\nu$_DARR_= 13 kHz; $\tau_{1H-\pi/2}$= 3 $\mu$s; $\tau_{13C-\pi/2}$= 4 $\mu$s; $\tau_{15N-\pi/2}$=6 $\mu$s; $\tau_{\mathrm{CP}}$= 0.75 ms; $\tau_{\mathrm{SCP}}$= 4 ms; $\tau_{\mathrm{DARR}}$ = 50 ms; $\tau_{1H-dec}$= 6 $\mu$s; $\Delta t_{1}$ = 180 $\mu$s; $\tau_{t1}$ = 7.56 ms; $\Delta t_{2}$ = 132.0 $\mu$s; $\tau_{t2}$ = 3.3 ms | GLB_t1_ = 100 Hz  GLB_t2_ = 60 Hz  GLB_t3_ = 80 Hz |
| 3D CP-NCOCX | ns= 40; $\tau$_aq_ = 15.36 ms; $\nu$_1H-carr_= 3.615 ppm; $\nu$_13C-carr_= 181.446 ppm; $\nu$_15N-carr_= 119.773 ppm; $\nu$_1H-CP_ = 48.7 kHz; $\nu$_15N-CP_ = 35.7 kHz; $\nu$_15N-SCP_ = 37.7 kHz; $\nu$_13CO-SCP_=50.7 kHz; $\nu$_1Hdec_= 83.3 kHz; $\nu$_DARR_= 13 kHz; $\tau_{1H-\pi/2}$= 3 $\mu$s; $\tau_{13C-\pi/2}$= 4 $\mu$s; $\tau_{15N-\pi/2}$=6 $\mu$s; $\tau_{\mathrm{CP}}$= 0.75 ms; $\tau_{\mathrm{SCP}}$= 4 ms; $\tau_{\mathrm{DARR}}$ = 50 ms; $\tau_{1H-dec}$= 6 $\mu$s; $\Delta t_{1}$ = 180 $\mu$s; $\tau_{t1}$ = 7.56 ms; $\Delta t_{2}$ = 132 $\mu$s; $\tau_{t2}$ = 3.3 ms | GLB_t1_ = 100 Hz  GLB_t2_ = 80 Hz  GLB_t3_ = 100 Hz |
| 3D CP-CANCO | ns= 20; $\tau$_aq_ = 10.24 ms ; $\nu$_1H-carr_= 2.365 ppm; $\nu$_13C-carr_= 54.705 ppm; $\nu$_15N-carr_= 115.482 ppm; $\nu$_1H-CP_ = 63 kHz; $\nu$_13C-CP_ = 50 kHz; $\nu$_15N-SCP-CA-N_ = 35.1 kHz; $\nu$_13CA-SCP_= 22.1 kHz; $\nu$_15N-SCP-N-CO_ = 37.7 kHz; $\nu_{13CO-SCP-eff}$= 50.7 kHz; $\nu$_1Hdec_= 83.3 kHz; $\tau_{1H-\pi/2}$= 3 $\mu$s; $\tau_{13C-\pi/2}$= 4 $\mu$s; $\tau_{15N-\pi/2}$=6 $\mu$s; $\tau_{CP1H-13C}$= 0.75 ms; $\tau_{SCP13C-15N}$= 4 ms; $\tau_{SCP15N-13C}$= 4 ms;$\tau_{1H-dec}$= 6 $\mu$s; $\Delta t_{2}$ = 180 $\mu$s; $\tau_{t2}$ = 7.56 ms; $\Delta t_{1}$ = 60 $\mu$s; $\tau_{t1}$ = 3.36 ms | GLB_t1_ = 50 Hz; g3 = 0.1  GLB_t2_ = 100 Hz  GLB_t3_ = 100 Hz |
| 2D ^13^C-^13^C rINEPT-TOBSY, | ns = 32; $\tau$_aq_ = 20.48 ms; $\nu$_1H-carr_= 3.979 ppm; $\nu$_13C-carr_= 41.153 ppm; $\tau_{J/2}$ = 1 ms; $\Delta t_{1}$ = 24 $\mu$s, $\tau_{t1}$ = 5.28 ms; $\tau_{13C-TOBSY}$=39 kHz; $\tau_{mix-TOBSY}$=7.38 ms; uses $POST-{C9}_{6}^{1}$ | GLB_t1_ = 100 Hz  GLB_t2_ = 100 Hz |
| 2D ^1^H-^13^C rINEPT | ns = 32; $\tau$_aq_ = 30.72 ms; $\nu$_1H-carr_= 3.979 ppm; $\nu$_13C-carr_= 95.527 ppm; $\tau_{J/2}$ = 1 ms; $\Delta t_{1}$ = 75 $\mu$s, $\tau_{t1}$ = 11.25 ms; $\tau_{1H-\pi/2}=3\mu s$; $\tau_{13C-\pi/2}=4\mu s$ | GLB_t1_ = 50 Hz  GLB_t2_ = 20 Hz |
| NCACX ^15^N T2 | ns = 128, $\tau_{\mathrm{aq}}$ = 7.68 ms; $\nu$_1H-carr_= 3.615ppm; $\nu$_13C-carr_= 54.705 ppm;$\nu$_15N-carr_= 119.773 ppm; $\tau_{1H-\pi/2}=3\mu s$; $\tau_{13C-\pi/2}=4\mu s$, $\nu$_15N-CP_ = 35.7 kHz; $\nu$_15N-SCP_ = 35.1 kHz; $\nu$_13C-SCP_ = 22.1 kHz $\nu$_1H-CP_ = 48.7 kHz; $\tau_{CP1H-15N}=0.75ms$, $\tau_{CP15N-13C}=4ms$; $\nu$_1Hdec_= 83.3 kHz;  vdlist: 0.0769, 0.4614, 0.9997, 1.4611, 1.923, 2.384, 2.845, 3.23, 3.614, 3.999, 4.46, 5.075, 5.537, 6.075, 6.613, 7.306 ms | EM = 20Hz (Topspin 4.1.4 processing) |
| cp90T1rho1H | ns = 128, $\tau_{\mathrm{aq}}$ = 7.68 ms, $\nu$_1H-carr_= 3.615 ppm; $\nu$_13C-carr_= 98.864 ppm; $\tau_{1H-\pi/2}=3\mu s$; $\tau_{13C-\pi/2}=4\mu s$, $\nu$_13C-CP_ = 50 kHz, $\nu$_1H-CP_ = 63 kHz; $\tau_{CP1H-13C}=1ms$; $\nu$_1Hdec_= 83.3 kHz;  vplist: 0.01, 0.5, 1, 2, 3, 4, 5, 6, 7, 8, 9, 10 ms | EM = 20Hz (Topspin 4.1.4 processing) |

^a^All of the ^1^H-X CP transfers use a 20% ramp on the ^1^H channel. All ^15^N-^13^C Specific-CP transfers used a 5% ramp on the ^15^N channel.

^b^ns is the number of scans averaged; $\tau_{aq}$ is the total acquisition time in the directly detected dimension; $\nu$_X-carr_ is the center frequency; $\nu$_X-CP_ is the nutation frequency of the applied power during the CP step; $\nu$_X-SCP_ is the nutation frequency from the applied power during the specific CP step; $\nu$_1Hdec_ is the decoupling pulse for ^1^H; $\nu$_DARR_ is the ^13^C – ^13^C mixing pulse power on ^1^H; $\tau_{X-\pi/2}$ is the nuclei specified 90° pulse length; $\tau_{13C-TOBSY}$is the ^13^C TOBSY pulse length; $\tau_{CP}$ is the contact time for CP; $\tau_{SCP}$ is the contact time for specific CP; $\tau_{DARR}$ is the ^13^C-^13^C DARR mixing time; $\tau_{mix-TOBSY}$ is the total TOBSY mixing time; $\tau_{1H-dec}$ is the SPINAL64 $\pi$ pulse length; $\Delta t_{x}$is the increment in the specified indirect time dimension; $\tau_{tx}$ is the total acquisition time for the specified dimension, $\tau_{J/2}$is the ½ echo delay; $\nu_{13CO-SCP-eff}$ is the effective SCP pulse to ^13^C carbonyl carbon from ^15^N.

^c^ Spectra are processed in NMRPipe. Gaussian line broadening was applied according to the parameters specified. Zero filling was applied twice before applying each Fourier transform.
